# A taxonomy of multiple stable states in complex ecological communities

**DOI:** 10.1101/2023.08.30.555051

**Authors:** Guim Aguadé-Gorgorió, Jean-François Arnoldi, Matthieu Barbier, Sonia Kéfi

**Affiliations:** ISEM, University of Montpellier, CNRS, IRD, EPHE, Montpellier, France; Centre for Biodiversity Theory and Modelling, Theoretical and Experimental Ecology Station, CNRS and Paul Sabatier University, 09200 Moulis, France; PHIM Plant Health Institute, University of Montpellier, CIRAD, INRAE, Institut Agro, IRD, 34090 Montpellier, France; Santa Fe Institute, 1399 Hyde Park Road, Santa Fe, NM 87501, USA

**Keywords:** Multiple Stable States, Regime shifts, Complex systems, Community ecology

## Abstract

Many natural and man-made systems, from financial markets to ecosystems or the human brain, are built from multiple interconnected units. This complex high-dimensionality hinders our capacity to understand and predict the dynamics, functioning and fragility of these systems. One fragility scenario, particularly relevant to ecological communities of interacting species, concerns so-called regime shifts: abrupt and unexpected transitions from healthy, species-rich communities towards states of degraded ecosystem function and services. The accepted explanation for these shifts is that they arise as abrupt transitions between alternative stable states: multiple stable configurations of a system under the same internal and external conditions. These alternative states are well-understood in low-dimensional systems, but how they upscale with system complexity remains a debated question. In the present work we investigate the emergence of multiple stable states in a number of complex system models. We find that high-dimensional models with random interactions can unfold at least four different regimes of multistability, each emerging under a specific interaction scheme. More importantly, each multistability regime leaves a distinct and quantifiable fingerprint, providing a framework to analyze experimental evidence of abrupt shifts. By bridging previous results and studying multistability regimes, their fingerprints and their correlation with empirical evidence in ecology, our study helps define a common ground to understand and classify multiple stable states in complex systems.

## INTRODUCTION

Natural ecosystems are deteriorating at un-prece-dented rates, with climate change deeply altering their functioning and the services they provide to human societies. There is, therefore, an urgent need to understand how ecosystems react to environmental changes, a task made difficult by the inherent complexity of communities of many interacting species. Of particular importance are ecosystems found to respond abruptly to gradual changes in environmental conditions [1; 2]. Key examples of these so-called *catastrophic shifts* are the abrupt eutrophication of shallow lakes [3; 4], the desertification of arid ecosystems [5; 6], the bleaching of coral reefs [7] or the degradation of tropical forests into treeless landscapes [8]. The occurrence of abrupt transitions suggests that some ecosystems can exist in multiple stable states within a range of environmental conditions. Small perturbations could then induce transitions between those states, leading to large shifts from species-rich communities towards undesired states of deteriorated ecosystem functions and services.

The possibility that dynamical systems can be multistable and thereby undergo abrupt shifts is relevant across a variety of research fields [9]: Sharp shifts have also been reported in financial markets [10], the human gut microbiome [11] or neuronal activity in the brain [12], all of which are characterized by a complex architecture of many interacting elements. Uncovering what mechanisms make these systems multistable is a key step towards predicting their fragility.

The mechanisms generating multistability have been thoroughly described by single- or few-unit dynamical models that do not always capture the key ingredients at play in the complex system they aim at describing [13; 14; 15]. In ecology more specifically, the current understanding of multistability and related *tipping points* largely ignores the role of species diversity and the complexity of their interactions, potentially failing at characterizing species-rich dynamical processes [1; 14; 16; 17]. Why, and how, multiple stable states can emerge in species-rich ecosystems remain largely open questions (Fig. 1, [15]).

**FIG. 1.**
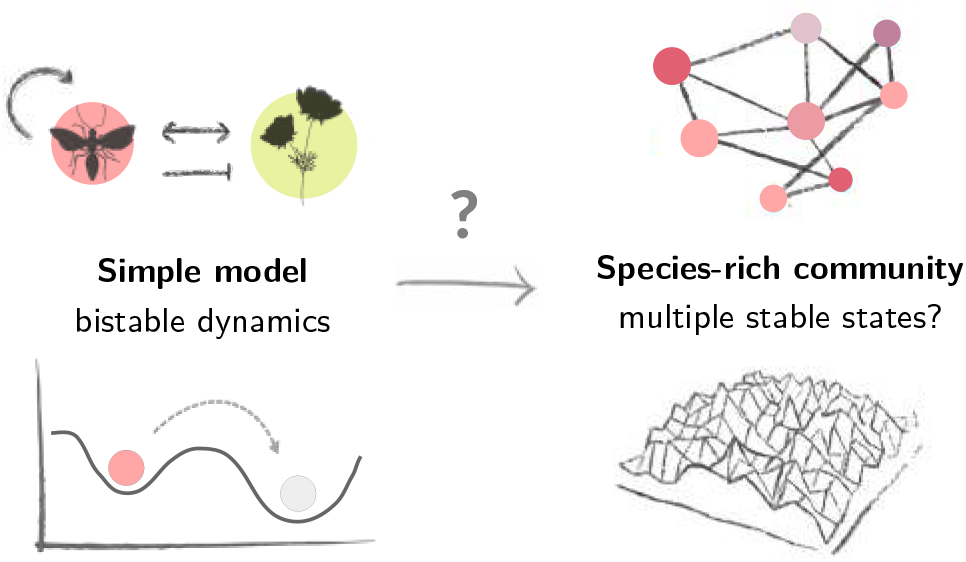
How does low-dimensional bistability scale up to the community level? Our understanding of abrupt transitions in ecosystems mostly relies on simple models that do not take into account the role of species diversity. In the present work, we study the emergence of four different regimes of multistability in species-rich community models and across complex systems.

Recent research has uncovered specific scenarios in which multiple fixed points do emerge in species-rich communities. Theoretical studies focusing on mutualistic systems, *i*.*e*. assuming only positive interactions between all pairs of species in a community, have identified the emergence of system-wide bistability arising from obligate and nonlinear cooperation [18; 19]. Another large body of research is unveiling a very different scenario, where heterogeneous competition can engender vast numbers of similar community states (see *e*.*g*. [20; 21; 22] and Supporting Information (SI) II.F and III.A for an extensive review of the litera-ture). Beyond purely theoretical advances, multistability has also been recently studied in experimental communities of microbial species [23; 24].

How these scenarios are connected, if others exist, and how to detect them empirically remains unclear: we lack a general common framework to describe the emergence of multistability in complex ecosystems. Such a framework should be able to accommodate the various multistability regimes recently described, while at the same time proposing how each could be detected in natural systems.

## METHODS

In this study, we present a theoretical and computational analysis of multistability in species-rich ecological communities, and replicate it across models of other complex biological systems, namely gene regulatory networks, cancer-immune interactions and random neural networks (see SI III). Our aim is to bridge the gap between theory and observations, by characterizing the nature of different multistability scenarios and their relation to driving mechanisms and empirical patterns.

While our results hold across different dynamical models (see SI III), we here analyze a system made up of interacting species, each undergoing bistable population dynamics. Studying a model that harbors species-level bistability has two advantages. First, it allows us to investigate if, and how, local bistability can generate global, *i*.*e*. system-scale, multistability (Fig. 1), or, alternatively, if multistability can emerge without the need for local bistability. Second, and more importantly, it will prove useful to describe how different multistability regimes can emerge from a single model.

In ecology, single-species bistability occurs for example when an Allee Effect limits population growth at low densities [25]. Here we study a community with species subject to an Allee Effect and linked to each other via random interactions. Since the influential work by Robert May [26], modeling species-rich communities with random interaction strengths has provided a unified lens to understand many ecological scenarios: when many species and interaction schemes become intertwined, the details of a community cease to matter and one can ask whether generic patterns emerge (see SI I.C and [26; 27]).

With this in mind, we analyze here mutualistic and competitive interactions, encoded in matrices *A* and *B*, and explore other interaction types in SI II.D. We study the dynamics of *N* species, labeled *i* ( 1, 2, …, *N*, with the abundance *x*_*i*_ of species *i* following

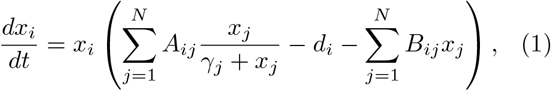

where *A*_*ii*_ is the maximum growth rate of species *i* when isolated and *B*_*ii*_ determines the strength of self-regulation. Mutualism is included by considering that the abundance of species *j* can increase the growth rate of *i*, saturating at high abundances to a maximum *A*_*ij*_ value [28]. The parameter *γ*_*j*_ determines the amount of *j*-individuals necessary to achieve half of the maximum mutualistic contribution to growth rate, as a proxy for how difficult it is for species to cooperate [28]. The matrix elements *B*_*ij*_ *>* 0 determine the strength of negative interactions between species *i* and *j* due to competition for space or resources [29]. Interaction matrices *A* and *B* are built as two independent Erdős-Rényi graphs [26]. Each edge *j* → *i* exists with probability *p* and is sampled from a random distribution, or else *A*_*ij*_ = 0 or *B*_*ij*_ = 0 with probability (1 − *p*) (see Methods and SI I.C.4, II.F.3). Under the assumption of species independence (*A*_*ij*_ = *B*_*ij*_ = 0 for *i* ≠ *j*), species *i* can be either present 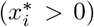 or extinct 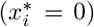, depending on whether the initial condition *x*_*i*_(*t* = 0) falls below or beyond the Allee Effect threshold (see SI II.A). The community can then be at any of 2^*N*^ stable states, encompassing all possible species combinations from community extinction to global coexistence. How does this change when species interact? Can different interaction types generate different multistability scenarios (Fig. 1)?

## RESULTS

The chosen model exhibits a multilayer structure [30], with interaction types segregated into two different matrices, namely *A* for mutualistic interactions and *B* for competition. This feature enables us to investigate the individual effects of each interaction separately. We explore next the multistability scenarios that emerge under relatively homogeneous mutu-alism, competition and their combination, and then describe the effects of increasing the heterogeneity of interactions.

### Global and local stable states in mutualistic communities

Recent research has studied species-rich communities interacting only through cooperation (*i*.*e. B*_*ij*_ = 0 for *i* ≠ *j*) [18]. In line with those studies, and provided interactions are nonlinear, interaction strengths are strongly homogeneous and species connectivity is high (*A*_*ij*_ ≈ ⟨*A*⟩ ∀*i* ≠ *j, p* ≈ 1, see SI II.B, II.F.3), our results predict that communities can become bistable, wherein all species can either go extinct or coexist simultaneously (Fig. 2A right, see SI II.B and [18; 31] for a detailed description). This happens in a parameter range where species do not survive if alone (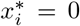 if *A*_*ij*_ ≈ 0, see SI II.A) but cooperation is strong enough. The community (Eq. 1) then collectively reproduces the bistable dynamics of the 1-D system: no species will survive unless the total initial abundance overcomes a predictable threshold (see SI II.B.1), in which case all species survive together (Fig. 2A right). This indicates that *global bistability* can emerge in systems where bistable species cooperate strongly.

**FIG. 2.**
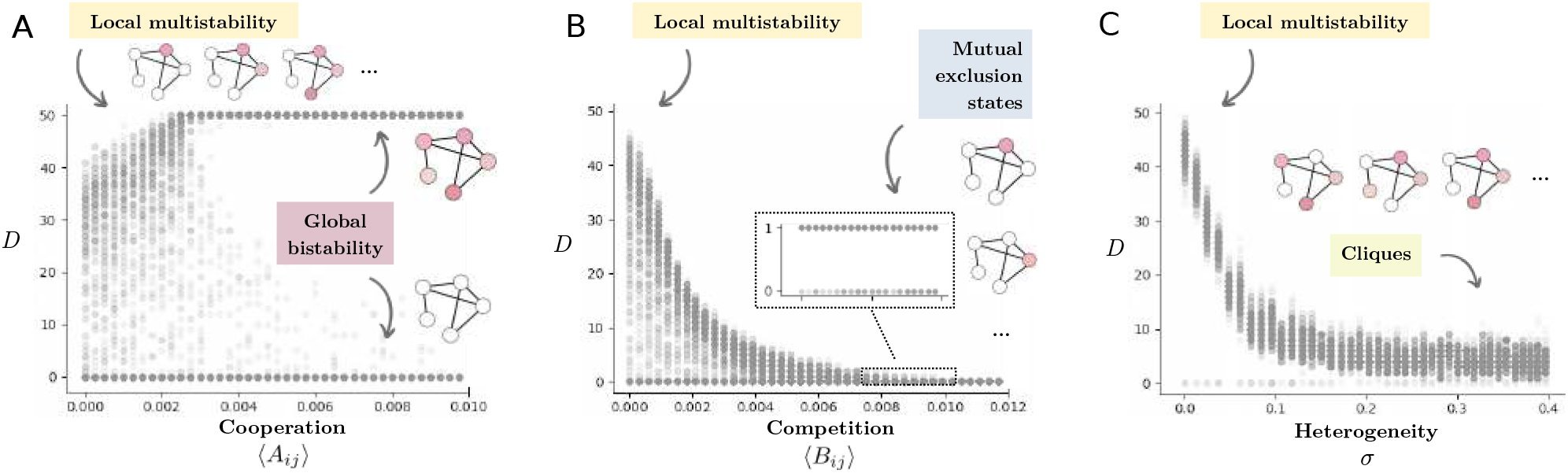
Multiple stable states under increasing mutualism, competition and interaction heterogeneity. We explore how local multistability, resulting from independent bistable species, upscales under increasing mutualism (A), competition (B) and interaction heterogeneity (C). Each gray dot represents the final diversity (*D*, number of surviving species) of a simulation starting from random initial conditions and *N* = 50 species (200 random initial conditions are generated for each value of the x-axis). (A) Under increasing mutualism, the 2*N* states of independent species bistability upscale towards an all-or-nothing bistable regime. (B) Competition reduces the amount of species that can survive in these local states, until we see *N* states, each with only one species surviving. (C) Under heterogeneous interactions, the system can be at many stable states but all with similar diversity. We consider low heterogeneity (*σ* = 0.05) for (A) and (B), and weak interactions (⟨*A*_*ij*_ = *A*_*ii*_*/*10, ⟨*B*_*ij*_ ⟩ = *B*_*ii*_*/*10) for (C).

However, this perfect all-or-nothing regime breaks down if some species can survive without the need for cooperation with others and interactions are relatively weak (see SI II.B.2). In this scenario, many of the 2^*N*^ *local multistability* states, where some species are present and others not, can emerge (Fig. 2A left). This scenario becomes particularly relevant close to the tipping point of the globally bistable system. Here the abrupt extinction of the whole community be-comes blurred by many intermediate states, and transitions between these states are characterized by gradual turnover of only one or few species (see SI II.B.2).

### Local and mutual exclusion states in competitive communities

What type of multiple stable states can emerge under competitive interactions alone (*i*.*e. A*_*ij*_ = 0 for *i* ≠ *j*)? In figure 2B we maintain the above homogeneity constraints on parameters, interactions and species connectivity, and gradually increase the strength of average interspecies competition ⟨*B*_*ij*_⟩. Under very weak competition, we find that the community can be again in many of the possible 2^*N*^ species presence-absence configurations (Fig. 2B left). As competition increases, the number of species that can coexist decreases as expected [32; 33]. However, local bistabilities can still generate a multiplicity of stable states: because stronger competitors can be present or absent individually, and therefore exclude (or not) weaker species, the community can still be in many local states: their multiplicity will be bounded by competition strength (Fig. 2B middle and SI II.C).

Under very strong competition, we recover the well-known scenario of *mutual exclusion* [2]: there exist *N* states with only one surviving competitor each (Fig. 2B box). Beyond that, under even higher interspecies competition and provided local Allee Effects are in place, only the global extinction state will prevail: even the best competitor will be pushed below its Allee Effect threshold due to initial competitive interactions (Fig. 2B bottom right, SI II.C.3 and [34]).

### Different multistability regimes under mixed interactions

Ecological communities are hardly purely mutualistic nor competitive, and they usually encompass a mixture of positive and negative interspecies effects [35] that is likely to vary widely across ecosystems [36]. To explore the consequences of this, we study the emergence of multistability across a wide range of interaction type combinations. Under the same homogeneity and connectivity constraints, we compute in figure 3A the number of observed stable states Ω for different ⟨*A*⟩, ⟨*B*⟩ conditions after a given number of simulations *s* with random initial conditions (see Methods and SI I.D).

**FIG. 3.**
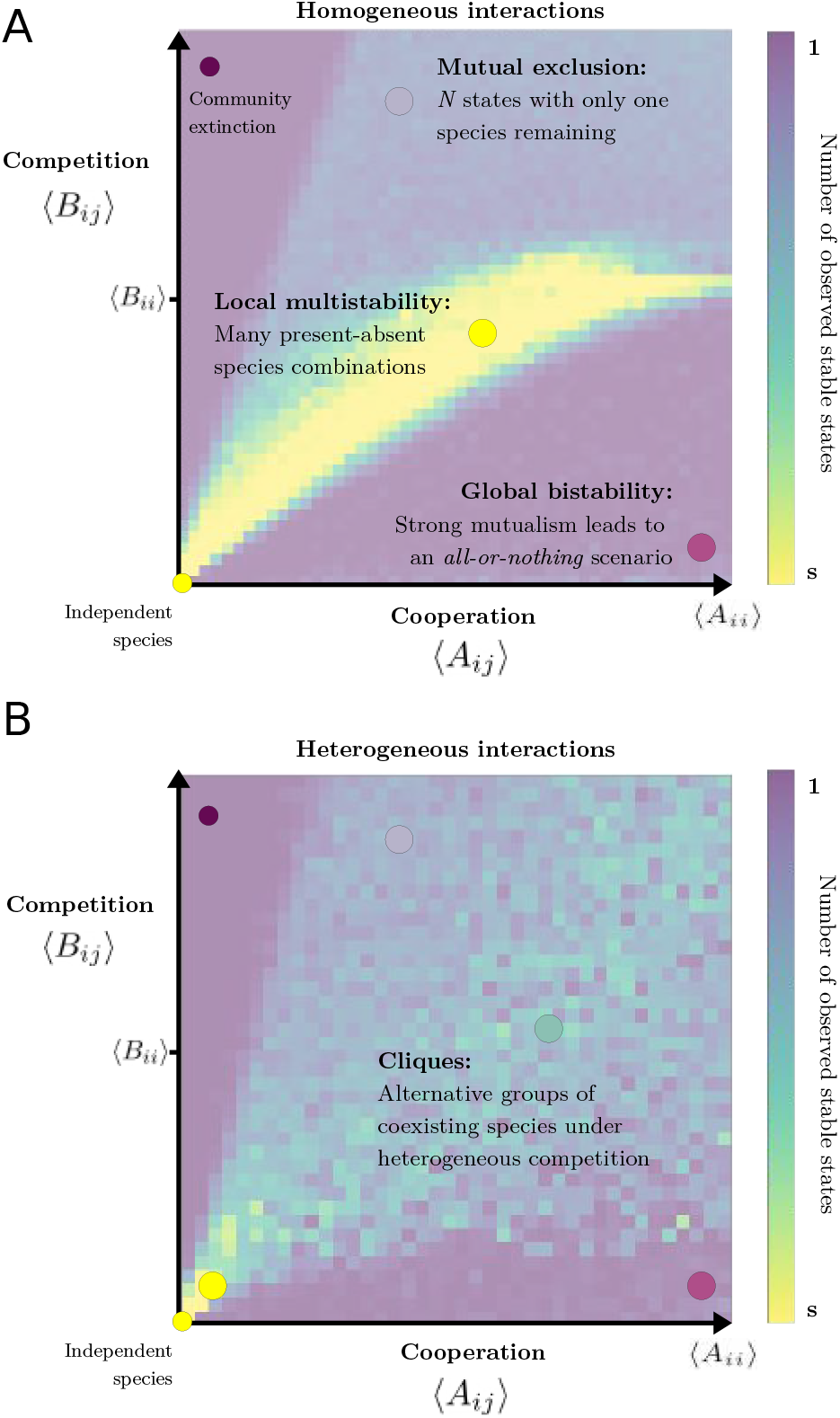
Four multistability regimes in ecological communities: Number of stable states (Ω, color scale) observed after *s* simulations with random initial conditions, for homogeneous (A) and heterogeneous (B) interaction strengths. Highly-mutualistic systems show a global bistability pattern (Ω ∼ 2) emerging from single-species bistabilities. At the other extreme, strong compe-tition generates mutual exclusion states as well as community extinction (Ω ∼ *N* + 1). In between these, single-species bistabilities can generate a high multiplicity of lo-cal states (Ω *>> N*). Heterogeneous interactions (B) generate a fourth multistability regime characterized by many states with similar biomass and diversity. The map spans (max(⟨*B*_*ij*_ ⟩) = 2⟨*B*_*ii*_⟩ and max(⟨*A*_*ij*_ ⟩) = 5max(⟨*B*_*ij*_ ⟩).

We find that global bistability arises when the community is strongly dominated by mutualism, so that the system exists in two stable states (Fig. 3 purple). At the other extreme, strong competition can generate *N* mutual exclusion states (Fig. 3 blue), each with only one surviving species, and community ex-tinction at even stronger ⟨*B*_*ij*_⟩ (Fig. 3 top left). Under a mixture of interaction types, we find that the two domains are separated by a local multistability region with many possible community states (Fig. 3 yellow). These states arise when the effects of mutualism and competition are balanced out and local bistabilities emerge, allowing for a large number of species presence-absence combinations (see SI II.E.1). These results indicate that a strongly homogeneous system can already give rise to three different types of multiple stable states, depending on the set of positive and negative interactions at play.

### Emerging cliques under heterogeneous interactions

The previous results are valid under homogeneous interactions, meaning that interaction strengths and connectivity are relatively uniform across species (see Methods and SI I). Ecological communities, however, are known to exhibit heterogeneous interactions [37; 38]. How does heterogeneity alter the presence of multiple stable states? As we increase the standard deviation of interaction strengths, our models capture the emergence of another multistability domain, which takes over a large proportion of the phase space (Fig. 3B). This new regime is characterized by a high number of stable states, all with similar diversity (Fig. 2C and SI II.F, III), as well as a fraction of initial conditions that do not stabilize through time but rather result in persistent abundance fluctuations (see SI II.F.2).

The emergence of multistability under heterogeneous interactions can be explained within the context of intransitive competition [39; 40]: non-uniform competition strengths preclude the establishment of a linear competition hierarchy. This implies that species can beat some competitors and lose to some others, so that a single competition matrix can encode different groups of coexisting species that balance their negative interactions without a single “winner” [41].

The same multistability regime emerges under purely linear but heterogeneous interaction strengths (see SI III, [42]) or under homogeneous strengths but non-uniform network connectivity (0 *< p <* 1, see SI II.F.3). It is in this context of sparse interaction matrices that the term *cliques* was first used to describe these alternative subgroups of coexisting species [22]. Because we do not restrict our analysis to graphs of coexisting and mutually exclusive species (see SI II.F), we employ the term here in its broader meaning borrowed from the social sciences: a small, often exclusive group of interacting individuals, each a metaphor for a different species.

Consistent with our findings, a growing body of research studying species-rich communities under heterogeneous interactions has described the emergence of a highly complex set of multiple fixed points (see *e*.*g*. [20; 42] and SI II.F, III.A for a detailed literature review). The number and stability of these states, however, varies depending on the specific assumptions of each ecological model (see [22; 43; 44] and SI II.F.2). For instance, random asymmetric interactions (*A*_*ij*_ ≠ *A*_*ji*_) between a very large number of species (*N* → ∞) can lead to unstable cliques, associated to chaotic fluctuations in the presence of noise or species migration [44; 45; 46]. In this context, unstable cliques become more relevant as models of temporal community variability [46; 47; 48] rather than stable community states separated by abrupt transitions (see II.F.2).

However, we find that a fraction of cliques can achieve stability when communities harbor a moderate number of species, such as those found in experimental or controlled environments (*N* = 10^1^ ∼ 10^2^, see e.g [48; 49]). This means that multiple stable states can still exist under intransitive competition and asymmetric interactions (see SI II.F.2). Other ecological models have also identified mechanisms through which cliques can maintain stability amidst heterogeneous competition [21; 22; 50]. Furthermore, we do not observe a sharp and uniform heterogeneity threshold beyond which only cliques emerge in systems of moderate size [26; 42]. Instead, different multistability regimes can manifest depending on the strengths of species interactions (Fig. 3B, [46] and SI II.F.1).

Overall, different ecological assumptions can result in variations in the stability or quantity of cliques that an intransitive system can support (see SI II.F and III.A). Nevertheless, the fundamental nature of the cliques regime remains consistent: heterogeneous interactions can lead to the emergence of numerous stable states with similar species diversity, along with persistent species fluctuations.

### Fingerprints of each multistability regime

At least four different multistability regimes can emerge in a single species-rich community model. Each is driven by specific interaction schemes and can encompass a very different number of stable states (Fig. 2, 3). But what makes the community states in these regimes really different, and how could we identify them in data? When confronted with experimental evidence of complex multistability, can our framework help in discerning what type of states and underlying mechanisms are at play?

Here, we show that key state properties are interrelated differently for each of the four multistability regimes. Among many potentially relevant metrics (see II.G), we present three that can be effectively tested in experimental setups. These follow from simple questions: *does the studied system stabilize into fixed states?, how many of these stable states do we observe?* and *what are the properties of these states?*

In the following we measure (1) the fraction of initial conditions that reach a fixed stable state *S* without fluctuations in species abundances [47; 48]; (2) the number of observed stable states Ω and (3) the diversity of surviving species in these states *D*, a key property linked to community function [51]. Considering these three observables, we generate a collection of systems with random interaction strengths and heterogeneity (see Methods). For each system, we generate trajectories from random initial conditions and plot the final state in the space of possible (*S*, Ω, *D*) metrics (Fig. 4A).

**FIG. 4.**
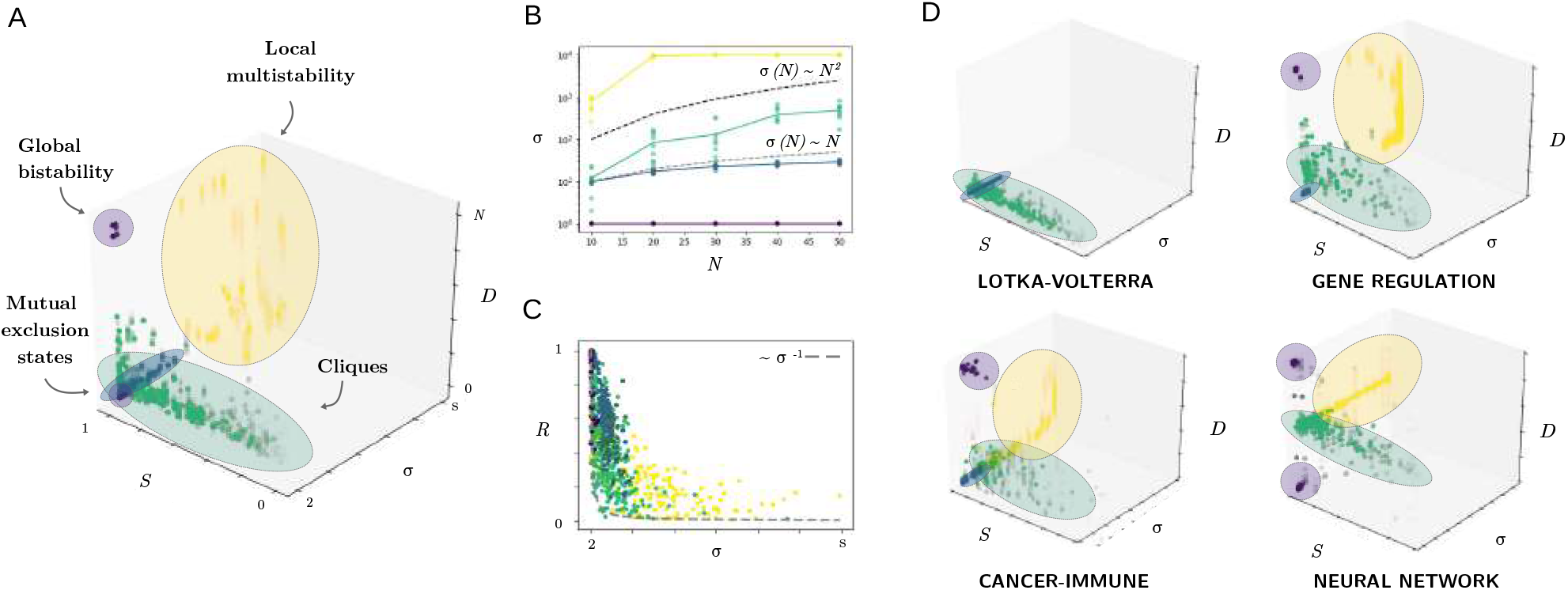
Each multistability type leaves a distinct empirical fingerprint. Here we analyze the signatures that each multistability type would leave in different experimental tests. Each dot represents the final state of a simulation: we generate 400 communities with random ⟨*A*⟩, ⟨*B*⟩, *σ* and *N* = 50. For each community, we run *s* = 300 simulations starting from random initial conditions (see Methods and SI I). (A) Proportion of initial conditions that reach a stable state *S*, number of different stable states observed Ω and diversity *D* of surviving species in each state. Each multistability family seats within a well-defined cluster, with colors chosen consistent with those of figure 3. (B) Scaling of the number of observed states Ω with species number *N*, with *N* ranging from 10 to 50. Here, we generate up to *s* = 104 initial conditions to ensure we explore a large number of possible stable states Ω. 200 communities are generated with parameters chosen to fall inside each domain of figure 3. (C) We also test stable states against random perturbations of variable magnitude (Δ*x*_*i*_, see Methods and SI) and count the fraction of times the same state is recovered as a proxy for *basin stability*. Our tests show a lower resilience bound at *R* ∼ Ω^−1^ (dashed line): the basin stability of stable communities rapidly decays when they are surrounded by a multiplicity of other states. (D) Qualitatively similar fingerprints are found for different complex systems: Generalized Lotka-Volterra communities (GLV), gene regulatory networks (GRN), cancer-immune interactions and random neural networks (see SI III).

We also study two additional tests prevalent across experimental studies: how state properties scale with system size [52] and the effects of pulse perturbations altering species abundances [53]. For the first, we measure how Ω and *D* scale with the number of species *N* (Fig. 4B and SI II.G.1-2). For the second, once each system reaches a stable state, we perturb it with random abundance changes Δ*x*_*i*_, and measure the fraction of communities that recover their original composition, *R*, as a proxy for resilience [54] and basin stability [55] (Fig. 4C and SI II.G.3).

These simple tests show that each multistability regime is likely to leave an experimental fingerprint of its own (Fig. 4). Across simulations and models, global bistability scenarios have an expectedly simple signature (Fig. 4, purple): they are characterized by two stable states involving all or no surviving species, a pattern that remains stable across system sizes (Ω(*N*) = 2 and *D*(*N*) = 0 or *N*). Moreover, the *D* = *N* state is very resilient to perturbations, as cooperation between many species drastically reduces the basin of attraction of ***x*** = 0 (see SI II.B.2).

In contrast, local multistability (Fig. 4, yellow) comprises many possible stable states, some of which might take very long to stabilize, resulting in apparent fluctuations (see SI II.E.2). The multiplicity of states explodes when increasing species number (Ω(*N*) ∼ 2^*N*^), a signature that would only be observable in simulations or laboratory experiments with many replicas (*N* = 20 species could already generate up to Ω ≈ 10^6^ states). An easier-to-spot fingerprint is that diversity *D* can take many values between 0 and *N*, whereas it is far more constrained in all other regimes (see SI II.G.2). Interestingly, local states are also particularly prone to small shifts after perturbations, as any species crossing its Allee Effect boundary will result in a single-species turnover in community composition (Fig. 4C).

Competition drives the other two regimes at play. Under strong and homogeneous competition, multiple stable states are characterized by the mutual exclusion fingerprint, involving Ω(*N*) ∼ *N* states, each with a single survivor (*D*(*N*) ∼ 1, Fig. 4 blue). Also typical of transitive competition, these states rarely fluctuate [40].

Clique states emerge as competition becomes heterogeneous (Fig. 4, green). The total number of cliques is inherently difficult to estimate and can depend on the properties of the studied system (see SI II.F.2). Across the studied models we find that the number of stable cliques is likely to be higher than system size (Ω(*N*) ∼ *N*^*θ*^, 1 *< θ <* 2, see SI II.G.1), meaning that systems with an initial pool of 50 species could exhibit hundreds of different stable states (Fig. 4B). Although many cliques can emerge, an essential property is that they all tend to have very similar diversity: shifts between cliques might imply some species turnover, but rarely a drastic change in community diversity (see II.G.2). As recently observed in microbial communities [48], the last key signature of cliques is their high tendency to exhibit chaotic fluctuations (*S <* 1, Fig. 4A and SI II.F.2).

Together with these fundamental fingerprints, our tests show the prevalence of a strong inverse correlation between the number of stable states of a system and their resilience to random perturbations *R*, with a predictable lower resilience bound at *R* ∼ Ω^−1^ (Fig. 4C and see SI II.G.3). This puts forward a somewhat obvious but relevant hypothesis in random complex systems: the fragility of a stable state increases drastically and predictably as it becomes surrounded by a multiplicity of other states, as their agglomeration reduces the basin stability of each state.

## DISCUSSION

Four different regimes of multistability emerge in a high-dimensional community model of interacting species, each of the species being assumed to have an Allee Effect. Interestingly, each multistability regime emerges under a clearly identifiable interaction scheme (Fig. 3) and is characterized by a distinct signature (Fig. 4). This raises two questions: Are any of these patterns related to empirical observations in ecology? Do the multistability regimes described here and their fingerprints hold across models of other complex systems beyond ecology?

### Linking the four multistability regimes to ecological evidence

The local multistability regime implies many combinations of present and absent species. Because species dynamics are weakly intertwined in this regime, environmental changes should rarely result in abrupt shifts involving many species losses, but rather gradual changes in the composition of communities (see Fig. 2A and II.G.2). Evidence for local multistability signatures would also imply that the observed patterns are predictable from single-species dynamics (see SI II.A). This scenario is likely related to the perspective of ecosystems as *loose collections of species* [56; 57], which is consistent with observations of gradual species replacements under environmental changes, as identified in forest [58; 59] or benthic ecosystems [60]. As competition increases, the presence or absence of a competitor can prevent the survival of others, hereby reducing the overall diversity of local stable states (Fig. 2B). A related empirical observation is that of *checkerboard* abundance distributions, where species tend to have negative associations with each other [61]. These checkerboard patterns suggest that multiple stable states are influenced by the dominance of different but transitive competitors [62; 63].

A better understood regime concerns the limit-case of mutual exclusion states under strong competition. Because they are easier to engineer, for example via single-resource competition, multiple states with only one surviving competitor have been observed in detail in simpler systems ([64; 65] but see [23]). If stronger competition between bistable species can also lead to global extinction remains untested to our knowledge in many-species experiments.

Our results highlight the emergence of a multistability regime with many clique states under intransitive competition. These states are characterized by holding a similar number of species (Fig. 2C, 4 and SI II.G), meaning that shifts between cliques are characterized by moderate species turnover, but no major change in total diversity (see SI II.G.2). In this context, they can be linked to two related empirical scenarios. First, opposed to the concept of ecosystems as loose collections of species is that of *communities as superorganisms* [57; 66]. In this view, evidence of sharp shifts in community composition indicates that only specific subsets of species might be allowed to coexist [67; 68]. Second, cliques have also be linked to *ecological succession* and related *priority effects* : the processes by which the assembly of a community is affected by historical contingency, so that which of many similar communities is attained depends on the order of species invasions and extinctions [50; 69; 70]. Clique regimes are also linked to the emergence of cyclic and chaotic fluctuations, a pervasive signature of intransitive competition in nature [47; 48].

Finally, the global bistability regime underlying an all-or-nothing regime can be linked to *catastrophic shifts*: large and abrupt transitions between two markedly different ecological states [1; 18]. High-dimensional random systems, however, only appear to show the global bistability signature under a very specific setting of nonlinear and dominantly positive interactions. Accumulated empirical evidence for communities undergoing catastrophic shifts in nature (see *e*.*g*. [3; 8; 71; 72]) does not indicate that these communities are inherently mutualistic, but rather involve an heterogeneous mixture of interaction types. This discrepancy between theoretical predictions and observations hints to a possible missing element in the models studied here. We hypothesize that a key step to bridge this gap is to uncover the role played by empirical network structures, following the observation that the distribution and architecture of inter-species links is neither random nor uniform (see e.g. [36; 37; 73]).

### Equivalent multistability regimes and fingerprints across complex systems

We have replicated the present analysis on four high-dimensional models of complex biological systems, namely Lotka-Volterra interactions [42], the cancer-immune interplay [74], gene regulatory networks [75] and interactions between neural clusters in the brain [76] (see SI III).

We find that the four multistability regimes that emerge in a species-rich community model also do so in these systems. Again, which multistability regimes emerge in each model depends on a simple and generic taxonomy: Global and local states emerge in cooperative or weakly interacting systems with bistable units. Mutual exclusion states and cliques emerge in the presence of competition, depending if it is strong and transitive or highly heterogeneous (see SI III). More importantly, the empirical fingerprints of each multi-stability type are qualitatively equivalent across models (Fig. 4D) and replicate the same taxonomy: linear models without node bistability do not show the global and local signatures (Fig. 4D, Lotka-Volterra), and models without node extinctions do not harbor a mutual exclusion phase (Fig. 4D, neural network).

We hypothesize that these results are also generic to complex dynamical systems beyond biology. Determining the origin of this generality, whether it suggests a fundamental equivalence among complex systems or it arises from the shared simplicity of the models, poses a considerable challenge. Nevertheless, the fact that the same four regimes consistently emerge across models suggests the potential for establishing a general taxonomy of four multistability regimes in random complex systems.

## CONCLUSION

The emergence of multiple stable states in complex systems holds relevant implications across research fields. However, our understanding of this process has been limited by low-dimensional frameworks and generally confined to specific models and assumptions [15]. In the present work, we studied how multistability can emerge in complex systems of randomly interacting units. By analyzing archetypical models of complex biological systems, we identified the emergence of four different classes of multistability. Of these, two arise in cooperative or weakly interacting systems composed of locally bistable units, while the other two manifest under predominantly competitive dynamics. Furthermore, each regime leaves a distinct fingerprint that can be observed and quantified empirically.

In ecology in particular, our work highlights how the current low-dimensional understanding of alternative stable states should scale up when analyzing species-rich communities with multiple interaction types [15]. Each multistability regime can be related to well-established observations in ecology, while unveiling their characteristic signatures provides a tool to further test their occurrence in empirical studies. Moreover, and across the studied models, abrupt shift signatures only emerge under overwhelmingly positive interactions, which does not coincide with natural evidence of catastrophic transitions. This further stresses the need to understand the pivotal role of non-random network structures in explaining community-scale catastrophic shifts in nature [77; 78].

Beyond the field of ecology, our taxonomy suggests a constrained number of multistability behaviors expected to generically emerge in high-dimensional models with random interactions. These results help define a unified framework to understand the nature and conditions by which multistability emerges in complex systems.

## MATHEMATICAL METHODS

### Species interactions

Intraspecies parameters across the models are positive-defined and generated with Log-normal distributions with standard deviation of the logarithm *σ* = 0.05 (Fig. 2A,B, 3 top) and *σ* = 0.8 (3 bottom, see SI II.F.1). Interaction matrices *A* and *B* of dimension *N × N* have intraspecies diagonal vectors *A*_*ii*_ and *B*_*ii*_. Diagonal and off-diagonal terms are generated from different Log-normal distributions (see SI I). Single-layer matrices encompassing both positive and negative interactions are generated from Gaussian distributions (see SI III). We explore the effect of varying the mean (Fig. 2A,B, 3), standard deviation (Fig. 2C) and connectivity (see SI II.F.3) of interspecies interactions.

### Numerical simulations

Numerical simulations are described in detail in SI I and python codes are available at https://github.com/GuimAguade/A-taxonomy-of-MSS-in-complex-ecological-communities. We define a model by setting a dynamical equation, intraspecies and interaction parameters and a finite number of simulations *s* to be run (50 *< s <* 400 for fFigures 2-4 and *s* = 10^4^ for figure 4B). For each, we generate random initial conditions (see SI I.C), integrate the system of differential equations with a Runge-Kutta method of order 5(4) [79], store the final state of the system and test it for stability against time and species migrations (see SI I.D.1).

### Stable states, fingerprints and perturbations

Simulations allow us to count the number of different states reached for a given system (Fig. 3, SI I.D.2) and compute their properties (Fig. 4, SI II.G.1). After a stable state is reached, we perturb it with a random Gaussian change in each species abundances Δ*x*_*i*_ and compute if the dynamics stabilize to the original species composition (Fig. 4C, SI I.E, II.G.3).

## ACKNOWLEDGEMENTS

The authors thank the members of the BioDICée team at ISEM and the members of the Institut Natura e Teoria en Pireneus (INTP) for discussions and support throughout the project. G.A.-G. thanks L. Arola, I. Lajaaiti, B. Pichon and R. Solé for insight and feedback and A. Vives and J. Leigh for support on graphical design. Special thanks to J. G. Cole and B. Parham for inspiration. G.A.-G. was supported by a 2022 postdoctoral fellowship of the Fundación Ramón Areces. S.K. was supported by the grant ANR-18-CE02-0010-01 of the French National Research Agency ANR (project EcoNet).

## Notes

https://github.com/GuimAguade/A-taxonomy-of-MSS-in-complex-ecological-communities

### Competing Interest Statement

The authors have declared no competing interest.

