## Supporting Information for "A taxonomy of multiple stable states in complex ecological communities"

### Supporting Information to:
### A taxonomy of multiple stable states in complex ecological communities

Guim Aguadé-Gorgorió,[1] Jean-François Arnoldi,[2] Matthieu Barbier,[3] and Sonia Kéfi[1,4]
[1] *ISEM, University of Montpellier, CNRS, IRD, EPHE, Montpellier, France*
[2] *Centre for Biodiversity Theory and Modelling, Theoretical and Experimental Ecology Station, CNRS and Paul Sabatier University, 09200 Moulis, France*
[3] *PHIM Plant Health Institute, University of Montpellier, CIRAD, INRAE, Institut Agro, IRD, 34090 Montpellier, France.*
[4] *Santa Fe Institute, 1399 Hyde Park Road, Santa Fe, NM 87501, USA*

(Dated: August 28, 2023)

The structure of the following supplementary material is as follows:

First, we present and define the fundamental problem under study and describe the structure of the numerical simulations to explore it.

Second, we study in a stepwise process how introducing interactions modulates multistability landscapes for the reference model of the Main Text, and how mathematical techniques can help in understanding each of the steps. We also describe the fingerprints characteristic to each multistability type.

Third, we present and study a set of models describing different complex systems, and compare the different types of emerging multistability to the results of the Main Text.

#### CONTENTS

I. Model definition and numerical simulations 3
- A. Description of the problem 3
- B. Backbone of the numerical implementation 5
- C. Generating initial conditions, intra-species parameters and interaction networks 6
  1. Initial conditions 6
  2. Intra-species parameters 6
  3. Species interactions 6
  4. Ecological networks 7
- D. Testing for stability and comparing across observed states 9
  1. Stability 9
  2. Comparison between stable states 10
- E. Perturbations 11

II. Studying multistability under different interaction types, strengths and heterogeneity 12
- A. Independent bistable species 12
- B. Bistable species, mutualistic interactions 14
  1. Dimensionality reduction under homogeneity constraints 14
  2. Intertwining between global and local effects 15
  3. Visualization in 2D systems 15
- C. Bistable species, competitive interactions 17
  1. Failure of dimensionality reduction techniques 17
  2. Intertwining between exclusion and local bistability effects 17
  3. Mutual exclusion and community-wide extinction 18
  4. Visualization in 2D systems 18
- D. A note on predator-prey interactions and oscillations 19
- E. Bistable species, mixed interaction types 21
  1. Effectively neutral interactions 21
  2. Relaxation times and fluctuations at the mutual exclusion threshold 22
- F. Bistable species, heterogeneous interactions 23
  1. The heterogeneity threshold and the emergence of cliques 23
  2. Stability of cliques and chaotic fluctuations 24
  3. Heterogeneity at the network structure level 27
- G. Fingerprints of the four multistability families 28
  1. Number of states and system size 28
  2. Diversity of states and species overlap 29
  3. Number of states and their resilience to perturbations 31

III. Other models of complex system multistability 33
- A. Generalized Lotka-Volterra interactions 33
- B. Cancer-Immune interactions 36
- C. Gene regulatory networks 39
- D. Dynamics of neural networks 41

Bibliography 43

References 43

Supplementary figures 47

#### I. MODEL DEFINITION AND NUMERICAL SIMULATIONS

##### A. Description of the problem

In the present paper we discuss the presence and nature of multiple stable states in complex dynamical systems. The central motivation of the research lies within the concept of low-dimensional alternative stable states and regime shifts in community ecology (Scheffer *et al.*, 2001). Our question is to understand how multiple stable states emerge in systems with many interacting units, and how does this change our low-dimensional understanding of ecological shifts (Kéfi *et al.*, 2022). The fundamental task is therefore to study a set of ordinary differential equations of the form

$$\frac{dx_i}{dt} = F_i(x_i) + \sum_j^N G_{ij}(x_i, x_j) \tag{1}$$

In the particular setting of ecology, $x_i$ is the abundance of species $i$, that has self-regulatory dynamics $F$ and interacts via function $G$ with a pool of $N$ different species. This variables might have alternative meanings, such as gene activity or cellular density (see section III), and equation (1) is likely to take a variety of nonlinear forms depending on the model under study.

Our central task is to find and study the presence and characteristics of attractor states, namely vectors $\boldsymbol{x}^* = (x_1^*, x_2^*, ..., x_N^*)$ such that

$$F_i(x_i^*) + \sum_j^N G_{ij}(x_i^*, x_j^*) = 0 \qquad \forall i, j \in [1, N] \tag{2}$$

Uncovering and counting stationary states in high-dimensional dynamical systems governed by interacting elements, be it Boolean networks (Kauffman *et al.*, 1993) or random energy landscapes (Derrida, 1981), is a central and remarkably complex task across different areas of physics (Fyodorov and Khoruzhenko, 2016). It often involves the use of analytical tools much beyond the scope of the present article, such as the replica method (Mézard *et al.*, 1987) or the Kac-Rice formula (Fedeli *et al.*, 2021; Ros *et al.*, 2019a).

More specifically, and originally motivated by catastrophic shifts between stable states in ecological systems (Scheffer *et al.*, 2001), we are here interested in finding those attractors that are stable. This further complicates the problem, as we have to identify those $\boldsymbol{x}^*$ values for which the Jacobian matrix of the system, with elements

$$J_{ij} = \left( \frac{\partial}{\partial x_j} \frac{dx_i}{dt} \right)_{\boldsymbol{x}} \tag{3}$$

has negative eigenvalues (Strogatz, 2018). Here, we will make use of simple analytical approximations when possible, and more often use numerical simulations to solve (2,3), as explained below. We refer the reader to interesting literature on analytical approaches to evaluate stability in ecological communities in section II.F and also in a vast literature discussing it (see e.g. (Allesina and Pascual, 2008; Allesina and Tang, 2012; Barabás *et al.*, 2017; Domínguez-García *et al.*, 2019; Donohue *et al.*, 2016; Grilli *et al.*, 2016; Kéfi *et al.*, 2019; Mougi and Kondoh, 2012; Yodzis, 1981) to name but a very few).

There are many relevant characteristics of a system and its potential to harbor multiple stable states we could study. To simplify things and convey the main results in a clear voice, we here focus in a small set of state properties.

- First, for a given dynamical system with $N$ species and defined parameters, **we want to find if there are stable states**. In particular, we want to count the amount of random initial conditions that end up in stable compositions instead of exponential outgrowth, cycling behavior or persistent fluctuations. We define $S$ as the fraction over $s$ simulations that progress into a stable attractor. Scenarios with $S = 1$ indicate all observed trajectories end up in a stable state, while $S = 0$ indicates no stable basins have been found and other dynamics are at play.

- Of the $s \times S$ simulations that end up in stable community compositions, **we want to find the number of different states $\Omega$ that can be observed**. If solving the system with numerical simulations, this will not always be the total number of states the system can be in, but only the subset we can find by generating a finite amount of random initial conditions. Despite this might give a reduced or partial view of the whole multistability landscape, the exercise is closer to that of experimental tests where only a limited number of replicas can be performed.

- Once stable states are observed and counted, **we want to evaluate certain basic properties of these states**. Because species $i$ and $j$ are not specified and there are no necessarily keystone species, the main property of a state that could relate to its biological or ecological function is diversity $D$: the number of species that have a positive abundance $x_i^* > 0$ and hence are present in the stable state. Total biomass is defined as $X = \sum_i x_i$. However, parameters in the studied models will not always be sorted from real experimental or empirical datasets. Henceforth, the presence or extinction of a species will be more relevant than at what biomass it stabilizes. We will then use total biomass measures when useful to explain qualitative underlying dynamical behaviors, such as those observed in Figure 2 of the Main Text.

- As discussed below, we will later introduce random perturbations as a measure to analyze potential state shifts and state resilience. This will allow us to **evaluate a complementary subset of properties of states under perturbations**. First, we will perturb stable states and compute how often they relax back into the original composition. The fraction of times this happens will provide a proxy for *basin stability* (Menck *et al.*, 2013), an analogue to linear stability that allows us to understand the proximity of other stable states and the basin of attraction of each. For those states that do transition, we will measure the Jaccard $J$ distance, the total species turnover between two states, as the simplest measure of state difference or *distance* (Jaccard, 1912), together with species overlap $O$, indicative of how many species are shared between the original and perturbed state. We will also capture the size of the perturbation that resulted in a shift and the correlation between these different perturbation properties (see section II.G for a detailed description).

Less central metrics will be introduced and described when necessary. In summary, we will here find how often a complex dynamical system reaches stable states after $s$ initial conditions, how many different states can be observed $\Omega$ and measure their diversity $D$. We will later perturb these states and count the fraction of shifts and the properties of the newly found states.

##### B. Backbone of the numerical implementation

Because the analytical problem (1-3) will in general be very complex, we generally rely on numerically solving the equations with python. All codes are available at
https://github.com/GuimAguade/A-taxonomy-of-MSS-in-complex-ecological-communities.

A single simulation implies solving an initial value problem, meaning that the dynamical equation (1) is defined together with initial conditions and the duration of the dynamical process, and we have to integrate the dynamics of the system. The backbone of the process follows a recursive implementation:

1. A model is fully defined. Doing so involves setting a specific dynamical process through $F_i$ and $G_{ij}$ and all the parameters involved (see below).

2. We set the number of repetitions or simulations $s$ for which the system will be run and the elapsed time or duration $\Delta t$ of a simulation. Each simulation involves the following:

   - For each simulation, we generate an array of random initial conditions $\boldsymbol{x}(t=0) = (x_1(0), x_2(0), ..., x_N(0))$ (see below).
   - We now have a complete initial value problem $(d\boldsymbol{x}/dt, \boldsymbol{x}(0), \Delta t)$, that we integrate with a Runge-Kutta method of order 5(4) (Dormand and Prince, 1980). This is implemented via the `solve_ivp` function of the `scipy.integrate` library.
   - The final state at $t = \Delta t$ is saved. We test its stability through time and against species invasions (see below).
   - If the state is not stable, we increase by one the count for unstable simulations. If the state is found stable, we see if it is new or else we have seen it before. If it's new, we save it in the list of observed states. If it has been seen before, we increase by one the number of times it has been seen.
   - Finally, if the state is stable, we perturb it with a non-infinitesimal change in species abundances (see below), meaning that we are not testing for linear stability but rather for the presence of other basins of attraction. We save the newly found state and compare it to the unperturbed one.

3. After the above process, we have a list of $\Omega \leq s$ different stable states for a given model, and how often each has been seen. This will allow us to compute the above "central" metrics of our problem, namely the fraction of stable simulations $S$, the number of observed stable states $\Omega$ and their diversity $D$. We also measure different properties of the possible state shift.

The above process of finding the number of stable states of a system after $s$ simulations is then used to numerically solve all the questions of the present work or test them against analytical results. In particular, we can see how the central metrics are affected by modulating species interaction strengths (Figures 2 and 3 in the Main Text) or network connectivity, the choice of initial conditions (see sections II.A and II.B.2), the effect of perturbations (Figure 4 in the Main Text) or the definition of the model itself (see section III).

##### C. Generating initial conditions, intra-species parameters and interaction networks

###### 1. Initial conditions

At each simulation we generate a set of random initial conditions to test their dynamical evolution. This allows us to both explore the landscape of possible attractor states, and also obtain a hint on the basin stability of a state, by computing the volume or fraction of initial conditions that fall into it or later compute the necessary change in species abundances to escape it (see (Menck *et al.*, 2013)).

Random initial conditions can be generated in many ways, and we here choose 2: (1) sample the phase space without correlations between species abundances and (2) sample the phase space by fixing the total initial abundance, as a proxy for a more realistic experimental setting where total abundance, but not how it is distributed among species, is known.

(1) is obtained via random uniformly distributed variables, that we choose to be centered at the bistability threshold of each model ($x_i(t=0) \in U[0, 2x_i^{*,U}]$, see section II.A and section III). This allows us to set the probability of species survival and extinction equal if species do not interact, and will test useful to compare results across models (see section III). (2) is defined by a Dirichlet distribution, that allows us to set a fixed total initial abundance and then randomly distribute it across species with evenness controlled by scale parameter (Kotz *et al.*, 2004). Except for particular aspects (see section II.A), the method of sorting random initial conditions will not be relevant to the results when compared to the amount $s$ of initial conditions generated, and we mostly stick to method (1) for simplicity. Method (2) will mostly be discussed when observing how the diversity of local multistability states depends on initial conditions.

###### 2. Intra-species parameters

Because of the way the models are defined, intra-species parameters, as well as most interaction parameters across models (see section III for several exceptions), are positive-defined. This is to ensure, for example, that $-d_i x_i$ remains a death term, so that all $d_i$ terms have to be positive. Moreover, we want to control the amount of heterogeneity across species, *e.g.* how different their death parameter is. To do so, we generate all intra-species parameters from log-normal distributions to ensure positivity, implemented within the `random.lognormal` module of the `numpy` library.

Because the main purpose of the work is to draw qualitative insights and we have no experimental system or data to fit, we choose intra-species parameters so that we can control the probability of survival or extinction resulting from single-species bistability when present (see sections II.A and III). This means that we will set the mean of $\{\gamma_i, d_i, A_{ii}, B_{ii}\}$ in the reference model so that bistability is ensured, except for Figure 3 (top) where $d_i$ is increased beyond the threshold that would allow the species to survive, something that could also be attained by setting $A_{ii} = 0$ (see section II.A for the analytical expression). Intra-species heterogeneity, modulated by the standard deviation $\sigma^{ii}$ of the above distributions, will allow us to compare coarse-graining methods to more realistic scenarios (see section II.B.2 and Fig. 4) but, because we will want to focus in the role of interactions and their heterogeneity $\sigma^{ij}$, we choose to define a relatively homogeneous set of intra-species parameters with $\sigma^{ii} = 0.1$, that is still within the uniformity constraints of (Gao *et al.*, 2016; Laurence *et al.*, 2019), and only change it in specific sections of this Supporting Information (see below). For the rest of the paper, the standard deviation of inter-species interactions is defined as $\sigma^{ij} = \sigma$.

Mean intra-species parameters that ensure single-species bistability for the model studied in the Main Text are: $\{\langle\gamma_i\rangle = 1.0, \langle d_i\rangle = 0.1, \langle A_{ii}\rangle = 0.5, \langle B_{ii}\rangle = 0.1\}$. Because we want to focus on the role of species interactions and single-species effects are well understood (see section II.A), these parameters will be fixed throughout our analysis and in all figures unless stated otherwise.

###### 3. Species interactions

Interspecies interactions are introduced via well-known *community matrices* $A$, $B$ (and $P$ in section II.D), a central concept in theoretical ecology. In species-rich communities, however, inferring the weight of a vast number of approximately $N \times N$ interactions is likely to be an utterly complex task. Here is where the theory of *disordered systems* came to place as an extremely useful method to tackle ecological communities as complex systems. Originating in the physics of glasses (Mézard *et al.*, 1987), the approach has been used in a wide variety of fields and has provided powerful insight in complex systems analysis. We refer the reader to (Barbier and Arnoldi, 2017) as a step-by-step account for how the theory of disordered systems seats in-between different approaches in ecology.

The central idea of the disordered approach is that, due to system complexity, we can not (or do not want to) assign any clear role to individual species nor their interactions. Instead, we focus on simple information regarding the interaction matrix as a whole, and try to uncover how generic community-scale patterns emerge from questions such as "what is the average strength of interspecies interactions?" or "how different these interactions are among species?". Because of the lack of structure of such a method, it allows to uncover what type of fundamental patterns can emerge generically from high-dimensional systems or, instead, what behaviors can only emerge if additional and non-trivial structure would be considered. This makes it a particularly useful framework to start our research: instead of exploring a myriad of ecologically specific organizations, such as food-web architecture (Neutel and Thorne, 2014) or parasite-host interaction motifs (May and Anderson, 1990), we first aim at asking: if the system is *essentially* composed by many species that interact in multiple and generic ways, what type of multistability scenarios can emerge? Are they consistent with ecological evidence, or is there some pattern that random interactions cannot explain?

It is within this context that the seminal paper by Robert May introduced the possibility of defining the weights of interaction matrix elements in species-rich theoretical models as random parameters (May, 1972b), considering that matrices are likely to be so large that they could encompass a probabilistic distribution of $A_{ij}$ values. This came in hand within the framework of Random Matrix Theory, that following the same simplistic approach to complexity provided a useful mathematical machinery to understand fundamental properties of ecological communities (see e.g. (Allesina and Tang, 2012; Gibbs *et al.*, 2018; Grilli *et al.*, 2016; May, 1972b) among many).

In the original works of (May, 1972b) or (Allesina and Tang, 2012), random matrices where used as a proxy for the system's Jacobian, a tool to study linear stability. Here we consider random interaction matrices within the wider framework of disordered systems techniques such as those applied in community assembly (Barbier *et al.*, 2018), so that we do not necessarily focus on interaction strengths around a systems' equilibrium. Moreover, and depending on the model under study, we will take two similar approaches. In models with a single layer of interactions (see section III), these will be built through a single matrix $A$, with interaction terms $A_{ij}$ sorted from a Gaussian distribution with controlled mean $\mu$ and variance $\sigma_2$. This means that interactions with different signs are likely to coexist, and the weight of each is strongly determined by $\mu$. This is a common procedure for the original random matrix approaches (Allesina and Tang, 2012; May, 1972b) as well as for the generalized Lotka-Volterra model of ecological interactions (see section III.A and (Bunin, 2017)). The diagonal terms of the matrix, related to species self-regulation or self-cooperation, are the $A_{ii}$ terms discussed above.

We also study models in which different interaction types are separated into different layers $A$ and $B$ (see Main Text), also because they can follow different functional responses (Pilosof *et al.*, 2017). In the model of the Main Text, for example, mutualism follows a saturating expression (see section II.B) while competition is linear (see section II.C). This means that they could not be written in a single matricial form, and, as opposed to other models where interactions can be mixed (see e.g. (Qian and Akçay, 2020)), here two species $i$ and $j$ can *simultaneously* interact via positive and negative processes, such as secretion of mutually beneficial growth factors in bacteria together with competition for resources (Griffin *et al.*, 2004). Although these two process are likely to be intertwined up to a certain level (Griffin *et al.*, 2004), being able to modulate the strength of each separately allows us to explore the whole landscape of possible multilayered interaction strengths (Fig. 3 in the Main Text). The multilayer scenario proposed by Eq. (1) in the Main Text implies that now $A$ and $B$ are positive-defined, and we generate off-diagonal values again from log-normal distributions.

###### 4. Ecological networks

The other key element of species interactions in complex ecosystems is the structure of species connections (Sole and Montoya, 2001). We now know that not all species interact, and the links between those that do can allow for non-trivial structures and complex interplay of network motifs. A large body of work has been devoted to uncover the nature of these structures and their evolutionary origins and functions (see e.g. (Bascompte, 2010; Ings *et al.*, 2009; Pilosof *et al.*, 2017; Sole and Montoya, 2001; Valverde *et al.*, 2018) among many others). In the field of species-rich community ecology, however, obtaining analytical results when $A$ has a nontrivial shape remains very complex, if not unfeasible. Moreover, theoretical works that are relevant to our research point towards the possibility that network structure, if it doesn't deviate extremely from the disordered scenario, does not seem to play a major role when compared to system dynamics (Barbier *et al.*, 2018; Gao *et al.*, 2016). On the other hand, however, it does seem that alternative network structures can either favor or disrupt the presence of strong feedback loops within a community (Kéfi *et al.*, 2016; Neutel and Thorne, 2014) and even modulate the presence of global bistability thresholds (Lever *et al.*, 2014).

Because we frame our models within the disordered systems approach, and hence do not assign specific roles to particular species, we start by not defining any particular network structure, such modules or motifs (Bascompte and Stouffer, 2009), nor hierarchical species classifications such trophic levels (Pimm and Lawton, 1977). Following a classical approach in theoretical community ecology (May, 1972b), in the present work we define the structure of ecological networks following an Erdős-Rényi graph (Erdős *et al.*, 1960). Specifically in the Erdős-Rényi-Gilbert version of the model (Gilbert, 1959), each edge $j \to i$ (and hence each $A_{ij}$ value) has a fixed probability $p$ of being present or absent (Fienberg, 2012), and the expected total number of edges (and non-null elements in $A$) is $\binom{N}{2}p$ in the undirected graph and $pN(N-1)$ in the directed ecological network. As presented in (Fried *et al.*, 2016) and discussed in (Fried *et al.*, 2017), this makes the graph in our approach dense, in the sense that more species lead to more links, whereas in the Erdős-Rényi formulation the probability of interacting scales as $1/N$, so that every species has on average a fixed number of interactions (Fried *et al.*, 2017). Here of course the diagonal terms are $A_{ii} > 0$, represent intra-species dynamics and don't alter the Erdős-Rényi structure of the community. In this random approximation to ecological structure, the only parameter modeling network structure will be species probability of connection, $p$, defined as the probability that two species effectively interact (May, 1972b).

When generating the matrix elements $A_{ij}$, a prior call will be to throw a random number between 0 and 1 and see if it falls below $p$. If not, $A_{ij} = 0$ and species $i$ and $j$ will not interact. This will prompt the discussion of section II.F.2, where we will see how reducing connectivity, by adding 0's in an otherwise homogeneous matrix, acts in a very similar qualitative way as increasing parameter heterogeneity. We leave for future work the study of how non-trivial structures, by deviating from this disordered limit, can participate in novel ways in creating or annihilating stable states of species coexistence.

##### D. Testing for stability and comparing across observed states

As discussed in I.B, the central point of our numerical implementation is to generate a model, integrate it $s$ times, and see if (and how many) different stable states appear. As previously mentioned, analytically solving such a problem becomes very hard in high-dimensionality, and would require us to restrict to specific model constrains (see section II.F). On the other hand, two key aspects of the numerical procedure are (1) to evaluate if a final state of the dynamical process is indeed stable, and (2) to properly compare if two of such stable states are equivalent.

###### 1. Stability

The concept of ecological stability represents a long-standing debate (Donohue *et al.*, 2016; Kéfi *et al.*, 2019). Many ecological communities, even under environmental or anthropogenic pressures, seem to be persistent in time (Fernández *et al.*, 1999) or resilient to perturbations (Dakos and Kéfi, 2022), while others seem to fluctuate without species abundances settling into specific values (Lundberg *et al.*, 2000). But accurately describing and quantifying what stability means, and how we can assess if a state is more or less stable than another, is far from trivial. One of the main reasons is the complex and high-dimensional nature of what stability is and how it can be measured: stability can refer to different function- or composition-level responses to different types of perturbations, and system-wide or single-species metrics, evaluated through abundance or time, can provide different and even contradictory answers (Domínguez-García *et al.*, 2019).

We do not aim here to propose a novel framework or methodology to assess ecological stability as done for example in (Allesina and Pascual, 2008; Allesina and Tang, 2012; Arnoldi *et al.*, 2016b; Barabás *et al.*, 2017; Domínguez-García *et al.*, 2019; Donohue *et al.*, 2016; Grilli *et al.*, 2016; Kéfi *et al.*, 2019; Mougi and Kondoh, 2012; Yodzis, 1981). Our goal is to provide a simple analysis of whether an initial value problem ends up in an attracting species composition that does not change in time. To do so, we define a stable state without necessarily testing for linear stability to infinitesimal perturbations with the Jacobian matrix. Instead, we put the focus on whether a final state of a simulation (a) remains the same in time and (b) is stable to species reinvasions.

The first point is straightforward to test in mathematical theory: it would suffice to settle two time windows and check if $x_i(\Delta t_1) = x_i(\Delta t_1 + \Delta t_2)\ \forall i$ for a single initial condition vector. Because some of the studied scenarios require long times to stabilization, we set $\Delta t_1$ very long, often $\Delta t_1 > 10^3$ and even in the order $10^4$ in regions close to critical thresholds where relaxation times are slowest (dictated from observing relaxation times across the models, see e.g. Fig. 8 for a visual example), and a much shorter $\Delta t_2 = 10^1$.

However, here is where one potential weakness of numerical estimation comes into place: because there is constant species remigration and integration is discretized, a mathematically stable state can be accompanied by a small proportion of noise-like fluctuations, and asking if $x_i(\Delta t_1) = x_i(\Delta t_1 + \Delta t_2)$ at the floating-point precision does not always work. To face this problem and avoid the system to detect equivalent states as different, we have to define a dissimilarity threshold, below which we admit that two states, even if slightly different, are considered the same. With this tool, we can differentiate between two equivalent states and two states that are sufficiently different and so indicate that a system has not stabilized.

In the present research we test two possible methods. Both start from the idea that a minimum amount of fluctuation has to be neglected. We want to compare the two states $\boldsymbol{x}$ after $t = \Delta t_1$, and $\boldsymbol{y}$ starting at initial condition the previous state ($\boldsymbol{y}(t = 0) = \boldsymbol{x}$) and continuing for a duration of $t = \Delta t_2$. The first straightforward option is to set a value $\delta$, based on estimating fluctuation size (see e.g. Fig. 1), so that if

$$|x_i - y_i| > \delta \tag{4}$$

it means that the species $i$ in both states has to be considered different. From numerical estimates (not shown), our computational framework works best around $0.02 < \delta < 0.08$ (not shown). Below, equivalent states can be considered different because of very small differences resulting from floating point precision or recursive species extinction and remigration. Above, the system might not be able to account for relevant though small fluctuations. From here onwards we set $\delta = 0.05$ to maintain homogeneity.

Because imprecision fluctuations appear to scale with species abundance (not shown), another common option would be to define an index by accumulating the relative biomass difference at the species-level

$$I^i_{x.y} = \frac{2|x_i - y_i|}{x_i + y_i} < I_M \tag{5}$$

Here $I_M$ is the maximum relative fluctuation that we have found for a single-species in a stable state and is set to $I_M = 0.01$, as visually shown in the example of figure 1. States where total species dissimilarity after the second time window is larger than $N$ times this value are considered to not have stabilized. As we will later discuss, this will happen in two different settings: under exponential growth, cycles and chaos, but also in scenarios where relaxation time is particularly long and exceeds a reasonable $\Delta t_1$ (see section II.E.2). The second method proves more efficient across different scenarios, as fluctuations often scale with species abundance and it does take that into account. However, it can also fail when comparing species that have particularly small or negligible abundance. In this setting, small fluctuations can become amplified by a near-null denominator, making the system evaluate two almost-extinct states as different.

Interestingly, the complementary approach of using a relative dissimilarity index is similar to the Gini inequality coefficient that has been previously used in theoretical ecology as a means to compare the differences between states along an environmental gradient (Liautaud *et al.*, 2019). However, for very small stable abundances below $x_i = 1$, the dissimilarity index starts to diverge and comparisons fail (as seen for example in the generalized Lotka-Volterra model in section III.A). Because the absolute difference method is stable across large and small $x_i$ values, we choose to use both for comparison but present all results that follow using the first.

With this process we test the stability of a state against time, beyond very small differences in species abundances. Another way to test if a state is truly an attractor of the system is to test it against species migration. This is because it could be that a state appears persistent in time though it is unstable in the direction of a specific species growth. If particular initial conditions or even integration errors generate an extinction of that species, we would be counting states that are not stable against all possible intrinsic dynamical challenges. To avoid this error, we incorporate a constant rate of species immigration to our original system

$$\frac{dx_i}{dt} = F_i(x_i) + \sum_{j}^{N} G_{ij}(x_i, x_j) + m_i \tag{6}$$

This implies that all species have a constant -but very small- influx into the community, and is often implemented in many ecological community models (see e.g. (Bunin, 2021) or the original island biogeography work by (Kessler and Shnerb, 2015)) . Here we implement migration by indirectly imposing a lower limit to species abundance, beyond which $x_i$ cannot keep decreasing. Now, any final state is constantly tested against the potential growth of all latent species. If it remains the same in time, and all potential invaders do not grow or alter the abundance of surviving species, we have a two-step persistence test that will prove sufficient to the analysis that comes below.

###### 2. Comparison between stable states

So far we have integrated the dynamics of the system for given initial conditions and tested if the observed endpoint is a stable state against time and species invasions. The key task of the present work is to analyze how many different stable states can a system engender, and several properties and fingerprints of these stable states.

Once again, it is analytically straightforward to evaluate if two stable states $\boldsymbol{x}^*$ and $\boldsymbol{y}^*$ are the same. Because they are encoded as vectors, it will suffice to compute if $x_i = y_i \; \forall i$. So, evaluating if two stable states are the same or different ones becomes the equivalent problem as to evaluate if a state has remained the same after a given time window.

Because up to hundreds of different states can be stored and compared during simulations, one tested option is to start by reducing the floating point precision up to 3 decimal points, beyond which integration fluctuations can appear. This is easiest to see for total coexistence, for which only one possible state should exist. We have not tested the computational complexity of this in depth, but at first appearance it does not seem a major improvement in reducing the elapsed simulation time and we have not used it throughout simulations.

All in all, once a final state is reached, we ask if it has previously been seen. To do so, we compare each species one by one with the absolute difference method $\delta$ discussed above. If each and all species differences are below $\delta$ for the two compared states, they are considered the same. The system is therefore weak in model settings where one could have mathematically different states separated by particularly small $x_i - y_i$ differences. However, this will not be the case across the studied models where states are separated by sufficiently large differences.

##### E. Perturbations

Once a stable state is reached, our final test is to perturb it and evaluate its response in the simplest possible terms. Also, we do not aim here at evaluating linear stability by performing an infinitesimal displacement. Rather, we aim at obtaining a Monte-carlo approach to the basins of stability of the possible stable states. Similar to the traditional ball-in-a-cup metaphor, these will be proportional to the perturbation width needed to shift towards other states (Menck *et al.*, 2013).

Perturbations (and the resilience metrics they encompass) can take many forms in ecosystems (Bender *et al.*, 1984; Domínguez-García *et al.*, 2019). To evaluate basin stability and the nature of shifts, we are interested in maintaining the multistability landscape fixed and seeing how trajectories evolve in it (Beisner *et al.*, 2003; Menck *et al.*, 2013). Because of this, here we focus on pulse perturbations in the form of random changes in population abundances.

After reaching a stable state $\dot{x}_i = 0 \; \forall i$, we perturb the system with $x_i + \delta x_i$, where for simplicity $\delta x_i$ are taken from a Gaussian distribution with mean at zero and variable standard deviation, here $\sigma_P \in [0, 3]$. This means we are a priory not thinking about an extinction-like event, but a more general scenario where populations can either grow or decay. We will modulate the standard deviation of such perturbation as a means to study how states react to smaller or larger changes.

After the perturbation, we integrate the trajectory of the perturbed state for an equivalent time window $\Delta t_1$. We evaluate using the absolute difference method if the final state that is reached is the same of the unperturbed one, or a new one. This allows us to obtain a new set of metrics: how often a system returns to the original state (an indirect proxy for basin stability and resilience $R$, see Fig. 4 in the Main Text). Other metrics that we have studied involve how large was the perturbation that produced a shift, and also how different the two states are as a proxy for the width of the shift observed.

To do this comparison, we followed previous research that used the Jaccard index $J$ to compare the presence of surviving species in both states (Liautaud *et al.*, 2019). The Jaccard index simply adds a one each time one species is present in one state but not in the other and vice versa. It is, therefore, a measure of total turnover between two states. Because our species do not have particular traits, are not labeled and biomass changes might not be representative of ecosystem function, total turnover might be a representative way to understand how different two states really are and how different their function could be. A complementary metric is of course to measure the fraction of species overlap $O$, how many species are shared between the two states divided by their total diversity. This gives another view of whether the shift brings the system to a consistently different state or a similar one, bringing up again the potential discussion of alternative stable states as gradual species changes or sharp community transitions (Liautaud *et al.*, 2019).

After these test we concluded that simpler metrics (see Figure 4 in the Main Text) provided more interesting signatures (see section II.G). On the contrary, measuring Jaccard distance between states and their overlap did not result in very well-defined clustering of multiple stable state (MSS) categories. Beyond the obvious result of global bistability involving a total turnover, the rest of MSS categories resulted in a similar distribution of potential transitions with small $J$ (not shown) and $O$ (see section II.G.2). The rest of the analysis therefore will only discuss the role of $R$, the probability that states are recovered after a random perturbation, and $O$, the number of species that are shared between two different states after a perturbation.

#### II. STUDYING MULTISTABILITY UNDER DIFFERENT INTERACTION TYPES, STRENGTHS AND HETEROGENEITY

##### A. Independent bistable species

The starting point of the analysis for the Main Text as well as other models (see section III) is to understand how does a model behave in the absence of inter-species interactions, meaning that system (1) in the Main Text now writes

$$\frac{dx_i}{dt} = x_i \left( A_i \frac{x_i}{\gamma_i + x_i} - d_i - B_i x_i \right) \tag{7}$$

$A_i$ is the maximum growth rate of species $i$. Mutualism inducing an AE is included by considering that species abundance increases growth rate, saturating at high abundances to this maximum $A_i$ value. The parameter $\gamma_i$ determines the amount of $i$-individuals necessary to achieve half of the maximum growth rate, as a proxy for how difficult it is for them to cooperate. Species are assumed to have a natural linear death rate $d_i$ when alone, and also compete for space or resources at strength $B_i$. The following approach can be replicated for the other models considered below.

Before we include species interactions, in this first approach $\dot{x_i} \neq f(x_j)$ so that each species is independent from the rest. This will prove useful to later understand arising multistabilities in the community. This type of single-species bistable dynamics has been thoroughly studied (Courchamp *et al.*, 2008). In particular, each species always has a stable state at $x_i^* = 0$. Beyond that, it can be that a stable-unstable $(s, u)$ pair appear, with values

$$x_i^{*,s} = \frac{1}{2B_i}(A_i - B_i\gamma_i - d_i + \sqrt{(A_i - B_i\gamma_i - d_i)^2 - 4B_i\gamma_i d_i}) \tag{8}$$

and

$$x_i^{*,u} = \frac{1}{2B_i}(A_i - B_i\gamma_i - d_i - \sqrt{(A_i - B_i\gamma_i - d_i)^2 - 4B_i\gamma_i d_i}) \tag{9}$$

However, this states are not always present. We will observe them if the square root is real, so that

$$(A_i - B_i\gamma_i - d_i)^2 - 4B_i\gamma_i d_i \geq 0 \tag{10}$$

from where it is easy to see that

$$A_i > A_i^c = B_i\gamma_i + d_i + 2\sqrt{B_i\gamma_i d_i} \tag{11}$$

The meaning of this condition is straightforward: if growth arising from mutualism $A_i$ is not larger than a function of all parameters detrimental for growth $(B_i, \gamma_i, d_i)$, species $i$ will go extinct $(x_i^* = 0)$. If $A_i$ fulfills (5), the species can either be at $x_i^* = 0$ or $x_i^{*,S}$, a bistable scenario. In this scenario, whether species $i$ is at one or another state will only depend on initial conditions: if $x_i(t = 0) < x_i^{*,U}$, species will fall to zero, and vice versa (Fig. 2top: single species bistability). The fact that a species can be at two different stable states for the same parameters, only depending on its initial abundance, poses the underlying mechanism for so-called regime shifts.

How will a community built upon $N$ of these species behave? Because species do not interact, the community will be a realization of individual stable states. If (5) is fulfilled for all $i$, then the full system could be at any combination of present and absent species, with total biomass range

$$X^* = \sum_{i=1}^{N} x_i^* \in \left[0, \sum_{i=1}^{N} x_i^{*,s}\right] \tag{12}$$

At the limits of this range, there is one state with zero species, and one with all species present. In between those states, however, there is a large multiplicity of community states: with species behaving as binary present-absent entities, there are $2^N$ different compositions or stable states. For a community built only of $N = 20$ species, there are more than one million states with different present and absent species.

In which of these states the community will be observed depends on the initial conditions $x_i(t = 0)$. A possibility would be that individual initial conditions are independent from one another, rendering a Poisson binomial distribution in the number of surviving species: Because species and initial conditions are independent, the survival/extinction of a species is simply a Bernoulli experiment. If initial conditions are sorted from a uniform random distribution in $[0, a]$, the probability that species $i$ survives is

$$p_i = \frac{1}{a}(1 - x_i^{*,u}) = \frac{1}{2aB_i}\left(2B_i - A_i + B_i\gamma_i + d_i + \sqrt{(A_i - B_i\gamma_i - d_i)^2 - 4B_i\gamma_i d_i}\right) \tag{13}$$

and, because each species might have different probability of survival, the combination of trials becomes a Poisson binomial distribution where the survival of $k$ species is dictated by

$$P(k) = \sum_{\Omega}\prod_{i\in\Omega} p_i \prod_{j\in\Omega^c}(1 - p_j) \tag{14}$$

where $\Omega$ includes all subsets of possible combinations of species rendering $k$ survivals and $\Omega^c$ its complement.

A more realistic scenario, however, is to admit that we can count or control the total abundance of the community, but not how it is distributed across species. In this context, would a small change on initial conditions drive the community towards a very different state? Because of species independence, the community transitions around an almost-continuum set of close-by states, without any large shift expected from small perturbations (Fig. 2). Changes in total biomass are continuous with and proportional to changes in species initial abundances. This is already indicative of the subsequent fingerprints that will be observed for so-called *local* MSS: the number and nature of states is directly proportional (and easily related) to the volume of initial abundances explored (Fig. 4 of the Main Text).

This picture of a non-interacting community is not of much interest for realistic scenarios, but it will prove a useful starting point for the following sections (see Fig. 2A in the main text). Some questions regarding multiple stability in communities made up of bistable species can be then reframed:

- How will different types of ecological interactions alter this highly-multistable landscape?
- Is it possible to find conditions by which the community can become bistable, hence allowing for full-scale shifts, and are these conditions expected in natural ecosystems?
- Can the same dynamical system exhibit different multistability regimes?
- Are different multistability regimes qualitatively equivalent across complex systems, or else they are very particular to specific parametrization schemes?

#### B. Bistable species, mutualistic interactions

##### 1. Dimensionality reduction under homogeneity constraints

Admit now a system with non-vanishing positive interaction terms $A_{ij} > 0$, so that $A_{ij}$ determines the mutualistic impact of species $j$ on species $i$.

$$\frac{dx_i}{dt} = x_i \left( \sum_{j=1}^{N} A_{ij} \frac{x_j}{\gamma_j + x_j} - d_i - B_i x_i \right) \tag{15}$$

Now each species improves its growth but also that of other species, each impacted by the abundance of $j$ via $A_{ij}$. As discussed in the main text (as the precise scenario where global bistability emerges following (Gao *et al.*, 2016)) and later in this Supporting Information, it not need be that a species improves its own growth for the dimensionality reduction to work: in fact, it is precise when a species alone is not bistable (see next section). This can be implemented by setting $A_{ii} = 0\ \forall i$. Now the presence or absence of one species due to the AE also depends on the rest of the community.

Recent advances have shown that the original system can be reduced to a 1-dimensional equation for the average species abundance provided $A_{ij} > 0\ \forall i, j$ (Gao *et al.*, 2016). Besides the assumption of purely mutualistic interactions, the dimensionality reduction technique also involves assuming that the interaction network has little degree correlation, so that the topological neighborhood of all nodes is independent of their degree or connectivity and so all species feel the same average external effect (Gao *et al.*, 2016). Finally, the approximation also needs node activities to be uniform when the dynamics are nonlinear, which can be obtained by assuming small variance in the dynamical parameters of Eq. (9).

Very briefly, the dimensionality reduction technique follows from averaging out both the global mutualism that each species perceives from the community, and the participation of each on this global mutualistic effect. If all other intra-species parameters $\gamma_i, d_i$ and $B_i$ are sufficiently similar (Gao *et al.*, 2016), the average or *effective* abundance $x_e$ of each species becomes

$$\frac{dx_e}{dt} = x_e \left( \alpha_e \frac{x_e}{\gamma + x_e} - d - B x_e \right) \tag{16}$$

with the average abundance obtained as an average over nearest-neighbor activity

$$x_e = \frac{1^T A \mathbf{x}}{1^T A 1} \tag{17}$$

and the effective mutualistic impact obtained from averaging the nearest-neighbor weighted degree

$$\alpha_e = \frac{1^T A \mathbf{s_{in}}}{1^T A 1} = \frac{1^T A^2 1}{1^T A 1} \tag{18}$$

These results imply a relevant scenario for understanding community-level regime shifts: the presence of bistable dynamics at the local can generate a perfectly bistable community, by the synchronization of all mutualistic effects into a single effective positive feedback loop of $x_e$ on itself (Kéfi *et al.*, 2016). Furthermore, if the conditions imposed by the mean-field process are fulfilled, we can compute straightforwardly both the system biomass (Fig. 3, top-left)

$$N \cdot x_e^{*,S} = \frac{N}{2B} \left( \alpha_e - B\gamma - d + \sqrt{(\alpha_e - B\gamma - d)^2 - 4B\gamma d} \right) \tag{19}$$

and the transition towards bistability at

$$\frac{1^T A^2 1}{1^T A 1} = \alpha_e > \alpha_e^c = B\gamma + d + 2\sqrt{B\gamma d} \tag{20}$$

These findings have harnessed attention in our search for the mechanisms by which complex ecological communities can display regime shifts: it appears as if such upscaling process, from local to global bistability scenarios, could be underlying the complexity of ecosystem resilience. In the sections below we discuss the arisal of different scenarios by which global bistability loses its universality and easily becomes blurred or disappears. Nevertheless, the technical limitations, extensions and applicability of the dimensionality reduction method (beyond the emergence of different state types) has been analyzed in depth in a variety of research works that have been very useful to our research. The reader might go to the following if interested in dimensionality reduction techniques in particular: (Arnoldi *et al.*, 2016a; Laurence *et al.*, 2019; Thibeault *et al.*, 2020; Tu *et al.*, 2021, 2017; Vegué *et al.*, 2023).

##### 2. Intertwining between global and local effects

One caveat of the above method that is relevant to our work is the possibility that many local stable states interfere with the global all-or-none picture. The idea here can be understood under the following reasoning: the dimensionality reduction technique predicts a system where, under low-enough mutualism, all species are extinct and $x_e^* = 0$. When mutualism increases and crosses $(1^T A^2 1)/(1^T A 1) = \alpha_e > \alpha_e^c = B\gamma + d + 2\sqrt{B\gamma d}$, synchronized cooperative effects imply that suddenly all species cross their Allee Effect and are able to coexist, generating the already-studied critical transition of figure 3top.

However, to assume that under low mutualism all species are extinct implies assuming that species are not bistable, and hence cannot survive, when alone. Then, the dimensionality reduction method in (Gao *et al.*, 2016) becomes precise (admitting homogeneity) by either setting $A_{ii} = 0 \; \forall i$, or at least making the diagonal elements follow the same statistics as the rest of the cooperation matrix. To us, this is a particular setting: we would be considering a system where intra-species effects are either null or equivalent to inter-species ones, whereas the tradition in theoretical ecology has been to consider self-regulation and inter-species interactions as two different dynamical entities (see e.g. (Puccia and Levins, 1985) or (Yodzis, 1981)). Moreover, to admit a system where all species need their mutualistic partners for survival is not necessarily true for all mutualistic communities: plant-pollinator couples are likely to require each other for survival (Memmott *et al.*, 2007) but other species can be bistable by themselves in isolated laboratory (Fronhofer *et al.*, 2015) or field (Nagel *et al.*, 2021) experiments.

Now, if species are inherently bistable, so that $A_i > A_i^c = B_i\gamma_i + d_i + 2\sqrt{B_i\gamma_i d_i}$ is fulfilled at least for some $i$, one would expect that a fraction of local states, with some degree of presence or absence, can be observed at low mutualism (Figure 2A in the Main Text). Furthermore, if $A_i$ is not exactly the same for all species, one would not expect that they overcome their Allee Effect in synchrony. This is indeed what we observe in simulations (Fig. 2A in the Main Text and figure 3 bottom): at very high mutualism species are either extinct or all coexist, but as we get closer to the $\alpha_e^c$ transition, some species that do not contribute much to global mutualism can survive without fostering the presence of all others. As mutualism increases, only the very weak cooperators can be present without fostering all others, until bistability becomes the rule (Fig. 3 bottom-center). This can also imply that the transition towards global coexistence happens for larger $\langle A_{ij} \rangle$ than expected if all species contributed equally strongly to others' survival.

The above considerations on species surviving independently are valid even for the relatively homogeneous species parameters constraints imposed by (Gao *et al.*, 2016). When heterogeneity in all non-interaction parameters ($\sigma_i i$ in section I, affecting $A_{ii}$, $d_i$, $\gamma_i$ and $B_{ii}$ but not $\langle A_{ij} \rangle$) kicks in, the survival of each species happens at very different threshold values (Eq. 3). This effect further dismantles the purely global scenario, increasing the regions when some species can survive with the others and vice versa (Fig. 4).

The above results are particularly important to our analysis as they show that, even under the assumption of purely mutualistic interactions between bistable species and a fully connected network, not all parameter choices will allow for the perfect bistable upscaling expected from (Gao *et al.*, 2016), and states with different species configurations might already coexist in this otherwise positive and homogeneous setting.

##### 3. Visualization in 2D systems

How can we visualize how local, independent-like states like those of Fig. 3 (bottom) interfere with the global all-or-none picture provided by dimensionality reduction techniques of (Gao *et al.*, 2016), as observed in Fig. 2A of the Main Text? A simple way to visualize what is happening is to recapitulate these global-local comparison a 2-dimensional system.

If bistable species do not interact, many local stable states can interfere with that global bistable picture (Fig. 2). Suppose two bistable species $x$ and $y$ following Eq. (1) under mutualism

$$\frac{dx}{dt} = x\left(A_{xx}\frac{x}{\gamma_x + x} + A_{yy}\frac{y}{\gamma_y + y} - d_x - B_x x\right) \tag{21}$$

$$\frac{dy}{dt} = y\left(A_{yy}\frac{y}{\gamma_y + y} + A_{xx}\frac{x}{\gamma_x + x} - d_y - B_y y\right) \tag{22}$$

If $x$ and $y$ are independent, there are two large initial conditions volumes $V_i$ (Fig. 5 red area), consistent with the basin of attraction of each species AE, where one species can be present without the other - hereby allowing for local *local* states (See Fig. 2B in the Main Text). These compositions will be observed when $x(t = 0)$ is large while $y(t = 0)$ is small or vice versa. As species start to cooperate, the presence of $x$ will alleviate the AE of $y$. The ic regions that allowed for local states will shrink, while the global coexistence basin will become larger. For strong cooperation, the system will only be at global coexistence or extinction states.

These 2D visualizations allow us to understand how mutualism shapes the possibility of local present-absent *local* states, that in turn disrupt whole-community bistability. Only strong mutualism eradicates these multiple states maintaining global bistability, but in a region where the global extinction basin also becomes small.

##### C. Bistable species, competitive interactions

###### 1. Failure of dimensionality reduction techniques

We have seen that in purely mutualistic communities, single-species bistability can upscale to the community level if cooperating species cannot survive when alone. In that setting, the system shows an all-or-none scenario arising from the single-species dynamics. What kind of local-to-global upscaling occurs in a community under competitive interactions?

Now the previous system allows for intraspecies cooperation only, while competition can happen between species.

$$\frac{dx_i}{dt} = x_i \left( A_i \frac{x_i}{\gamma_i + x_i} - d_i - \sum_{j=1}^{N} B_{ij} x_j \right) \tag{23}$$

The matrix elements $B_{ij} > 0$ determine the strength of competition between species $i$ and $j$ due to a multiplicity of factors such as an overlapping niche implying competition for space or resources (Roughgarden, 1983). Because self regulation at $B_{ii} > 0$ was already in place, single-species dynamics are identical to the above cases, with bistability between $x_i^* = 0$ and $x_i^{*,S}$. Following the previous results and under homogeneous-enough parameters, we could define a system for the abundance of an *effective* species

$$\frac{dx_e}{dt} = x_e \left( A \frac{x_e}{\gamma + x_e} - d - \beta_e x_e \right) \tag{24}$$

This encapsulates the notion that all species would suffer global competition as an average self-regulation effect with rate

$$\beta_e = \frac{1^T B \mathbf{s_{in}}}{1^T B 1} = \frac{1^T B^2 1}{1^T B 1}. \tag{25}$$

If all species suffered this same average effect, the community would be bistable, with total biomass $X$ stable at $X^* = 0$ and

$$X_T = N \sum_{i}^{N} x_e^{*,S} = \frac{N}{2\beta_e} \left( A - \beta_e \gamma - d + \sqrt{(A - \beta_e \gamma - d)^2 - 4\beta_e \gamma d} \right) \tag{26}$$

and a transition towards community extinction happening when competitive strength overcomes

$$\beta_e > \beta_e^c = \frac{1}{\gamma} \left( A + d - 2\sqrt{Ad} \right) \tag{27}$$

However, we know that competition between species doesn't decrease their abundances symmetrically (Arnoldi *et al.*, 2016a). In fact, as we will later see with two-species systems, one can understand how competition will eradicate the global coexistence state, but not make it transition to a global extinction, but to competitive exclusion states where one of the two species survives and symmetry and the *effective abundance* $x_e$ picture are broken. As known for Generalized Lotka-Volterra models (see section III.A and e.g. (Bunin, 2017)), the diversity of competitive communities is known to decay under increasing $\langle B_{ij} \rangle$ in a gradual process where weaker competitors go sequentially extinct (Fig. 6).

###### 2. Intertwining between exclusion and local bistability effects

In our model and those involving competition between bistable species, the gradual decay of diversity under increasing competition coexists with the presence of single-species Allee Effects. Because the $x_i^* = 0$ state is still stable for each species, the full system can be at the state with $S < N$ surviving species equivalent to competition between monostable species, but also in any of the states with lower diversity ($S' \geq S < N$) where some of the competing species are not present (Fig. 6 middle).

A community made up of competing bistable species, therefore, will not replicate a bistable all-or-none behavior. What we see is a combination of many local present-absent states, explained by independent species bistable dynamics (Section II.A), and the gradual extinction of weaker species due to competitive exclusion (Fig. 6, center). This phase of local multistability constrained by exclusion is likely to be connected to what is called *selective exclusion* in (Lee *et al.*, 2022). This simple setting provides a clear example of a scenario where bistable elements in a complex interacting network do not allow for a global bistable system.

Species parameters non-related to interactions are sorted from log-normal distributions with standard deviation $\sigma_{ii}$. In the case of pure mutualism, we saw how heterogeneity, by making single-species survival thresholds different, changes the qualitative behavior of the previously homogeneous system: symmetry becomes broken and not all species follow the same effective dynamics. The competitive case, however, is already non-symmetric for the homogeneous case: the system tends to survival of strongest competitors. The qualitative nature of this multistability regime is based on the presence of transitive competition. Differences in single-species parameters contribute to this transitive constraint, by reinforcing the competitive hierarchy. This is likely to make mutual exclusion scenarios easier to reach at lower competition, but does not dismantle the qualitative nature of local exclusion effects (Fig. 7).

##### 3. Mutual exclusion and community-wide extinction

In the limit of high inter-species competition we recover a well-known and thoroughly studied multistable scenario: mutual exclusion. Here, the system can be at $N$ different states, each containing one surviving species. However, the classic mutual exclusion regime of competitive Lotka-Volterra systems happens when inter-species competition is greater than intra-species self-regulation ($B_{ij} \geq B_{ii}$, see section III.A). Here we see how mutual exclusion happens at lower $\langle B_{ij} \rangle$ values: because the $x_i^* = 0$ state is already stable for each species, increasing competition makes survival of the best competitor harder. This is because competition reduces the volume of initial conditions that would allow a species to overcompete the others without falling into its own Allee Effect trap (Fig. 8, right).

When compared to competition between monostable species, the stability of $x_i^* = 0$ states ensures that, for strong enough competition, all species can become extinct and mutual exclusion becomes effectively impossible unless a competitor starts at much higher abundance than others (Fig. 3 of the Main Text, right panel, top-left corner, and Fig. 8), a high-dimensional result previously predicted (Wang *et al.*, 1999) and observed (Kramer *et al.*, 2009) in 1- and 2-species systems. When heterogeneous interactions come in place, we will see how this same effect precludes the emergence of chaotic dynamics and fluctuations around metastable states (Biroli *et al.*, 2018) (see section II.F).

##### 4. Visualization in 2D systems

To visualize how negative interactions disrupt the all-or-none bistable scenario into alternative and mutual exclusion states, we can study a 2D competitive system. Our aim is to visualize how inter-species competition alters the basin of the global coexistence state until it disappears in favor of one or another competitor.

Now the 2D system reads

$$\frac{dx}{dt} = x\left(A_x \frac{x}{\gamma_x + x} - d_x - B_{xx}x - B_{xy}y\right) \tag{28}$$

$$\frac{dy}{dt} = y\left(A_y \frac{y}{\gamma_y + y} - d_y - B_{yy}y - B_{yx}x\right) \tag{29}$$

The intermediate region with both competitive exclusion and intra-species Allee Effects (Fig. 9 middle) is particular to bistable species competition: in the $N > 2$ scenario, this configuration corresponds to local exclusion states if more species where present, with more than one possible coexisting community in place. For stronger competition strengths, the system moves towards mutual exclusion (Fig. 9 right).

##### D. A note on predator-prey interactions and oscillations

Predator-prey interactions, in which one species feeds (and hence, increases its growth rate) by killing or decreasing the growth rate of another, are widespread in nature and constitute the backbone of so-called food webs (Berryman, 1992; Pimm *et al.*, 1991). If our previously discussed system harbored such trophic interactions, and in the absence of other interaction types, one possibility would be to rewrite the original competitive matrix to account for predation

$$\frac{dx_i}{dt} = x_i \left( A_i \frac{x_i}{\gamma_i + x_i} - d_i - \sum_{j=1}^{N} P_{ij} x_j \right) \tag{30}$$

or else account for predation as a separate dynamical entity from competition or self-regulation, also considering that the dynamical response of predation could follow a saturating response as in (Qian and Akçay, 2020)

$$\frac{dx_i}{dt} = x_i \left( A_i \frac{x_i}{\gamma_i + x_i} - d_i - B_{ii} x_i + \sum_{j=1}^{N} P_{ij} \frac{x_j}{\theta_j + x_j} \right) \tag{31}$$

Here the matrix $P$ has a particular and well studied form, $P_{ij} = -\epsilon_{ij} P_{ji}$, with $\epsilon$ describing the so-called *assimilation efficiency* (Yodzis and Innes, 1992). The matrix $P$ then indicates that one species benefits from the $i - j$ interaction while the other finds it detrimental, and in the $\epsilon_{ij} \neq 1$ case this happens at different weights, often indicating that not all energy or body mass lost by the prey is perfectly gained by the predator (Yodzis and Innes, 1992). Because $\epsilon_{ij}$ might be particularly difficult to evaluate, another approximation involves assuming that $P_{ij}$ and $P_{ji}$ have opposite signs, but random absolute value meaning no further correlation between pairs of trophic rates as implemented here or in e.g. (Tang *et al.*, 2014).

Predator-prey community interactions, because of the particularity of this matrix containing the exact same amount of $+$ and $-$ off-diagonal signs, can generate novel dynamical patterns apart from stable attractors. The best well-known scenario is the appearance of periodic solutions, already seen in the classical $N = 2$ system (Lotka, 1925; Volterra, 1927). Periodic oscillations in ecology have been studied thoroughly for decades (see e.g. (Berryman, 1992; May, 1972a; Strogatz, 2018)). Here we briefly remind the reader how predator-prey matrices do not necessarily lead to oscillatory behavior, keeping the local exclusion states in place.

Our model, under the above version that includes predator-prey interactions but maintains the otherwise-accepted presence of self-regulatory dynamics (Barabás *et al.*, 2017), does not appear to show oscillatory behavior. This is true at least for a wide parameter range within $\langle P_{ij} \rangle \in [0, 0.2]$ and a sufficiently large community size of $N = 50$ (Fig. 10), as well as when admitting species mutualism within $\langle A_{ij} \rangle \in [0, 1.0]$ (not shown), consistent with the overall rarity of oscillations also found in a similar multilayer model (Qian and Akçay, 2020). The main reason for this is species self-regulation, that here takes the form of a logistic $-\beta_i x_i^2$ term (Eq. 24). Limitations to single-species uncontrolled outgrowth, as well as intraspecific variation (Allesina *et al.*, 2021), are processes known to preclude the presence of oscillations in favor of stable coexistence (Barabás *et al.*, 2017; Case and Casten, 1979). In our model, as already seen for the competitive scenario, this stable coexistence scenario is intertwined with local exclusion effects resulting from single-species bistability (Fig. 10 left). Increasing the mean absolute of trophic coupling decreases overall species abundance and community diversity while maintaining the multiple stable states already seen in the local exclusion domain. *Global* bistability in the presence of predation remains present at very small $\langle |P_{ij}| \rangle$ and higher mean $A$ (Fig. 10).

Furthermore, for Lotka-Volterra interactions with logistic growth and across the studied models (see section III), we observe that oscillations are hardly seen as long as self-regulation overcomes a given threshold and species-richness remains relatively high (possibly at $N > 20$), in which case chaotic fluctuations, as discussed along the main text and sections II.F.2 and II.G, dominate over periodicity. Studying the precise form of this threshold is beyond the scope of this work.

Finally, as hypothesized in the discussion of the main text, our work proposes that a missing ingredient in our understanding of catastrophic shifts in ecological communities is the role of network architecture, and how this can in turn modulate global bistability scenarios. There are few but very interesting theoretical models that have studied this possibility under particular network architectures and constraints (see (Lever *et al.*, 2014) and also (Karatayev *et al.*, 2020) for a recent analysis that is specific to food webs). Food webs in particular are known to exhibit non-random architectures (Allesina and Pascual, 2008; Berlow *et al.*, 2004; Dunne, 2006; Dunne *et al.*, 2002). However, our first step in the present research has been to analyze the role of mixed interaction types and random network structures in multistability. Therefore, we hypothesize that delving deeper into the role of predatory dynamics while considering random networks only instead of structured ones could generate theoretical artifacts. We leave this analysis as a necessary next step in our research and we refer the reader to a large body of literature on oscillations and stability on large predator-prey communities and food webs that is more suited for this topic (see e.g. (Akjouj and Najim, 2022; Barabás *et al.*, 2017; Galla, 2018; Polis and Strong, 1996; Rooney and McCann, 2012)).

##### E. Bistable species, mixed interaction types

So far we have seen how relatively homogeneous interactions can generate different multistability regimes. Mutualism can make species bistability upscale to generate global bistability but also intermediate present-absent configurations. On the other hand, competition disrupts community coexistence and allows for local exclusion states with less and less diversity as competitive strength increases, until mutual exclusion is the norm.

Most ecological communities, however, contain a mixture of both interaction types together with predation and other less-common interactions such as commensalism (Pilosof *et al.*, 2017). What type of multiple stable states can emerge then? Our goal here is to thoroughly explain, taking the original example model as a red thread, how different types of multistability can emerge from a single dynamical system, and how it is easy to understand the regime under which each does.

The complete system under study now reads:

$$\frac{dx_i}{dt} = x_i \left( \sum_{j=1}^{N} A_{ij} \frac{x_j}{\gamma_j + x_j} - d_i - \sum_{j=1}^{N} B_{ij} x_j \right) \tag{32}$$

If we simulate this system and count the number of stable states and their properties for a given set of homogeneous $A$ and $B$ matrices (here we constrain $\sigma_{A,B} \leq 0.1$), we obtain the homogeneous map of Figure 3 (Main Text), where we can see the emergence of global bistability, local multistability and mutual exclusion states.

###### 1. Effectively neutral interactions

One interesting aspect of Fig. 3 (Main Text) is the appearance of a wide yellow region, indicative of a local multistability domain. This means that, even for non-negligible interaction strengths $A, B > 0$, there appears to be a large domain where species recover their bistability and many community configurations can be possible.

Here we show that this region responds, in multilayer systems, to a domain where interactions are effectively weak or neutral due to a balanced tension between mutualism and cooperation. To do so in a simple way, we suppose here a limit-case scenario where interactive parameters of the system are purely homogeneous, relaxing the above $\sigma_{A,B} \leq 0.1$ to $\sigma_{A,B} \approx 0.0$. If so, our example model can be written

$$\frac{dx_i}{dt} = x_i \left( F_i(x_i) + \bar{A} \sum_{j \neq i}^{N-1} \frac{x_j}{\gamma + x_j} - \bar{B} \sum_{j \neq i}^{N-1} x_j \right) \tag{33}$$

where $\bar{A}, \bar{B}$ are now parameters and we have isolated self-regulatory dynamics in

$$F_i(x_i) = A_i \frac{x_i}{\gamma + x_i} - d_i - B_i x_i \tag{34}$$

If we are seeing all possible species combinations in the yellow region, it means that species are independently bistable, consistently with what will be later seen in section II.G. For this to be so in the purely homogeneous case we ask for inter-species interactions to cancel out, so

$$\bar{A} \sum_{j \neq i}^{N-1} \frac{x_j}{\gamma + x_j} - \bar{B} \sum_{j \neq i}^{N-1} x_j = 0 \tag{35}$$

But because competitive interactions disrupt species symmetry, here $x_j$ could take different values at equilibrium. One possibility is to focus on the precise transition where all species are present, but state multiplicity emerges due to competition canceling out the mutualistic effect (the purple-to-yellow, global-to-local states transition in Figure 3 of the Main Text). In this specific line, all species are still present and, under homogeneity of $F_i(x_i)$, their abundance can be computed from Eq. (2) and does no longer depend on $i$, $x_i^{*,S} = \omega \neq \omega_i$. The calculation then simplifies to

$$\bar{A} \sum_{j \neq i}^{N-1} \frac{\omega}{\gamma + \omega} - \bar{B} \sum_{j \neq i}^{N-1} \omega = 0 \tag{36}$$

$$\bar{A}(N-1)\frac{\omega}{\gamma+\omega} - \bar{B}(N-1)\omega = 0 \tag{37}$$

from where one can learn, for a given $\omega$ and $\gamma$, the given $A/B$ ratio where interactions will cancel out:

$$\frac{\bar{A}}{\bar{B}} = \gamma + \omega \tag{38}$$

We can plot this relationship and compare it with a purely homogeneous multistability map. We can see how $B(A) = (\gamma+\omega)^{-1}A$ sits right at the transition where local multistability comes in place (Fig. 11). Computing effectively neutral domains when some heterogeneity kicks in is likely to become a more complex endeavor, but still the homogeneous case is an illustrative toy model for the emergence of independent-like effects in multilayer systems.

This result and its potential extensions across multilayer ecological systems might also be relevant to discussions on the concept of ecological communities as *loose collections of species*, where the presence and absence of a species might not necessarily affect the others (see e.g (Clements, 1936; Liautaud *et al.*, 2019) or the Discussion section of the Main Text). The possibility that this is not only happening between non-interacting species, but also along effectively balanced interaction schemes widens the possible scenarios where *loose collections* effects are at play.

##### 2. Relaxation times and fluctuations at the mutual exclusion threshold

The progression from global bistability to mutual exclusion is continuous for $B_{ij} < B_{ii}$, giving rise to a highly-multistable region with local states, but is sharp and abrupt at $B_{ij} \geq B_{ii}$, as seen in figure 12. This is easy to understand in analogy to the generalized Lotka-Volterra model. In this model, the community is at a single state with less and less surviving species for $B_{ij} < B_{ii}$, until it undergoes a transition to mutual exclusion once $B_{ij} \geq B_{ii}$ (see section III as well as (Kessler and Shnerb, 2015) for a detailed description of this critical point). What is new here is that, because species are locally bistable and can by themselves be present or absent, the $B_{ij} < B_{ii}$ domain can include a multiplicity of states (at least 2, global bistability), as seen in the above sections. However, as mutualism becomes strong ($A_{ij} \approx NA_{ii}$), species no longer perceive local Allee Effects, therefore they become effectively monostable and behave as in the GLV model. Here, the community jumps from global bistability towards mutual exclusion without allowing individual effects to emerge (Fig. 12A).

Interestingly, as later discussed in the Fingerprints section (II.G), species abundances around the $B_{ij} = B_{ii}$ threshold can take a long time to stabilize, a natural characteristic of phase transitions and criticality (Solé, 2011). This implies that, even for very homogeneous communities in the laboratory, where cliques are unlikely to emerge, there might be a very narrow window of slowly-relaxing or even seemingly fluctuating states close to the critical border between the global bistability and mutual exclusion phases.

##### F. Bistable species, heterogeneous interactions

The strengths of species interactions, and hence the matrices we use to describe them, are likely to be far from homogeneous in nature. For simplicity, here we modulate heterogeneity in species interaction strengths via the standard deviation $\sigma_{ij}$ of the sorted off-diagonal entries of $A$ and $B$ matrices, a common procedure in community ecology (May, 1972b). By doing so, we observe that a new phase appears, consistent with similar multistability scenarios observed in well-studied generalized Lotka-Volterra systems (see below).

With an aim on simple statements, here we study several aspects of this phase. First, we study how clique states appear as we increase heterogeneity, to find that the heterogeneity threshold $\sigma^c$ is not sharp nor uniform for small- to medium-sized communities. Second, we explore the stability and fluctuations of states in this phase and link it to a vast body of recent literature. Finally, we study how clique states also emerge under homogeneous interaction strengths if the connectivity in the Erdős-Rényi graph is modulated.

The term *clique* as a label used for subgroups of coexisting species was introduced to our knowledge in (Fried *et al.*, 2016). There, it arises from its use in the mathematical field of graph theory, where a clique is a group $G$ of vertices in an undirected graph, so that each possible pair of vertices in $G$ is connected (Alba, 1973). Under the strong-or-null competition assumption of (Fried *et al.*, 2016), the community matrix becomes a coexistence graph, and its cliques are well-separated groups of coexisting species. In our work, subgroups of coexistence species are likely to belong to cliques of a possible coexistence graph. However, we have not studied this in detail and hence cannot claim whether they are necessarily maximum cliques (Pardalos and Xue, 1994), although the fact that they are tested against migrations gives a hint in that direction.

Instead, we recover the original meaning of the term in the social sciences (later applied to graph theory in (Luce and Perry, 1949)): a clique is a group or collective of people interacting together via shared interests, often with a well-defined boundary of who belongs to the group (Mokken *et al.*, 1979). It is in this wider semantics that we think the term *clique* is a useful label to describe those multiple stable states, composed by small groups of species, that are separated by sharp boundaries instead of gradual species changes.

###### 1. The heterogeneity threshold and the emergence of cliques

Analytical work studying ecological models with tools from statistical mechanics and spin-glass systems has predicted different flavors and types of a transition similar to the one discussed here and in the Main Text. This transition, from monostability towards a phase with different types of multiple attractors, was first glimpsed within the *complexity-stability* debate (McCann, 2000): Robert May predicted a complexity limit to the stability of a species-rich ecosystem (May, 1972b), with complexity here indicative of both interaction strength heterogeneity and connectivity. As interactions become heterogeneous, the system might cross a sharp boundary beyond which it is likely to become unstable, as latter understood for more specific ecosystem models (Yoshino *et al.*, 2007).

Later research has deepened our understanding of this transition and, more importantly, what lies beyond it. In the specific field of theoretical community ecology, research on a family of Generalized Lotka-Volterra ecosystems (GLV, see section III.A) has uncovered the presence of a domain with many attractors beyond this heterogeneity threshold (Bunin, 2017; Diederich and Opper, 1989; Fyodorov and Khoruzhenko, 2016; Galla, 2018). Moreover, recent work has highlighted the universal nature of this transition across single- and multi-layered complex systems (Ipsen and Forrester, 2018).

As discussed in the Main Text, a simple conceptual understanding for ecologists of this high-dimensional phenomena lies in understanding the concept of *intransitive* competition. Now, by disrupting the possibility of a linear order of what species outcompetes the other, intransitive competition creates a complex set of rock-paper-scissors motifs, likely to break down the possibility of a single stable state of species coexistence.

For the GLV model in particular, and following the analysis by Guy Bunin (Bunin, 2017) for very large and weakly interacting ecosystems but implementing our notation, one would find that the threshold towards the multiple states phase happens for

$$\sqrt{N}\sigma^c = \frac{\sqrt{2}}{1+\theta} \tag{39}$$

where $\theta$ stands for the correlation between random interaction strengths, $\theta_B = \mathrm{corr}(B_{ij}, B_{ji})$, which is by definition equal to $\theta_A$ in our system (see section I). Because our interaction matrices $A$ and $B$ are not correlated ($\theta = 1$, symmetric interactions) nor anticorrelated ($\theta = -1$, antisymmetric interactions), $\theta = 0$ and one would expect the GLV version of our system to shift towards the cliques phase at $\sigma^c = \sqrt{2/N}$, at least for large $N$.

Although this exact prediction was not proposed for a system involving non-linear functional responses, we are here interested in comparing it to the studied scenario. Do we see such a transition? and more importantly, is it sharp and uniform across all the phase space for small- to medium-sized communities of $N \sim 10 - 50$?

This would mean that, beyond this $\sigma^c$ value, all homogeneous multistability families disappear into the clique phase. As stated in the Main Text, what we discuss here is how the transition is neither uniform nor sharp for communities of moderate size.

- As discussed above, theoretical models postulate the existence of a single heterogeneity threshold, beyond which communities are likely to loose their fixed point stability in favor of a more complex multistable phase (see the very recent article (Mallmin *et al.*, 2023) for a consideration of how the heterogeneity threshold depends on mean interaction strength). Here we see that the value that this threshold takes depends strongly on the average interaction strengths in the community. The critical value $\sigma^c \sim \sqrt{1/N}$ is a good estimate for the heterogeneity beyond which cliques start to emerge (Fig. 13), also tested for communities of $N = 10$ and $N = 20$ species (not shown). However, this is only so in systems with weak interaction strengths. When either cooperation or competition dominate the system, cliques emerge only at very high heterogeneity, or might not even appear in favor of the global bistability or the mutual exclusion phases, respectively (Fig. 13).

- As previously intuited and discussed by Guy Bunin (see Figure 5B in (Bunin, 2017)) and later works (Mallmin *et al.*, 2023; Sidhom and Galla, 2020), even for relatively large communities of $N = 100$, only a fraction of the simulated GLV systems showed multiple states at $\sigma \approx \sigma^c$. This means that the transition might be far from sharp for even smaller communities, and stable states can be intertwined with those of other categories as well as chaotic attractors (see e.g. (Mallmin *et al.*, 2023)). In figure 13 we can also observe the nature of this phase boundary for a moderate community size of $N = 50$. Although regions are relatively defined, we can see that borders and not sharp, and for example globally bistable scenarios can emerge close and beyond phase boundaries (Fig. 13).

These results show that there are certain considerations regarding the sharp transition to a multistable behavior when dealing with medium-sized communities: the $\sigma^c$ value beyond which cliques dominate the system depends on the sign and strength of species interactions, and even then, the transitions are not sharp and different MSS categories can coexist close to this boundary.

##### 2. Stability of cliques and chaotic fluctuations

A large body of recent research is uncovering relevant properties of the multistable phase, depending on the precise definition of the model, the source of heterogeneity or the intrinsic properties of its random interactions. The main goal of our work, as is probably clear to the reader by now, is not to provide analytically advanced calculations on complex system multistability. Rather, we aim at trying to translate theoretical results into concepts that can be of use to the ecology readership.

As a means to refer the reader to this body of research, below we present a table with a body of articles that explore emerging multistability in communities under heterogeneous competition. We briefly mention their different findings related to the questions that we believe more interesting to multiple stable states in ecology: how the number of different states scales with $N$ (see section II.G.2), and whether at least some of these states are stable or not (Table I).

The main challenge in characterizing the properties of the multiple equilibria phase with analytical tools from physics is to go beyond the totally correlated and anticorrelated interaction regimes, $B_{ij} = B_{ji}$ and $B_{ij} = -B_{ji}$ (Ros *et al.*, 2022). In ecology, unfortunately, we expect complex communities to deviate from this perfect matrix architectures, and only very recent models are uncovering what can happen for the intermediate regimes where interactions are not symmetric nor anti-symmetric (Mallmin *et al.*, 2023; Ros *et al.*, 2022).

The main learning that we can get from table I is that complex and disordered communities with heterogeneous competition are likely to harbor a very large (often exponential in $N$) number of different species compositions. Empirical evidence is not easy to reconcile with such a notion of thousands or millions of alternative species compositions in communities of tens to hundreds of species. Moreover, we are currently learning that not all these species compositions are in fact stable states: whether they are or not depends strongly on the correlation between interaction strengths $\theta_B = \mathrm{corr}(B_{ij}, B_{ji})$, or, in the case where some species exclude each other while many others do not compete, on the inter- vs intra-species competition ratio $B_{ij}/B_{ii}$ (Bunin, 2017, 2021; Fried *et al.*, 2016).

Some of the most recent results for the standard GLV model indicate that, if interactions are asymmetrical and uncorrelated, as we have assumed throughout our model and is expected to be common in ecology, the system is likely to display unstable states and, in the presence of migration, engender a chaotic regime (Mallmin *et al.*, 2023). This is particularly interesting within the context of persistent temporal variability in ecological communities, with relevant properties of the chaotic phase being uncovered as this Supporting Information is being written (Benincà *et al.*, 2015; Hu *et al.*, 2022; Mallmin *et al.*, 2023; de Pirey and Bunin, 2023; Roy *et al.*, 2020).

However, the ecological scenarios that we are interested in our research, with communities being persistent and potentially undergoing large and abrupt shifts between stable states, do not seem to totally coincide with the notion of chaotic dynamics. In communities of moderate size (much below the $N \to \infty$ limit of analytical predictions), are all species compositions unstable? And are the chaotic fluctuations of the unstable scenarios persistent in time?

- As discussed in the Main Text, one of the relevant results of our work is to pinpoint that not all cliques appear unstable. Although it is challenging to compute analytically, we find that the fraction of stable cliques depends on the size of the community $N$ as well as on specific choice of the model and the interaction scheme $(A, B, \sigma)$ involved (Fig. 14). Recent findings on asymmetric interactions present differences regarding the stability of states within the cliques domain (Biroli *et al.*, 2018; Ros *et al.*, 2022). For example, simulations for moderate $N$ and asymmetric interactions already pointed out that not all cliques emerging from a single community are necessarily unstable (see e.g. (Bunin, 2017; Kessler and Shnerb, 2015; Sidhom and Galla, 2020)). Models assuming strong but sparse competition matrices also find that stable cliques can emerge systematically even in the many-species limit (Bunin, 2021; Fried *et al.*, 2016, 2017). It appears that this strong-interaction scheme might be key in explaining how chaotic attractors and stable cliques are related to interaction strength and heterogeneity (Mallmin *et al.*, 2023). These results and our simulations (Fig. 14) help reconcile the patterns of intransitive competition with the possibility of multiple states that are stable in time.

- Interestingly, as previously analyzed in (Roy *et al.*, 2020, 2019) and earlier explored in (Kessler and Shnerb, 2015), cycles or chaos can emerge intrinsically from heterogeneous interactions, but their persistence through time depends on the presence of external sources of noise or, as in our case or that of metacommunity (Roy *et al.*, 2020, 2019) or island biogeography (Kessler and Shnerb, 2015) models, migration. Because species are not allowed to go totally extinct (see Section I), there is a constant influx of migration back into the community. It is in this context that chaos will be persistent: dynamics will approach an unstable clique but will bounce back to another in a kind of *pinball*-like effect. In an ecological setting where species are not allowed to re-invade, likely depicting isolated environments, what we see is that chaotic regimes tend to stabilize, consistent with simulations in (Sidhom and Galla, 2020). As seen in figure 15, a proportion of states is stable, and the opposite fraction of cliques that are variable in time decays much faster if no migration is allowed. Nevertheless, the decay time of these chaotic regimes has been found to be extremely large in specific settings, meaning they are to be taken into account even under an $m_i = 0$ regime (see (Stern *et al.*, 2014) and section III.D). We refer the reader to two very recent articles on similar properties of the chaotic phase in the context of the Generalized Lotka-Volterra model (Mallmin *et al.*, 2023; de Pirey and Bunin, 2023)

| **Reference** | **Model** | **Source of het.** | **Symmetry of $B$** | **Stability** | **$\Gamma \sim N$ scaling** |
|---|---|---|---|---|---|
| (Diederich and Opper, 1989) | Replicators | Random $B_{ij}$ | Asymm. | Metastable | Upper bound $\Omega \sim e^{\lambda N}$ |
| (Kessler and Shnerb, 2015) | GLV, migr. and noise | Competitive $B_{ij} < 0$ | Random | Disordered (unst. if migr.) and Glass-like (stable) phases | Subexp. for the glass phase |
| (Fyodorov and Khoruzhenko, 2016) | Non-linear replicators | Random $B_{ij}$ | Antisymm., conserv. field | Few states are stable | Exp. $\Omega \sim e^{\theta N}$ |
| (Fried *et al.*, 2016) | Gilbert graph | Binary ($B_{ij}$ =strong or $B_{ij} = 0$) | Both | Stable cliques | Subexp. $\Omega \sim e^{\gamma \ln^2(N)}$ |
| (Bunin, 2017) | GLV | Random $B_{ij}$ (weak) | Asymm. | Stable, unstable | - |
| (Fried *et al.*, 2017) | GLV, E-R graph | Binary (Strong or null $B_{ij}$) | Both | Stable cliques | Subexp. to sublinear if asymm. network |
| (Biroli *et al.*, 2018) | GLV, noise | Random $B_{ij}$ | Symm. | Marginal phase | - |
| (Roy *et al.*, 2019) | GLV | Random $B_{ij}$ | Both | Chaos and aging if $m_i = 0$ | - |
| (Sidhom and Galla, 2020) | GLV, nonlinear feedback | Random $B_{ij}$ | Different $B_{ij}, B_{ji}$ correlations | Multistab. and chaotic phases | Exponential |
| (Altieri *et al.*, 2021) | GLV, noise | Random $B_{ij}$ | Symm. | Marginal phase | Exponential |
| (Bunin, 2021) | GLV, $m_i$, noise | Binary $B_{ij} > 0$ or $B_{ij} = 0$ | Asymm. | Hierarchically stable | Exponential |
| (Fedeli *et al.*, 2021) | Nonlinear extension of (May, 1972b) | Random $J_{ij} > 0$ | Both | - | Exponential or bounded |
| (Altieri and Biroli, 2022) | GLV, noise and intra-spp AE | Random $B_{ij}$ | Symm. | Hierarchical and stable phases | Sub-exponential, spin-glass phase |
| (Ros *et al.*, 2022) | GLV | Random $B_{ij}$ | Uncorrelated (asymm.) | Unstable states (chaotic regime) | typical $\Omega \ll \langle\Omega\rangle$ |
| (Marcus *et al.*, 2022) | GLV | Sparse $B_{ij}$ | Symm. | Multiple eq. | - |
| (Ros *et al.*, 2023) | GLV | Random $B_{ij}$ | Asymm. | Rare stable states if correlations$> 0$. | typical $\Omega \ll \langle\Omega\rangle$ |
| (Mallmin *et al.*, 2023) | GLV | Random but strong $B_{ij}$ | Asymm. | Chaos and stable states | - |

TABLE I Overview of research analyzing multistability in species-rich ecological communities. Different approaches, often through the analysis of the GLV model, have been interested in uncovering properties of the emerging cliques domain. Here we provide a brief overview of the type of **model** studied, the way **heterogeneity** is introduced in the model, the symmetries in the elements of the studied **interaction matrix** $B$ and whether the emerging states are found to be **stable** and how their **number scales with system size**.

##### 3. Heterogeneity at the network structure level

So far, our approach towards generating heterogeneous interactions has been by modulating the standard deviation of the sorted $A_{ii}$ and $B_{ii}$ parameters. In this context, we have chosen to use a totally connected network ($p \approx 1$ in the Erdős-Rényi graph) to ensure homogeneity and focus on the role of species interactions, also inspired by different research highlighting the importance of interactions against network structure when trying to predict collective community dynamics (Barbier *et al.*, 2018; Gao *et al.*, 2016). Other research, however, has obtained a glimpse on the possibility that network structures might indeed play a role on the way species feedbacks modulate collective multistability (Lever *et al.*, 2014).

Here we do not yet study in depth the role of non-trivial network structures. We will, however, study the effects of displacing from the fully connected network into a classical community model where not all nodes interact. The potential for community models with sparse interactions showing potential for nontrivial behavior has been discussed for example in (Fried *et al.*, 2016) or (Marcus *et al.*, 2022). As done in the original work my Robert May (May, 1972b) and now a milestone in high-dimensional community ecology, studying the role of $\sigma$ has often been accompanied by studying the role of interspecies connectivity with a Erdős-Rényi graph model (Erdős *et al.*, 1960). A useful extreme version of the model, opposite to our approach so far, is to consider that species either do not interact ($A_{ij}$=0, coexistence) or interact strongly but homogeneously ($A_{ij} = a \gg 0$, leading to mutual exclusion) (Bunin, 2021; Fried *et al.*, 2016, 2017). This has been usually implemented by defining a probability $p$ that two nodes are connected (see section I.C.4). Interestingly to our analysis, it is in this precise scenario of strong, homogeneous but sparse competition that cliques are by definition stable even under the many-species limit (see section III.A). We will show here the results of simulating our model with $p < 1$ and how these are related to increasing the standard deviation of parameter distributions.

The reason behind this is particularly simple. Imagine an interaction network where we set, with probability $p$, $A_{ij} > 0$ (see section I.C), and with probability $(1-p)$ $A_{ij} = 0$. We do the same for matrix $B$. We want to measure how the value of $p$ modulates the standard deviation of the matrix. We take an approximation for very low heterogeneity in $A_{ij}$ values (hence all $A_{ij}$ are either 0 or $a$) and the diagonal elements are not considered for the computation of network heterogeneity. There are, in the mean field approximation, $N(N-1)p$ off-diagonal $a$ values in the matrix, and $N(N-1)(1-p)$ zero values. The standard deviation then follows

$$\sigma = \sqrt{\langle X^2\rangle - \langle X\rangle^2} = \sqrt{\frac{N^2(N-1)^2pa^2}{N^2(N-1)^2} - \frac{N^4(N-1)^4p^2a^2}{N^4(N-1)^4}} = a\sqrt{p(1-p)} \tag{40}$$

as it can be easily observed in figure 16. Interestingly, this no longer depends on $N$, meaning that whatever the size of the system, we can only reach a maximum heterogeneity value of $\sigma = a/2$ if we only modulate connectivity but not parameter heterogeneity. This is why we have chosen to focus on heterogeneity through increasing the standard deviation of parameter distributions and not network connectivity in the Main Text. As expected, by setting $p = 1/2$ but homogeneous $a$ and $b$ values, the multistability phase space of our model is similar to that of Figure 3 (bottom) in the Main Text: because competitive heterogeneity kicks in, now in the form of non-uniform connectivity, clique states and fluctuations emerge (Fig. 17).

#### G. Fingerprints of the four multistability families

As discussed in the Main Text, we study here how basic properties of MSS correlate, in the search for patterns that are specific signatures of each multistability type and could be tested against experimental community patterns after varying initial conditions (?). Although other metrics where originally explored (Jaccard distance between states after a perturbation, necessary perturbation width to generate a state shift, etc., not shown), we found that the most basic properties (and hence those that are conceivably easier to measure in the lab) might already hold significant information. In particular we to study the number of stable states and their diversity, and how these respond to changes in system size or random abundance perturbations. As a third dimension, the potential for persistent abundance fluctuations was also found key in characterizing the clique fingerprint (see Figure 4 in the Main Text).

Below we discuss (1) the number of observed stable states $\Omega$ for each MSS type and how it scales with system size $N$ (Figure 4B in the Main Text), (2) the diversity $D$ of these states and the proportion of species overlap $O$ between them (Figures 4A and 4D) and (3) the relation between the number of stable states in a system and their probability of recovery after a perturbation.

##### 1. Number of states and system size

Being able to count the number of stable states in a complex energy landscape is a fundamental problem across research areas, with spin glass physics being a prominent example (Ros *et al.*, 2019b; Toulouse *et al.*, 1987). To frame this within our work, we here aim at analyzing the shape of $\Omega(N)$: the number of observed fixed points as we increase the size of the system. In ecology, as discussed throught the present work, this accounts for describing the number of possible states a community can be at given the size of the initial species pool $N$ and migration $m$ (see e.g. (Ros *et al.*, 2022) for a latest contribution). Here, an interesting example from an apparently alternative field could be Kauffman's $NK$ model (Kauffman *et al.*, 1993). In it, the search for understanding how many fitness peaks would the system hold as more units $N$ or more neighbor connections $K$ are considered granted interesting results and a glimpse on how even very simple, Boolean networks could engender an exponential amount of stable system configurations (Weinberger, 1991).

Because we do not use advanced analytical techniques such as the Kac-Rice method, which we deem extremely powerful but difficult to apply beyond strong model assumptions (Ros *et al.*, 2019b, 2022), the task of counting the *total* number of stable states for a specific multistability regime becomes particularly difficult for us. This is because of the following limitations:

- **Computational cost:** Doing a survey for the number of possible stable states implies sampling a sufficiently large amount of initial conditions. For local multistability scenarios, for example, and because we know there are as many as $2^N$ different states, one would need at least $10^{15}$ different initial conditions to observe all possible compositions of a 50 species system. We need to start from the premise that, as in the field and across experimental tests, the number of replicas will always be smaller, and not all states will ever be realized in such cases. Moreover, as discussed in sections II.E.2 and II.F.2 and in figure 15, some regimes such as cliques or critical domains of the phase space need a very long time to stabilize to a fixed state. To increase the probability of making sure that we are seeing a fixed state instead of transient fluctuations, we need very long simulations (Fig. 15). Thus, the overall problem implies sampling a very large amount of random initial conditions and integrating them for very long time windows. Because of this, computational techniques are likely to provide only a hint on the number of stable states of a system. However, this is possibly the same type of scenario we would face when evaluating multistability in the lab.
- **Contamination across multistability regimes:** We are here interested in knowing how many stable states each specific regime has. This would be done by assigning a typical $\Omega(N)$ scaling to each region in figure 3 of the main text. However, as discussed in section II.F.1, MSS regimes are not separated by sharp boundaries in the phase space, and multistability scenarios can be intertwined throughout space. Here we choose to generate random systems that appear to fall inside each of the four domains. However, it is likely that each regime still contaminates the others: evaluation for the number of cliques close to the global multistability scenario can appear to show fewer states than expected, etc. This highlights again the role of systems with moderate number of species ($N < 100$ as a rule of thumb) that appear to blurr out the otherwise sharp scenarios of the ideal $N \to \infty$ domain.

This said, what can we say about the number of observed states at each MSS domain? As discussed throught the first subsections of section II, the three MSS categories emerging under relatively homogeneous interactions have a straightforward scaling with system size. Global bistability is likely to remain constant with system size, so that $\Omega(N) = 2$: only the all-or-nothing states should be present (Fig. 4B, purple in the main text). Under strong competition, mutual exclusion states scale as $\Omega(N) \approx N+1$: one would see the extinction state together with the $N$ states, each with one surviving species. As intra-species parameters become heterogeneous, and hence competition remains transitive but with a stronger hierarchy, the states with weaker competitors will have a smaller basin of attraction and become harder to find, likely leading to a $\Omega(N) \lesssim N+1$ scaling. Finally, the scaling of local multistability states depends on the strength of transitive competition and single-species heterogeneity (see sections II.B and II.C). In the limit-case scenario of perfectly balanced or null interactions, all species can be present or absent individually, and the number of stable states can be as large as $\Omega(N) \approx 2^N$ bound (Fig. 4A, yellow in the main text). As discussed in the main text, this is an unfeasible number to explore in many settings. The fingerprint here would not be that so many states can appear, but rather than every possible initial condition likely leads to a different final species composition.

When competition becomes heterogeneous, a large amount of states seems to emerge. Being able to count or estimate the number of states in the clique regime across ecological settings is a sought-after open problem in community ecology, as seen in Table 1. Because of the complexity of the problem due to the 2 points stated above, here our aim is only to provide a partial computational contribution to the problem. The main goal is still the description of different multistability domains and their flavors in general terms, so that results are of general use to the ecology and interdisciplinary readers interested in complex systems. More importantly, the number of states and the types of (in)stability involved in the cliques regime can depend on system particularities, with the degree of symmetry of interactions being a potentially key element (see Table 1).

However, it is still interesting to study $\Omega(N)$ of cliques across the different models we are dealing with. This is because our approach, even though limited by the computational cost of generating many systems or the imprecision of evaluating stability, is likely to be closer to empirical setups, where only a given number of replicas or a constrained region of initial conditions can be studied: What we study is the number of states that can be *observed* after $s$ simulations and that are *stable* in time and against species migrations.

As seen in figure 18, our results seem to point to the number of cliques being in general larger than the number of available species $N$, with a potential bound in $\Omega(N) \sim N^\gamma$, $1 < \gamma < 2$ that is replicated across most of the studied models (see below). Although this number is much smaller than the exponential number of states seen for example in (Fyodorov and Khoruzhenko, 2016) and closer to subexponential scenarios discussed for example in (Fried *et al.*, 2016), it can still be a very large number: communities with $N = 50$ species could already be in hundreds of different stable states ($\Omega(N = 50, \gamma = 1.5) \approx 353$).

This allows us to state that, at least for communities with a moderate number of species, intransitive competition creates a landscape where potentially many stable states can be present. Even if $\Omega(N) \lesssim N^2$ does not seem an impressive upper bound and is way far from exponential, it is still much beyond the low-dimensional alternative stable states kind of picture where a system can be at – and switch between – two or few stable states. It is because of this that we are able to link intransitive competition to the emergence of many possible contrasting species compositions.

Beyond the *many cliques* flavour, there are further points we can make by studying the results of figure 18.

1. The presence of local multistability (stemming from 1-D bistability, see fig. 21) is likely to increase the expected number of clique states. We can observe how the GLV model, where species are monostable when alone (see section III.A), appears to have less stable cliques than others for small $N$, with the number increasing fast in a scaling closer to that of (Fried *et al.*, 2016) although not exact (not shown).

2. As $N$ increases towards $N = 50$, $\Omega$ seems to reduce the gradient of increase for different models. It appears as if a larger fraction of initial conditions start to become unstable. We hypothesize this can be due to large $N$ increasing the necessary time to stabilization beyond what we are capable of measuring (Fig. 15), or also because, as system size increases, the chaotic regime starts to take over and less stable states can actually be found (Ros *et al.*, 2022).

###### 2. Diversity of states and species overlap

As previously discussed, we have chosen to study diversity $D$, the number of surviving species in a given state, as the simplest community-level attribute that can be a proxy to community function (Thompson *et al.*, 2012), at least within the disordered systems perspective where species are not given specific attributes nor trophic hierarchies etc (Barbier and Arnoldi, 2017). It is therefore interesting for us to understand what is the possible diversity that different stable states can take. Here, the question will also be relevant to convey the following idea: there are four different multistability categories, but these do not cover all the space of possible MSS, but rather constrain it. One could imagine that all possible scenarios can sit in at least one of the four MSS categories: many states with few species, many states with large differences in species number, few states with equivalent number of species... etc. Studying state diversity allows us to see that this does not seem so.

The first simple way to explore the number of species that MSS can take is, taking advantage of the simulations of the previous section, to plot the scaling relation $D(N)$: what is the diversity of the final stable state, provided the total pool of species increases? As discussed in the fingerprints section and throughout the whole main text, the results are as expected and plotted in figure 19. The global bistability regime contains two possible states (Fig. 19 purple): a state containing all species, falling on the line $D = N$, and a state with no surviving species $D = 0$. However, the second is not easily attained: because we are generating random initial conditions at high mutualism, the basin of attraction of the zero state is particularly small and it would need all species to have very small biomass at the same time. At the other extreme, mutual exclusion states contain only one surviving species (Fig. 19 blue).

The question is then whether cliques and local multistability states span all the possible configurations. For the latter (Fig. 19 yellow), we have discussed in the main text and in section II.A that, in fact, local multistability can take any possible diversity value. Moreover, it will only depend on the initial conditions. In figure 19 we see that, because initial conditions are random but centered around the average Allee Effect threshold, the average number of surviving species is $\langle D \rangle \approx N/2$. However, local multistability is only a narrow domain in the space of possible MSS regimes: it needs that interactions are particularly week and balanced out (see section II.E.1).

Finally, the number of surviving species in cliques is of particular interest, as discussed throughout the literature of Table 1. The result in figure 19 (green) is relevant to us: as the system size increases, the number of surviving species in clique states will not increase much with it, if at all. Also, as shown in Figure 2C in the main text, cliques don't appear to have very different sizes. We hypothesize that this has a direct link with what has been discussed in the previous section: as $N$ increases, the fraction of initial conditions that result in chaotic fluctuations will increase (always when admitting a source of migration, see section I, or noise). In this context, the amount of species that can coexist under intransitive competition is bounded, possibly due to a similar reasoning as that of May's complexity limit (May, 1972b): the number of surviving species in cliques will decrease until those that coexist harbor a set of interaction strengths whose heterogeneity is low enough for them to coexist. Testing the interaction strength structure of clique species would provide interesting information, in a similar approach as that for the structure of single-state coexistence in (Barbier *et al.*, 2021).

To further explore the flavor of state diversity, we also include another simple metric: because we perturb stable states and observe the state they reach (see section I.E), we can measure the amount of species overlap $O$ between these two states. That is, $O = 0$ means the two states do not share any species, and $O = D$ means they share all species. Of course, if $O = D = N$ the two states are the same (total coexistence). Overall, the relation between the diversity of a state $D$ and the amount of species that are also found in the perturbed state $O$ informs us whether the two states are proportionally very similar or different [1]. Here we aim at depicting how the types of states that can emerge for a given community have specific constrains in their diversity and species overlap.

In figure 20 we plot, for a gradient from strong cooperation to strong competition ($A$-10$B$, where 10 is introduced for visualization to account for the original difference between $A_{ii}$ and $B_{ii}$, or simply $A_{ij}$ for the single-layer GRN model), the final diversity of states $D$ and the species overlap between states and their perturbed counterpart $O$. Together with this, we also observe how this species overlap value $O$ correlates with the original number of species $D$. The dashed line in figure 20 right panels is the $O = D$ line: the perturbation brought the system to a state with total overlap. Deviations below this line indicate that not all species in the original state where found in the final state. Together with the model of the Main Text (Fig. 20A), we also include the fingerprints for the Gene Regulatory Network model of section III.C (Fig. 20B) as we believe it helps in understanding the results.

Global bistability states obviously appear under dominating mutualism and involve $D = N$ states (Fig. 20 top left, purple). The basin stability of these states is particularly large (see e.g. section II.B.2), and the extinction state $D = 0$ is hardly found unless initial conditions where all particularly small (Fig. 3 top left). Because of this large basin stability and the fact that such a large perturbation is unlikely (see section I.E), we rarely see transitions towards extinction and hence few overlap values appear (Fig. 20 center and right panels).

At the other extreme of strong competition, mutual exclusion states are characterized by having one surviving species $D = 1$ as well as the extinct state, here more probable. The overlap between such states is either one or zero, meaning transitions between two states involve a total change of species because one competitor will disappear and another (or none) will be present (Fig. 20 blue dots).

At this point it was already obvious that these two categories, mutual exclusion and global bistability, harbored only very specific multistability flavors. The question is whether the two more vague counterparts,

[1] The relation between $D$ and $O$ could be also introduced by a metric for the relative shift $(D - O)/D$ or the Jaccard distance (see I.D.2 and (Liautaud *et al.*, 2019)). The relative shift value tells us if the number of species that has changed $(D - O)$ is large or small in proportion of the original number of species present $(D)$. This value is 1 for a total shift (no overlap, whatever the number of species) and smaller as the shift between a state and its perturbed counterpart share a larger fraction of species (not shown).

local multistability and cliques, expand all rest of possible $D$ and $O$ configurations.

Local multistability, by definition and as clarified by figure 2A in the Main Text, can take all possible diversity values. Higher cooperation will push towards seeing only the most populated configurations and so on (Fig. 20 yellow). As in the view of Gleason's loose collections of species (Gleason, 1926; Liautaud *et al.*, 2019), here states are separated by single-species shifts, so that the amount of species overlap are correlated with the diversity of states. Those states that have more species will have higher overlap, and shifts between them will be, in a way, less abrupt.

The final point that we find most interesting to clarify is whether cliques can take many diversity values. As discussed in the Main Text, we understand cliques as many possible community states but all with similar number of species. In figure 20 (green) we can see that cliques can take many diversity values. However, this depends on the specific system parameters. For a given system (imagine we explore a bacterial community in the laboratory with a constrained mean interaction strength), the diversity that cliques can take (the vertical width of the green distribution at a given value) is small. Moreover, for some of the models, we see that the diversity of cliques is in fact similar throughout interaction strengths, which is something that is still more shocking and remains a question for future work (Fig. 20 A, top left panel).

Another key aspect of cliques is that, as for the rest of state types, the species overlap between states is in general almost as large as their diversity. In general, transitions between large cliques are often relatively small: most species are shared between the original and the perturbed state. It is only for smaller cliques under competition that transitions can be more relevant, as all species can be shifting between two small cliques (Fig. 20 right panels, $D \leq 10$).

The take-home message is the following: a given community (under an interaction scheme $(A, B, \sigma)$) can only allow for multiple stable states of similar diversity: except for the large and rare shift of global bistability, the rest of multistability categories harbor states that are conceivably similar and close to one another. Interestingly, this further indicates a missing ingredient for catastrophic shifts. Generally speaking, our understanding of catastrophic shifts comes as a shift from a species-rich state ($D >> 1$) towards a state where many species are no longer present ($O << D$). This space (the $*$ area in the right panels of figure 20) is not populated by any state type. As discussed in the Main Text, we believe that there is something missing in the structure of the interaction networks for MSS to escape the Diversity-Overlap correlation.

##### 3. Number of states and their resilience to perturbations

As stated in the Main Text, one of the main outcomes of perturbing stable states is that we can analyze their fragility in a simple test that could be replicated in the laboratory. In particular, we define and apply a pulse perturbation in species abundance as $x_i^* + \Delta x_i \ \forall i$. Here $\Delta x_i$ follow a normal distribution with zero mean and varying (but not infinitesimal, hence the capital $\Delta$) standard deviation. For the sake of generality, we set perturbation width random with $\sigma$ following a uniform distribution within $(0.001, 3.0)$, which can of course be modulated.

What we ask ourselves is how often a state relaxes to its original composition after the perturbation. We define $R$ as the fraction of states that recover their original abundances after $s$ simulations, so that $R = s_R/s \in [0, 1]$. Resilience of course involves a much more complex analysis (Dakos and Kéfi, 2022), and basin stability (Menck *et al.*, 2013) should be computed by accurately analyzing the effect of different perturbation widths on a per-state basis. However, in this already simple setting we are able to recover an interesting result.

As shown in figure 4C in the Main Text, the fraction of simulations that relax to the same state correlates negatively with the number of possible states in place. The first notion is easy to understand: the more stable states present in an $N$-dimensional landscape, the smaller their basins of attraction should be. Interestingly, our species are quasi-neutral in the sense that their intra-species parameters are sorted at random with small standard deviation (see section I.C.2). Because of this, there is no reason to believe there are strong asymmetries between the stability basins of states.

In a perfectly symmetrical scenario with $\Omega$ stable states, and considering we explore a finite landscape with $x_i(0) \in [0, x_M] \ \forall i$ that already contains access to the basins of all states, the $N$-dimensional volume would be approximately divided in $\Omega$ basins. Each state would then hold a basin volume $BV$ of

$$BV \approx \frac{x_M^N}{\Omega} \tag{41}$$

This does not consider that the $\boldsymbol{x} = \boldsymbol{0}$ state is likely to hold a different $BV$ from all others. Still, because the likelihood of a state to recover should correlate with its $BV$, the question is therefore to test the $R \sim BV$ correlation (Fig. 4C in the Main Text). Interestingly, the trend is indeed followed and $R \sim \Omega^{-1}$ seems to posit a lower bound. In fact, most states are slightly more resilient than that. This is likely because of the variation of perturbation width: some perturbations are possibly much smaller than the scale of $BV$ and do not reach the borders of the state basin. The scope of the present work is not to delve deeper into high-dimensional resilience. We expect, however, that $R \sim \Omega^{-1}$ becomes the rule for uniformly large perturbations. Moreover, we believe that testing this theoretical result in species-rich communities in the laboratory could provide further understanding of the symmetry between stable states in high-dimensional systems.

#### III. OTHER MODELS OF COMPLEX SYSTEM MULTISTABILITY

Following the complex systems approach of the present work, we here study a set of minimal models that characterize dynamical processes in complex systems from different research fields. Aiming at a certain degree of generality, here we study multistability categories in ecology (the classical generalized Lotka-Volterra (GLV) model), transcription processes in gene regulatory networks (GRNs), the cancer-immune interplay at the cellular level (Garay-Lefever (G-L) model, here expanded for the first time to a multiple-antigen scenario) and the dynamics of neural clusters in the brain.

All models can be written in terms of systems of ordinary differential equations. Furthermore, we have chosen these models as they represent classical complex systems, and also allow for interesting comparisons, each harboring a characteristic property not found in others. Briefly, the model of the Main Text has two interaction layers (matrices $A$ and $B$). The GRN model describes a similar dynamical scheme of bistable gene activation and inhibition, but encapsulated in a single interaction layer $A$. The GLV model is built by monostable units, and hence will not show all the four multistability categories. In comparison, the G-L cancer-immune model follows very similar dynamics with linear interactions $A_{ij}x_j$, but non-linear immune-mediated death makes cancer populations bistable if independent. The neural model adds an interesting aspect, as activity stable states are either positive or negative, and the lack of a stable extinction state $x = 0$ precludes the emergence of mutual exclusion states.

Each of the models could by itself demand a whole research project to understand the complexity of the multistability landscapes involved. The GLV is of course a good example of that (see section II.F and table 1). However, our primary objective is to highlight the idea that, despite there are variations specific to each model regarding the dynamics of individual units and their interactions, the various flavors of multistability observed in all models fall within the same classes described in the Main Text. To do so, we replicate two of the main results of the article: (1) the phase space for how $\Omega$ depends on mean interaction strength and standard deviation and (2) the $(U, \Omega, D)$ space of empirical fingerprints.

Additionally, we are currently expanding our analysis to other complex systems beyond strictly biological systems, with a special interest on complex models of bistable units and multiple interaction types that can harbor 4 (instead of 2) of the studied MSS categories. Potential candidates could be the Preisach model of high-dimensional hysteresis (Mayergoyz and Friedman, 1988), models of bistable microelectrode arrays (Salman *et al.*, 2021) or language models involving competition and conversion (Zanette, 2008).

##### A. Generalized Lotka-Volterra interactions

The generalized Lotka-Volterra (GLV) model with random interactions of mixed sign is a benchmark model of systems with many linearly-interacting units, not only ecological communities (Bunin, 2017) but also other systems such as economical stock markets (May *et al.*, 2008; Solomon *et al.*, 2000). One particular benefit of the random approach is that, to model a system of very large size, it approximates unknown interaction coefficients from randomized distributions, an approach that has proved useful in uncovering key results in complex systems (May, 1972b) (see (Meena *et al.*, 2023) for a recent analysis on the validity of randomly selecting coefficients as a prior to evaluate stability). In one of the possible ways of writing the GLV model, the abundance of species $x_i$ follows

$$\frac{dx_i}{dt} = x_i \left( r_i - d_i x_i + \sum_{j \neq i}^{N} A_{ij} x_j \right) \tag{42}$$

where $r_i$ is the species replication rate, $d_i$ its self-regulation or logistic growth term via a carrying capacity $d_i = K_i^{-1}$ , and $A_{ij}$ stands for the interaction strength of species $j$ on $i$, that can take either positive and negative values, hence encompassing a large set of ecological interaction scenarios.

As compared to the rest of studied models, the GLV is particular to our work because single-species dynamics follow the well-known logistic growth model

$$\frac{dx}{dt} = x\,(r - dx) \tag{43}$$

that does not harbor any species-level bistability: species abundance will grow to $x* = r/d$ (or $x^* = K$, the species' carrying capacity) if $r > 0$, or become extinct for negative growth rates, but cannot be at either one of the two stable states under a given $r$ (Fig. 21A).

By studying a system with monostable species, we can now compare it with other models that do have single-species bistabilities, and uncover how multistability categories emerge from low-dimensional bistability or from the interactions imposed by $A$.

Because we do not need to study a specific parameter region where single-species bistability is in place, our qualitative results are independent from both $r_i$ and $d_i$, and the key elements of the system are the mean and variance of the sorted interaction strengths $A_{ij}$. Here we study a generic parameter region of the model by setting $\langle r_i \rangle = 1$ and $\langle d_i \rangle = 0.25$, and the connectivity of the Erdős-Rényi graph equal to one. This approach provides enough qualitative understanding of multistability and will also allow comparison with the bistable models of section III.C. In this scenario, the 1-d system stabilizes at $x* = r/d = 4.0$.

As expected from previous research thoroughly described in section II.F and Table 1, the GLV model does not show all of the 4 multistability categories described in our work, but only those 2 that can arise in the absence of bistable dynamics. Because GLV species are monostable when alone, competition lower than self-regulation does not allow for different present-absent competing compositions (local multistability), but a single state of diminishing diversity (coherent with phases I and II in (Kessler and Shnerb, 2015)). Another main difference with competition between bistable species is the absence of global extinction at high competition. Because species are monostable and simply grow to $x* = r/d$ when alone, the last survivor will always remain in place. As seen below, this also precludes the possibility of an all-or-nothing globally bistable regime, the *nothing* scenario being unstable in GLV under species remigration (Fig. 22A, bottom right).

Briefly put, our formulation of the GLV model displays a cliques and a mutual exclusion phase (Fig. 22A). As done in section II.F, it is interesting to discuss these results in comparison with the large body of work displayed in table 1.

- **Cliques:** As expected, we find that clique states emerge in the regime of high heterogeneity, weakly competitive interactions. We hypothesize this relates to the presence of heterogeneous interactions fostering an intransitive competition scheme (Soliveres and Allan, 2018). The clique phase of figure 22A is best understood in comparison to Phase III in (Kessler and Shnerb, 2015), also discussed in the Supplementary Material of (Liautaud *et al.*, 2019), and is consistent with the very recent results presented in (Mallmin *et al.*, 2023) (see page 7, Figure 5). As discussed in section II.F.2, however, it is interesting to find that cliques in our setting can harbor both chaotic regimes (a pinball walk through metastable states, see II.F.2 and figure 15) as well as stable configurations, something only recently discussed in (Mallmin *et al.*, 2023). Different model settings ( e.g. admitting an infinite species approximation with symmetric (Biroli *et al.*, 2018) or antisymmetric (Ros *et al.*, 2022) interactions, finite species models with stochasticity (Kessler and Shnerb, 2015), ..., see Table 1) find that all such cliques should be unstable. However, stochastic models (Kessler and Shnerb, 2015), few-species simulations (Bunin, 2017) or those under binomial (yes-no) competitive interactions (Fried *et al.*, 2016, 2017) do find that, for higher competition, cliques can become stable (This happens possibly when inside the mutual exclusion phase of figure 22A, but provided particular interaction or noise mechanisms are in place, see below). The analytical tools needed to explain these partially stable scenario are far from trivial and beyond the scope of the present work. We cannot provide an analytical proof able to accurately discern how the specific flavor of the cliques phase (partial stability) is related to that of other models, and refer the reader to the very recent publication (Mallmin *et al.*, 2023) where qualitatively equivalent results to those of figure 22A are achieved. Because of this, the take-home message here is that, for a variety of interaction scenarios linked to heterogeneous competition, a set of clique states can emerge that are stable and hence classify as multiple stable states.

- **Mutual exclusion states:** For very low heterogeneity, a mutual exclusion phase emerges under $\langle A_{ij} \rangle > \rangle d_i \langle$ (Fig. 22A), a well-known result (Armstrong and McGehee, 1980). One very relevant analysis here is that of (Kessler and Shnerb, 2015): the critical point where the three phases converge at $\sigma = 0$ is linked to Hubbel's neutral theory of biodiversity (Hubbell, 2011). In this context, the mutual exclusion phase is linked to Phase IV in (Kessler and Shnerb, 2015), similarly discussed in (Lischke and Löffler, 2017), and highlighted in (Mallmin *et al.*, 2023). Once again, this phase admits subtle variations depending on the specific characteristics of the model, although it has always been related to the emergence of stable states instead of chaotic fluctuations (Bunin, 2021; Fried *et al.*, 2016, 2017; Kessler and Shnerb, 2015). In other models, more than one species can coexist in such states under the presence of migration stochasticity (Kessler and Shnerb, 2015) or considering binomial competition models where, of course, pairs of species can be found that do not mutually exclude each other (an $A_{ij} = A_{ji} = 0$ interaction pair) (Fried *et al.*, 2017). Interestingly, as heterogeneity increases, the mutual exclusion phase needs higher competition to emerge. This is consistent with Figure 1 in (Liautaud *et al.*, 2019), where a mutual exclusion pattern is seen for low $\sigma$ (community pattern iv) that becomes a clique pattern for higher $\sigma$ but the same mean competitive strength (community pattern iii). We hypothesize that higher heterogeneity, by dismantling the competitive hierarchy that determines who excludes who, allows for moderate species coexistence a little beyond $\langle A_{ij} \rangle > \rangle d_i \langle$.

We also generate many GLV systems and plot their observables in the $(U, \Omega, D)$ space proposed along the Main Text. By keeping the same axes limits of the main cube ($\Omega_{max} = s = 200$ simulations, $D_{max} = N = 50$

species) we are able to see the observable trait signatures that GLV multistability would leave as compared to other complex systems models. It is also relevant to remember that we are plotting the $\Omega$ axis starting at $\Omega = 2$: there are many scenarios with one single state and variable diversity, as predicted by (Bunin, 2017; Kessler and Shnerb, 2015), that are omitted as they do not show bistability. States in this phase are not accompanied by the global extinction state $x^* = 0$.

Interestingly, we observe the same clique and mutual exclusion signatures of other models (Figure 4A and 4D in the Main Text): the mutual exclusion domain, happening for strong competition and low heterogeneity, has about $N$ states with one surviving species each. The clique domain, on the other hand, has a slightly higher amount of states and surviving species, together with a highly variable $U = 1 - S$ value: clique states can often be accompanied by few or a lot of chaotic or cyclic oscillations (Fig. 4D in the Main Text, top left, and fig. 15). The absence of the 2 other MSS types is consistent with our generic taxonomy: because species are not locally bistable, the system will not show the signatures typical of global bistability (the whole system replicating 1D bistability) nor local multistability (the system falling at one of many possible present-absent species states).

We have presented in the Main Text discussion a possible explanation of how these two MSS phases might relate to empirical evidence of multistability in ecology. We refer the reader to additional species-rich modeling frameworks that have provided interesting discussion on the link between different flavors of cliques and empirical ecological scenarios (Amor *et al.*, 2020; Bunin, 2021; Case, 1990; Fisher and Mehta, 2014; Hu *et al.*, 2022; Kessler and Shnerb, 2015; Liautaud *et al.*, 2019; Roy *et al.*, 2020).

##### B. Cancer-Immune interactions

In the following section we will show how T cell recognition of cancer neoantigens, a key feature of cancer-immune interactions, might replicate GLV dynamics under certain assumptions. Interestingly, research indicates that this could be so with an added non-linear death term that can imply local bistability. In particular, we will extend a classical cancer-immune model (Garay and Lefever, 1978) to account for present-day knowledge of cancer heterogeneity at the neoantigen level (McGranahan *et al.*, 2016).

Cancer is a special disease, where the host's own healthy cells undergo a process driven by ecological, evolutionary and developmental alterations to become a rogue tumor (Aguadé-Gorgorió *et al.*, 2022). To overcome multiple homeostatic barriers, this cellular bulk becomes devoid of multicellularity traits and able to colonize the tissue and progress until the host's death. Within this context, one of the key barriers that tumor cells need to overcome is that of immune suppression. The immune system itself a complex system composed by many layers of cellular interactions (see e.g. (Perelson and Weisbuch, 1997) for a description). Among all this complexity, cytotoxic T lymphocytes (CTLs) belonging to the adaptive immune system are nowadays known to be able to recognize mutated surface proteins in cancer cells (known as cancer *neoantigens* (Schumacher and Schreiber, 2015)). With this, immune cells activate a non-self recognition-and-attack process that is able, under specific conditions, to eradicate neoplasias. This utterly complex process, however, was already thought possible more than a century ago (Ribatti, 2017) although its mechanisms where far from understood.

Recent decades have seen an immense improvement in how cancer-immune interactions are understood and, more importantly, harnessed towards the development of immunotherapy (Esfahani *et al.*, 2020). However, minimal mathematical models, despite lacking precise description of some specific molecular processes in place, where already able to provide insight into the mechanisms of immunosurveillance. One powerful example of this was provided by the Garay-Lefever model of cancer-immune system interactions (Garay and Lefever, 1978).

This cell-level model coarse-grains, as done across cancer-progression dynamical models, tumor populations to a single compartment $C$. This population can in turn be attacked by a T-cell population $T$. The main dynamics are cancer replication at rate $r$, and the formation of a cancer-immune cell-cell complex $O$ at rate $k_1$ and subsequent cancer death at rate $k_2$ (Solé, 2011):

$$C \xrightarrow{r} 2C \tag{44}$$

$$T + C \xrightarrow{k_1} O \xrightarrow{k_2} T + \varphi \tag{45}$$

By admitting linearity except for the commonly accepted cancer logistic growth, the system writes

$$\frac{dC}{dt} = rC\left(1 - \beta C\right) - k_1 CT \tag{46}$$

$$\frac{dT}{dt} = k_2 O - k_1 CT \tag{47}$$

The two-dimensional system can be simplified by assuming that the $O$ complex is short-lived and the dynamics of $T$ can be assumed to be at steady state $dT/dt \approx 0$ (Garay and Lefever, 1978; Solé, 2011). Defining a total constant population of effector cells as $E = T + O$ and admitting the steady state approximation, we obtain

$$T(C) = \frac{E}{1 + \frac{k_1}{k_2}C} \tag{48}$$

that allows us to write the dynamical expression for tumor growth

$$\frac{dC}{dt} = rC(1 - \beta C) - \frac{k_1 CE}{1 + \frac{k_1}{k_2}C} \tag{49}$$

often expressed in rescaled quantities

$$\frac{dc}{dt} = c(1 - \theta c) - \frac{\mu c}{g + c} \tag{50}$$

where the new term $g$ incorporates a proxy for how good the tumor bulk precludes immune penetration: higher $g$ indicates immune kill increases lower with tumor size $c$.

What are the potential dynamics of this 1D system? The 1D system can have three attractor states, one at $c^* = 0$ and the other two at

$$c^*_{\pm} = \frac{1}{2\theta}\left(1 - \theta g \pm \sqrt{(1-\theta g)^2 - 4\theta(\mu - g)}\right) \tag{51}$$

By graphical analysis of the gradients of the growth and death curves at $c = 0$ (Fig. 21B), one can prove that the this pair of attractors (of which $c_+$ will be stable) happens if

$$\frac{d}{dc}\left(c(1-\theta c)\right) < \frac{d}{dc}\left(\frac{\mu c}{g + c}\right) \tag{52}$$

This will be fulfilled if $\mu > rg$: immune-mediated death has to be larger than cancer growth times cancer cooperation (via bulk protection). If not, the system behaves qualitatively as a 1-dimensional logistic growth in terms of only having a stable positive attractor, cancer outgrowth.

The possibility of intrinsic cancer-immune bistability provided the notion that, under certain conditions on immune recognition and killing, the tumor could exist in a landscape where both total immune surveillance ($c^* = 0$) and immune escape ($c^* > 0$) steady states exist, providing a first overview of the potential mechanisms explaining why some tumors escape the immune system while others become arrested early (Fig. 21B).

Understanding the cancer-immune interplay and the role played by neoantigens and subsequent immune recognition $\mu$ has prompted massive improvements in the success of immunotherapy (Schumacher and Schreiber, 2015). A major caveat here is that tumors do not harbor a single neoantigen, but many, each with a markedly different immunogeneicity (McGranahan *et al.*, 2016). Because T cells can only recognize a single neoantigen at a time, it becomes necessary to introduce the fact that the tumor population $c$ now includes a subset of cancer clones, each with dynamics that are marked by a different immune-mediated recognition of their most dominant antigen (see (Aguadé-Gorgorió and Solé, 2020) for a more detailed account).

$$\frac{dc_i}{dt} = c_i\left(1 + \sum_j A_{ij}c_j\right) - \frac{\mu_i c_i}{\sum_j (g + c_j)} \tag{53}$$

Two major considerations need to be discussed for our multispecies version of the Garay-Lefever model. First, we have included the otherwise acknowledged fact that cancer clones interact, mostly due to space and resource competition, with themselves and other clones (Gatenby, 1991). Together with these, cooperative strategies have also been recognized possible across tumor types (Archetti and Pienta, 2019). For simplicity, all this is encapsulated in the elements of the $A$ matrix following linear GLV dynamics.

Together with tumor-tumor interactions, we also need to take into account the meaning of the immune death term. Here, the strength of immune-mediated death $-\mu_i c_i$, where $\mu_i$ accounts for the immunogeneicity of the dominant antigen in that clone (Aguadé-Gorgorió and Solé, 2020), is divided by a non-trivial term. The expression $1/\sum_j (g + c_j)$ incorporates the spatial effects of the tumor bulk on immune attack. Penetration by lymphocytes is hampered by dense cellular masses, and this is independent of what neoantigens they harbor (Kuznetsov *et al.*, 1994). Hence, the death rate of a given cancer population is reduced by cooperation through bulk-level protection.

It is also easy to understand why cancer cooperation takes the form of $1/\sum_j (g+c_j)$ instead of $1/(g+\sum_j c_j)$. If we suppose a quasi-neutral scenario with all parameters equal along clones no inter-clone competition, we could envisage that all clones stabilize at a similar $\bar{c}$ state. Then, we would have that immune death of a single clone at stability $\bar{c}$ decays as more rogue populations preclude immune penetration:

$$-\mu \frac{\bar{c}}{\sum_j (g + c_j)} \approx -\mu \frac{\bar{c}}{Ng + N\bar{c}} = -\frac{\mu}{N}\frac{\bar{c}}{g + \bar{c}} \tag{54}$$

And so bistability happens if $\mu > Nrg$: now we see that more cancer clones $N$ can generate what has been previously proposed as a neoantigen diversity threshold (Aguadé-Gorgorió and Solé, 2020; Nowak *et al.*, 1991). What the dynamics tell us is here that there is a maximum amount of different neoantigenic clones $N$, beyond which the immune surveillance state $c^* = 0$ is no longer stable, consistent with empirical evidence on the role of heterogeneity (McGranahan *et al.*, 2016). If, instead, the dynamical shape took the form $1/(g + \sum_j c_j)$, the bistability threshold would still happen at $\mu > rg$ at least at the quasi-neutral case. This would be a possible case in the limit where immune cells and their TCR repertoires where infinite: more cancer clones would be responded with more immune clones and dynamics would not be affected by heterogeneity. This would mean that each cancer clone would increase immune recognition as well as decrease immune penetration equivalently, but the first does not happen due to the divide-and-win phenomena of neoantigen heterogeneity (Nowak *et al.*, 1991).

The possibility of cancer clone bistability implies that the above system is in fact a very interesting realization of the previous Generalized Lotka-Volterra in a cell-cell interaction environment dominated by chemical-recognition processes. As compared to the ecological scenario, here each single cancer cell compartment in the system is locally bistable through an indirect Holling type-1 death rate (Fig. 21).

When compared to the GLV model, what is the effect of this additive local bistability, combined with cooperation by cellular populations? What we find is that in fact $g$, a proxy for how hard it is for the immune system to penetrate the tumor, strongly dictates the flavor of the multistability landscape involved, as well as obviously $r$ and $\mu$. As so, the approximated quasi-neutral threshold $\mu^c = Nrg$ dictates two very different scenarios. Under high-enough immune recognition $\mu > Nrg$, the system is locally bistable and reproduces all four possible multistability categories (Fig. 22B).

There are however certain considerations to discuss here. As opposed to the rationale by which cooperating bistable species should upscale to produce global bistability (Gao *et al.*, 2016), here the global phase is particularly small and bounded between null and very weak cooperation. This is because, as in the GLV, cooperation happens through a linear term. This means that strong cancer clone cooperation does not drive the tumor to a stable positive attractor, but rather results in tumor exponential outgrowth. At the other extreme, strong linear competition, intertwined with non-negligible immune-mediated death (the tumor is no longer very big and immune cells can penetrate) drives the system rapidly to extinction.

In between these two, there is nevertheless a domain of local multistability not seen in the GLV model, where not all clones necessarily coexist but many possible intermediate states are in place (Fig. 22B, yellow region). These clones can sustain a small competitive strength, allowing for a mutual exclusion phase that is rapidly erased by extinction. At increasing interaction heterogeneity, clique states emerge and can endure a larger region of the phase space due to the interplay between weak and strong competitors. All in all, this is an interesting take on how, even if the phase diagram of a system can take many different shapes, the qualitative nature of each MSS phase remains similar, as shown for the fingerprints below.

On the other hand, if either $\mu$ is too low or cancer bulk-level cooperation $g$ too high, the system abruptly changes to single-species monostability at $c^* > 0$ and recovers the GLV phase diagram (not shown). As expected from our taxonomy, this also means that global bistability and local multistability states disappear, and only competitive domains remain in place.

As done for the rest of the models, we also plot the potential signatures of each MSS family (Figure 4D in the main text). We find once again the same qualitative fingerprints emerge. There is a very narrow scenario where all cancer clones cooperate and coexist together or else go extinct (leaving the global bistability signature), that might easily fall towards cancer outgrowth under slightly increased cooperation. We now know that localized tumors tend to stabilize their growth dynamics (Norton, 1988), and that it is not exponential growth but metastasis, and its potential emergence under tumor clone heterogeneity, what makes cancer particularly lethal (Fidler, 1978). Heterogeneity here can be understood in the lens of local multistability or, because interactions between cancer clones that hold phenotypic differences might be heterogeneous (van Neerven and Vermeulen, 2023), a novel concept of cancer cliques. We hypothesize that further developing the meaning of the cliques phase in the context of tumor progression could explain how many cancer genoptypes and phenotypes can coexist in the tumor. The fact that cliques are separated by abrupt shifts that do not change the diversity of the tumor but do imply a change in phenotypes could also shed light in our understanding of treatment failure through tumor adaptation resulting from community-level plasticity (Fisher *et al.*, 2013).

Furthermore, it is interesting to discuss the fact that increased cooperation through bulk protection (increasing $g$ (Chen and Mellman, 2017)) or reduction of immune find-and-kill capacity (reducing $\mu$ (Kim, 2007) or increasing $N$ (Aguadé-Gorgorió and Solé, 2020; McGranahan *et al.*, 2016)) drives the system to a completely different multistability scenario that is qualitatively equivalent to the GLV model (Fig. 22A). Besides the MSS phases discussed above, the total extinction phase disappears in this new scheme, explaining the multiple ways in which cancers are able to escape the selective barrier of immune attack (Sharma *et al.*, 2017).

##### C. Gene regulatory networks

Genes and the products they encode for are crucial for life. Interestingly, genes and their overall activity are not independent from one another. In fact, the expression level of a gene affects many others and can be in turn affected by them. The state of this complex network made up of thousands of genes and their interactions then controls a myriad of processess such as cellular function or morphological development (**?**). It appears natural that such networks do not exist in a single state, but rather can switch between many configurations involving different gene expression levels.

Early models of gene regulatory networks (GRN) proposed a Boolean perspective on gene activity, meaning that genes behave as on or off switches (Kauffman *et al.*, 1993). In Kauffman's original $NK$ model, $N$ genetic switches interact with $K$ others via *epistasis*. When $K = 0$, each gene simply participates additively on genetic expression or *fitness*, hence a single optimal on-off combination exists that maximizes this function. However, as $K$ increases and the GRN becomes interwoven, interactions start frustrating the on and off switches in a similar fashion as in the Ising model (Cipra, 1987) or spin glasses (Toulouse *et al.*, 1987). Already in this simple model there is evidence that many possible (in fact, exponential) gene expression combinations can arise from a single GRN architecture (Kauffman and Weinberger, 1989; Weinberger, 1991). Because there might be about $N \lesssim 2 \cdot 10^4$ interacting genes, this result shed light in the underlying complexity of epistasis.

More detailed models have described gene activity as a continuous process (Bornholdt, 2008). In the context where a protein is assumed to form a dimer to perform its regulatory function, gene products can then activate the genes via positive feedback loops or else inhibit their expression (Isaacs *et al.*, 2003). In this context, a model for the concentration of a genetic product $p_i$ can be described to follow (Solé, 2011)

$$\frac{dp_i}{dt} = -\delta_i p_i + \alpha_i \frac{p_i^2}{1+p_i^2} \tag{55}$$

Interestingly, this indicates that the expression of each gene is in itself a switch: besides the trivial fixed point $p_i^* = 0$, there exist a couple of unstable $p_{i-}^{*,U}$ and stable $p_{i+}^{*,S}$ fixed points with positive genetic activity

$$p_{i\pm}^* = \frac{1}{2}\left(\frac{\alpha}{\delta} \pm \sqrt{\frac{\alpha^2}{\delta^2} - 4}\right) \tag{56}$$

This is easy to visualize by plotting growth and death curves and their intersections (Fig 21C).

Knowing that gene products alter the transcription of others via positive and negative effects, what are the possible dynamical outcomes of the networked system? How many possible states can a single model endorse? On the one hand, dimensionality reduction techniques propose that the whole network will behave as a 1-dimensional bistable switch (Gao *et al.*, 2016), while the well-established NK-model of Boolean networks first proposed by Stuart Kauffman already envisioned that many possible configurations could exist, endorsing a so-called *rugged* fitness landscape (Kauffman *et al.*, 1993) as a key first insight on the potential complexity of genetic epistasis (Lagator *et al.*, 2017).

Our point here is to study the emergence of multiple stable states of the following complex dynamical system

$$\frac{dp_i}{dt} = -\delta_i p_i + \sum_j A_{ij} \frac{p_i^2}{1+p_i^2} \tag{57}$$

This system only contains one layer of either positive or negative interactions: genetic products, as opposed to species, are assumed here to interact via a single process only (here encoded in $A$) and cannot undergo both positive and negative interactions at the same time. With this in mind we can directly plot in a 2-dimensional map the effects of modulating the mean genetic interaction strength $\langle A \rangle$ and its standard deviation $\sigma_A$ (Fig. 22C).

We can see that, depending on interaction types, the four MSS categories found for the multilayer ecological model of the Main Text also emerge in the context of bistable gene regulation (Fig. 22C). If the interaction of genes with genetic products was strongly governed by activation, as proposed in (Gao *et al.*, 2016), mutual cooperation makes the dynamics of each node synchronous, and the system replicates low-dimensional bistability (Fig. 22C, purple). At the other extreme, if interactions were mostly negative, one would see that only one gene product can remain in place, and the system would switch between who wins, depending on initial conditions (Fig. 22C, blue). At the extreme of very strong inhibition not even a single gene could survive, due to its activity being rapidly hampered by a small fraction of inhibiting proteins. These inhibitory population would likely be maintained not by migration, as in the ecological case, but by noise in the genetic expression of products even if previously inhibited by the non-stochastic dynamics (Raser and O'shea, 2005).

However, the interactions encoded in real GRN's probably seat between these two extremes (?). Assuming interactions are small and of both signs ($\langle B \rangle \approx 0$), two more MSS categories are in place. If interactions are weak and homogeneous, each genetic switch functions independently of others and up to $2^N$ states could be observed (Fig. 22C, yellow), with selection for function easily choosing the best combination (Kauffman *et al.*, 1993). In the likely case that some genetic effects on expression are stronger or weaker than others, the fourth category of MSS driven by heterogeneity would then emerge (Fig. 22C, green), consistent with early models predicting this rugged and highly multistable scenario (Kauffman *et al.*, 1993). Here cliques would of course not be groups of coexisting species, but rather those genes that can be expressed together while repressing others, often in cyclic or even chaotic behavior.

Interestingly, the signatures of these multistability categories are qualitatively equivalent to observed in the ecological model of the Main Text (Figure 4A and 4D in the Main Text). Positive or weak interactions between bistable genes can prompt an all-or-none switch (global bistability) as well as the extreme multistability of independent genes that would characterize a genotype-to-phenotype map with no genetic epistasis. On the other hand, when gene product inhibition is also taken into account, the system integrates the possibility of "genetic cliques", groups of genes that can be active together, as well as the one-wins scenario, an unlikely "mutual genetic exclusion" domain. Interestingly, one finds that in GRN models, the diversity of cliques can be much larger than for the other models in place. This prompts the analysis of how highly cooperative interactions, even under heterogeneity, can hold large cliques of mutually activating genes. This signature is not seen in GLV because positive heterogeneous interactions leads to species outgrowth (instead of large cliques or global coexistence). In the Allee Effect model of the Main Text, exploring cooperative $A_{ij}$ values much beyond $A_{ii}$ also accounts for cliques with higher species diversity, but also a higher fraction of fluctuating regimes (not shown).

Discussing the potential link of GNR multistability types and their signatures with empirical evidence is beyond the scope of the project and would also need us to take into account the relevant role of biological noise and stochastic dynamics. However, we hypothesize that a most likely scenario is that where gene inhibition and activation are approximately equipotent, and $\langle A_{ij} \rangle \approx 0$. Also, because GRN connectivity is much lower than one and node degree is heterogeneous, the idea of "genetic cliques" might be the one better describing the role of genetic epistasis in complex genoptype-to-phenotype mapping as in the picture proposed by (Kauffman *et al.*, 1993). We recommend the reader other references within the field that would allow for additional discussion (see e.g. (Bornholdt, 2008; Davidson and Levin, 2005; Isaacs *et al.*, 2003; Kauffman *et al.*, 1993; Mochizuki, 2005; Thattai and Van Oudenaarden, 2001; ?)).

##### D. Dynamics of neural networks

Large vertebrate brains are one of the most (if not the most) paradigmatic example of a complex system (Bassett and Gazzaniga, 2011; Sole and Montoya, 2001). The human brain in particular contains about 14 to 16 billion neurons, each connected by synapses to thousands of other neurons. How the well-understood functioning of neurons and their synapses upscales for consciousness and all its surrogates to emerge remains one of the central questions in science. Moreover, and because of this utterly high dimensionality, neural networks provide a benchmark example of how disordered systems approaches (see section I.C.3) have been useful to tackle complex systems: we can define general statistics of how neurons interact, but assigning specific identifiable roles is no longer useful (Barbier and Arnoldi, 2017; Mézard, 1989).

One interesting avenue of neuroscience and complex systems research has been the study of models of coupled oscillators (Zuo *et al.*, 2010). This particular field has been prominent in the search for dimensionality reduction techniques able to explain emerging, large-scale phenomena by simplified models (Bick *et al.*, 2020). Furthermore, the analysis of complex networks of oscillators is providing ample evidence that many non-trivial phenomena can emerge from the structure of the network itself (Arola-Fernández *et al.*, 2022), something that is in line with our concluding hypothesis of the Main Text.

On another level, it is also interesting to note that large-$N$ approximations have also been used in random neural networks to predict a stable-to-chaotic transition (Sompolinsky *et al.*, 1988) reminiscent of the one originally proposed in (May, 1972a) (and analyzed in depth in section II.F.1). All in all, multistability represents a fundamental problem in both brain dynamics (Golos *et al.*, 2015), where it might be linked to alternative states of consciousness (Kelso, 2012), as well as in neural networks, where each stable solution corresponds to different memory capacities of the network (Cheng *et al.*, 2006).

Tuning down the ambition of the task, here we study the possible multistability scenarios of a previously proposed firing-rate neural model. This family of models takes into account the fact that the synaptic input of a neuron can be separated into two clusters, a local and a distant one, therefore affecting the output dynamics (Litwin-Kumar and Doiron, 2012). This separation of spatial scales at the network level brought forward the possibility of defining the dynamics of not single neurons, but local clusters as the elementary units of the neural network (Wilson and Cowan, 1972).

One consequence of modeling network clusters that have an intrinsically strong self-coupling is the possibility that network units are bistable. This makes this specific frameworks interesting to our study, as we know that interactions between bistable units can provide a richer dynamical landscape. Following the original model by (Stern *et al.*, 2014), here the activation state of a cluster follows

$$\frac{dx_i}{dt} = -x_i + s\tanh(x_i) + g\sum_{j\neq i}^{N} J_{ij}\tanh(x_j) \tag{58}$$

where the self-coupling dynamics and long-range interactions are separated and modulated by coupling strengths $s$ and $g$ respectively. The original work considered an interaction matrix $J$ to follow a Gaussian distribution with mean 0 and variance $1/N$. Consistent with our analysis of other models, we here expand the model to incorporate scenarios modeling interacting clusters that are more predominantly excitatory ($\langle J_{ij}\rangle \gtrsim 0$) or inhibitory ($\langle J_{ij}\rangle \lesssim 0$).

At the unit level ($g = 0$), we realize that the attractors of the 1D system are inherently different from the rest of studied models. In particular, each cluster follows

$$\frac{dx_i}{dt} = -x_i + s\tanh(x_i) \tag{59}$$

which has an unstable solution at $x_i = 0$, and two stable attractors at $\pm x_i^*$. Hence, we are not dealing now with a present-vs-extinct or active-vs-inactive scenario, but rather with a model that resembles the original Ising model of up and down spins (Fig. 21D and (Cipra, 1987)). This is also relevant when building model simulations: because neural clusters do not go *extinct*, we will not add a migration term $+m_i$ in the system. As studied in depth in (Stern *et al.*, 2014), this will result in chaotic regimes being long-lived but finite, as pinball-like dynamics will gradually fade off into attractor states.

Again we plot the number of observed states $\Omega$, the fraction of initial conditions not stabilizing into a fixed point $U = 1 - S$ after a finite period of time, and the fraction $\phi$ of clusters that remain in the active ($x_i^* > 0$) domain against mean($J$) and $\sigma_J$ (Fig. 22D). For an averagely positive interaction matrix, indicative of mostly excitatory long-range interactions, the system realizes two possible stable states (Fig. 22D, purple). However, and because initial conditions are gaussian and centered at zero, global synchronization with all clusters at the positive or the negative state will depend on small differences on which "side" starts with advantage, generating a noisy domain (Fig. 22D right). At the other extreme, if $\langle J\rangle$ is negative, cohesion is broken and local bistabilities emerge, with many possible combinations of positive and negative activities in place, similar to the de-magnetized phase in the Ising model.

Interestingly, because local stable states are symmetric at $\pm x_i^*$, there is no well-defined meaning of *competition* here: a negative $J_{ij}$ term can both increase or decrease the activity state of $x_i$ depending on the sign of $x_j$. This in turn implies that there exist no mutual exclusion states: even under negative $\langle J \rangle$, a negative activity $x_i$ ensures that other clusters receive a positive impact from its $i \to j$ interactions.

As $J_{ij}$ becomes heterogeneous, triangular frustration appears in a way that resembles intransitive competition in ecology. Here not all states are possible and a small number of stable positive-negative configurations coexists with chaotic behaviors. As shown in (Stern *et al.*, 2014), these chaotic regimes will eventually stabilize to a fixed point, but their average transient live $\tau$ is exponential on the size of the network.

Again, we look at the signatures that this system leaves in terms of potentially observable traits: the fraction of stable neural regimes $S$, the number of different stable configurations observed $\Omega$ and the fraction of active clusters of each stable state $\phi = N_{active}/N$ (Figure 4D in the Main Text). Even if the model holds significant differences with the rest of studied systems, we find that again each multistability category leaves a similar signature. As there is no extinction but positive and negative neural activities, global bistability appears as an all-positive vs all-negative configuration, reminiscent of the magnetized phase in Ising's model (Cipra, 1987; Solé, 2011). In the more realistic scenario where interactions are not overly cooperative, as studied by (Stern *et al.*, 2014), the system shows too particularly different fingerprints: if interactions are weak, each cluster recovers it's independent positive-or-negative configuration and many local states appear, with the same amount of $+$ and $-$ units on average. On the contrary, when heterogeneity in the interactions between neural clusters kicks in, a smaller number of states emerges often precluded by abundant non-stabilizing states, consistent with the chaotic regimes studied in depth by (Stern *et al.*, 2014) (Figure 4D in the Main Text, bottom right).

It is beyond the scope of the present work to produce a discussion linking neural cluster multistability types and their signatures with empirical measurements in the brain or their relation with complex tasks. Nevertheless, one would expect that extreme scenarios of complete cluster synchrony (at $\langle J \rangle \gg 0$) and uniform multistability (at very low interaction heterogeneity or strong inhibition) are mathematical artifacts of the model. On the other hand, the existence of transitory fluctuations coexisting with stable multistability at $\langle J \rangle \approx 0$ is particularly relevant, as firing-rate fluctuations have been identified in cortical brain recordings (Churchland *et al.*, 2010; Korn and Faure, 2003) and could hold further information of potential synchronization and critical behavior in the brain (Cocchi *et al.*, 2017; Solé, 2011; Solé *et al.*, 2021).

#### SUPPLEMENTARY FIGURES

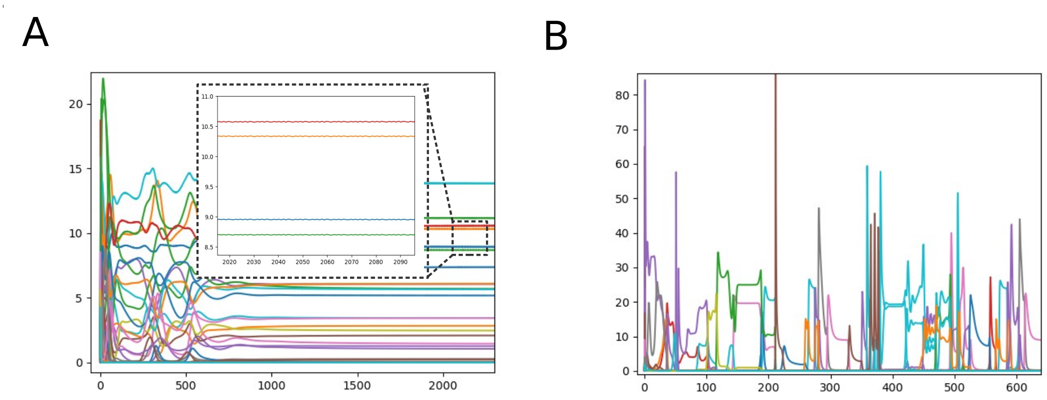


FIG. 1 **Fluctuations and numerical estimates of stability.** Numerical fluctuations in time can be either a result from small integration noise (A) or intrinsic fluctuations resulting from the dynamics (B). The dissimilarity index allows us to differentiate the first from the latter and therefore to admit as stable some states that are not exactly the same at floating-point precision.

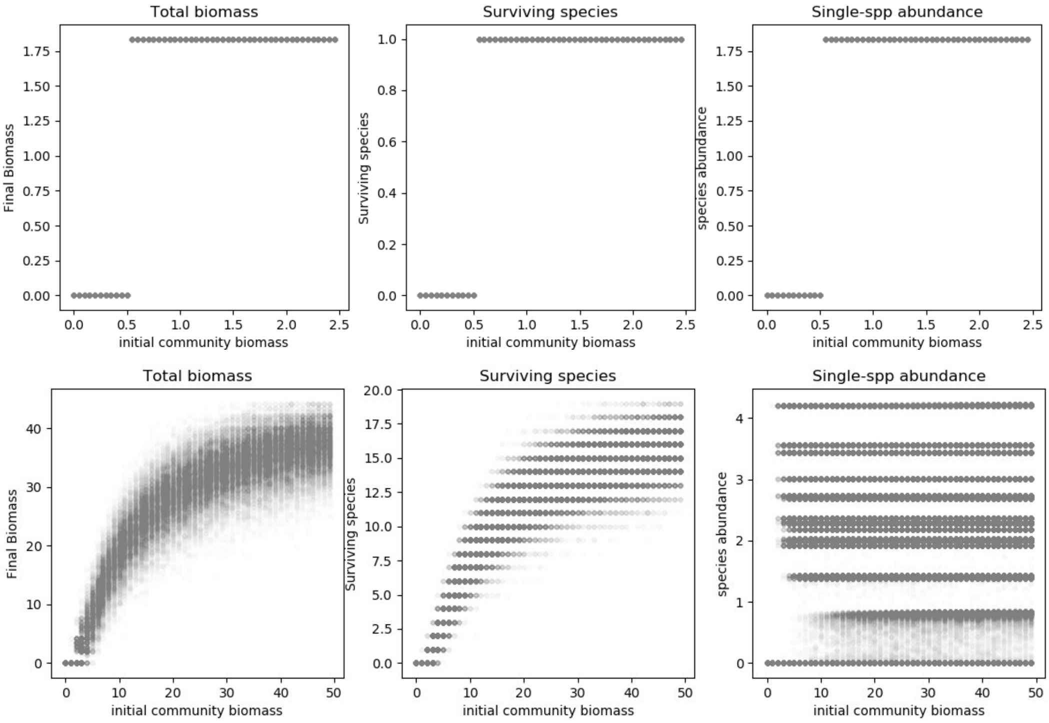


FIG. 2 **Multistability for one and many independent bistable species. Top:** One species undergoing bistable dynamics will fall in either one of two possible stable states depending on initial and threshold conditions ($x_i(t = 0) < x_i^{*,U}$). The presence of well-separated states indicates the possibility of a regime shift between species presence or extinction. **Bottom:** A community built on many of such non-interacting bistable species, however, will not be bistable, but harbor many states. Small changes in total initial conditions are only likely to move the system to a close-by state in terms of biomass or abundance of survivors.

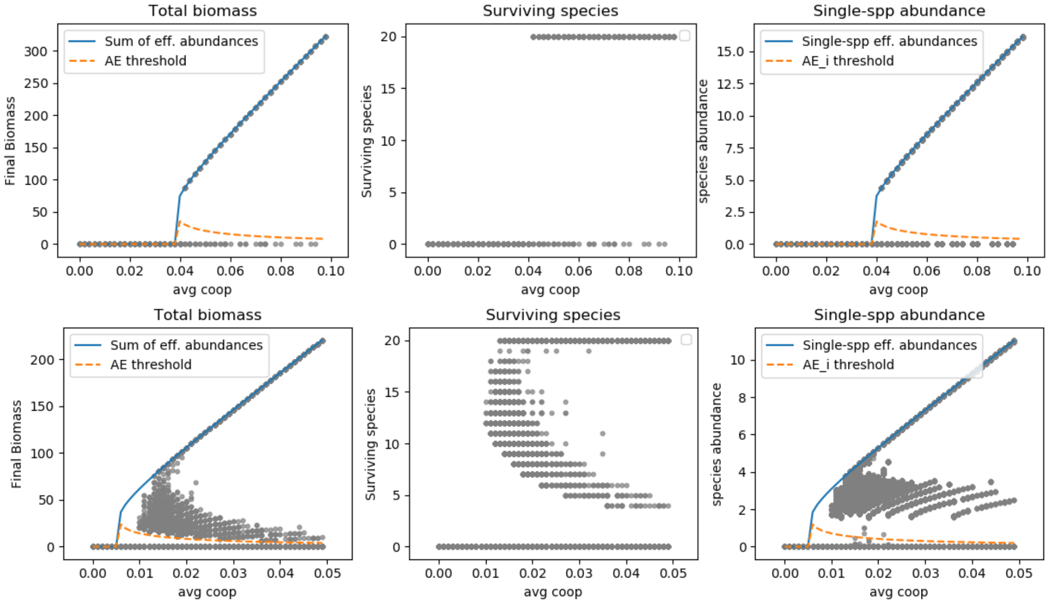


FIG. 3 **Multiple stable states in mutualistic communities: global *vs* local MSS.** Total biomass, total diversity and single-species biomass as we increase average mutualistic strength in Eq. (9). Blue and orange lines are predicted from the dimensionality reduction technique in section II.B.1: the blue line is the sum of $N$ species biomass, each with equivalent effective dynamics, while the orange line is the sum of $N$ species survival thresholds (each predicted to happen at the same value). Interestingly, the orange line also marks the basin stability of the extinct state, which we see is asymptotically small under increasing mutualism. Throughout the analysis, this will help us understand why $\boldsymbol{x} = 0$ is hard to observe when global bistability is in place. **Top:** A global bistability scenario can upscale from cooperation between locally bistable species, provided an homogeneous-enough community where species cannot survive alone. **Bottom:** If species can survive when alone, local present-absent compositions pervade the landscape of possible community states until cooperation strength governs the system. This is mostly relevant at the critical transition, where the system will be unlikely to undergo an all-or-none abrupt shift.

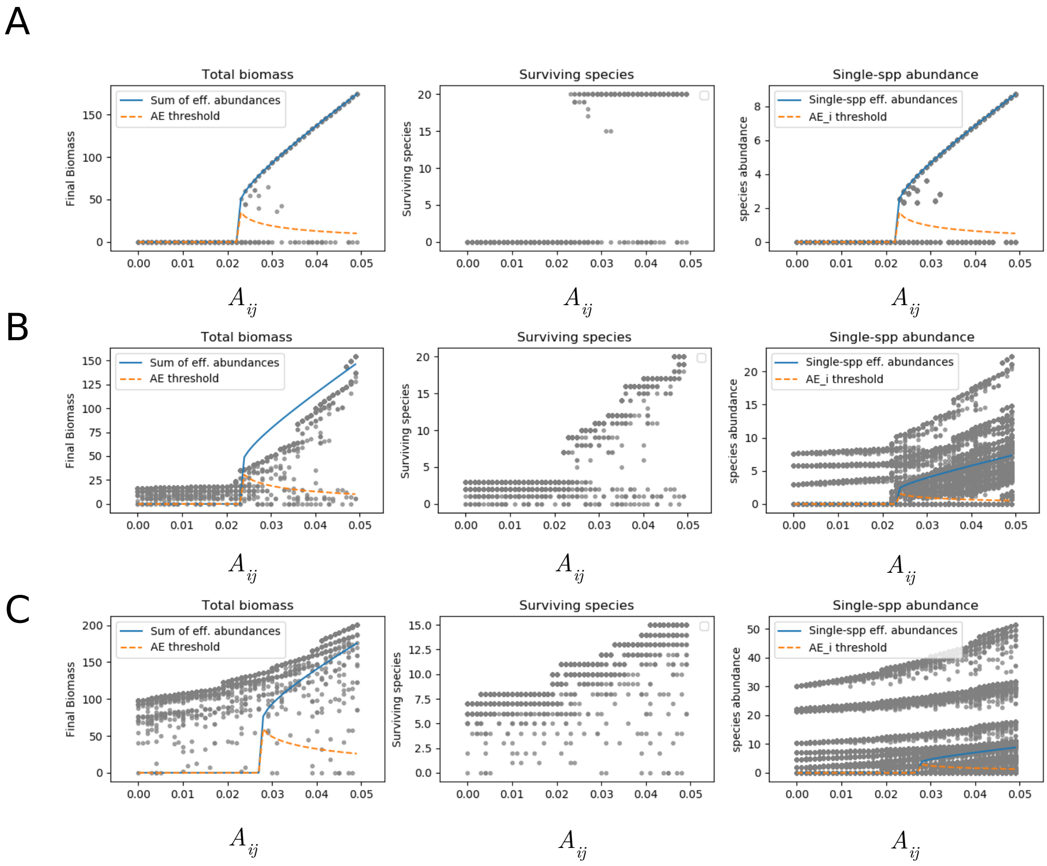


FIG. 4 **Global bistability is dismantled by heterogeneous species parameters.** Total biomass, total diversity and single-species biomass as we increase average mutualistic strength, for $\sigma_{ii} = 0$ (A), $\sigma_{ii} = 0.4$ (B) and $\sigma_{ii} = 1$ (C). Wide differences in single-species bistability thresholds result in the loss of species symmetry and the lack of a well-defined global bistability regime.

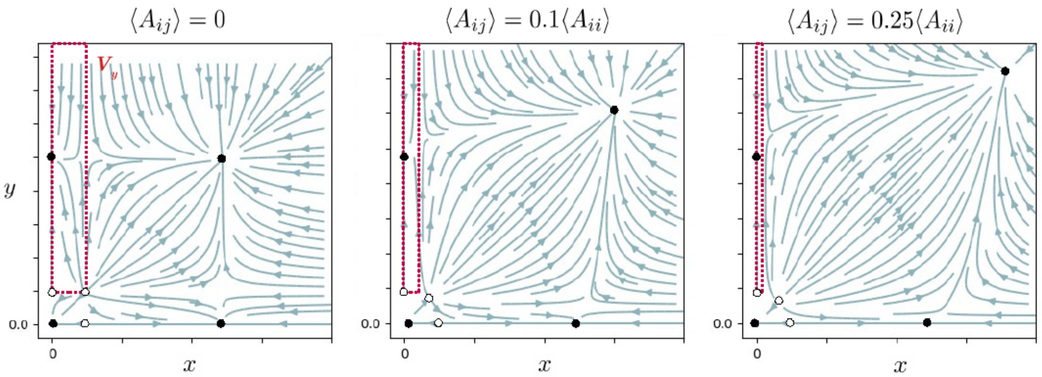


FIG. 5 **Global and local MSS in mutualistic 2D systems.** The red area corresponds to the volume of initial conditions $V_y$ where species $y$ would be present without species $x$, hereby dismantling the coexistence *global* state expected from dimensionality reduction techniques. These states might be of importance at low mutualistic strengths, meaning they can pervade a none-to-all transition (see Figures 2A and 2B in the Main Text). At high mutualism, all basins become negligible except that of the global survival state, explaining the observations of Fig. 4B-D of the main at $\sigma > 0.05$.

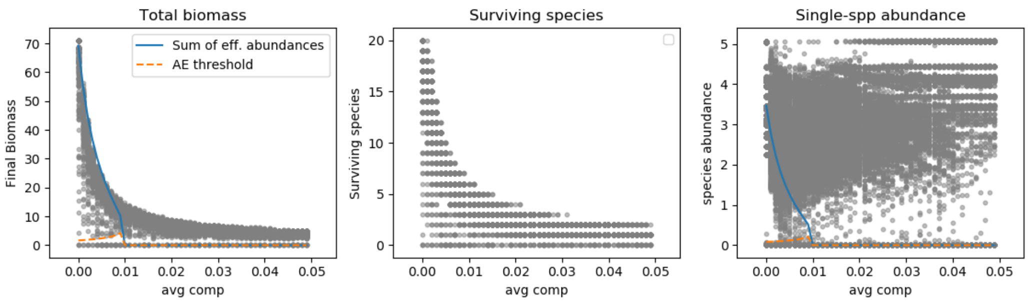


FIG. 6 **No global bistability in competitive communities of bistable species.** (Linked to figure 2B in the main text) Competition between bistable species with independent Allee Effects does not generate a globally bistable community. What we see is a combination of local present-absent species combinations, bounded by extinctions due to increasing competitive exclusion effects. The system only becomes bistable in the trivial case of only one species remaining. The dimensionality reduction procedure (Gao *et al.*, 2016; Laurence *et al.*, 2019) captures the decay of global biomass (left), but by predicting that all species feel the same increasing self-regulation and not go extinct. Symmetry is in fact broken as more species become extinct (center) and their individual abundances are non-uniformly altered by competition (right)

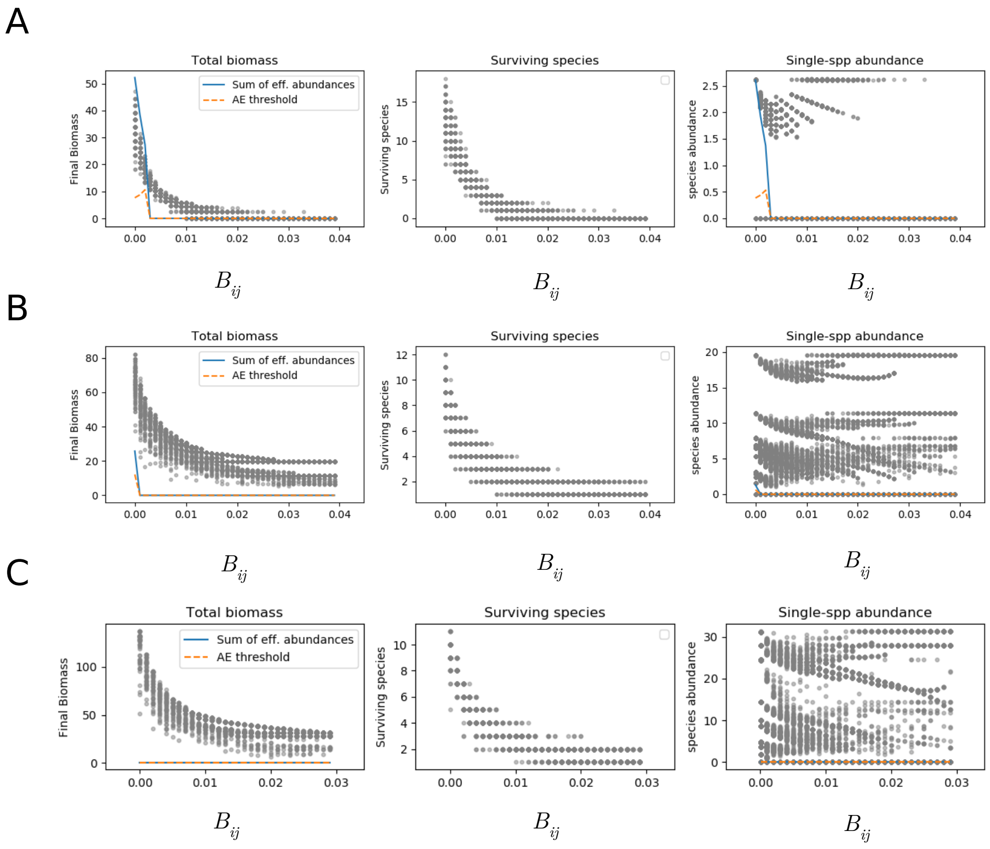


FIG. 7 **Local exclusion states under heterogeneous species parameters.** Total biomass, total diversity and single-species biomass as we increase average competition strength, for $\sigma_{ii} = 0$ (A), $\sigma_{ii} = 0.4$ (B) and $\sigma_{ii} = 1$ (C). Wide differences in single-species bistability thresholds maintain a transitive competition scheme in place, further reinforcing the effects of mutual exclusion.

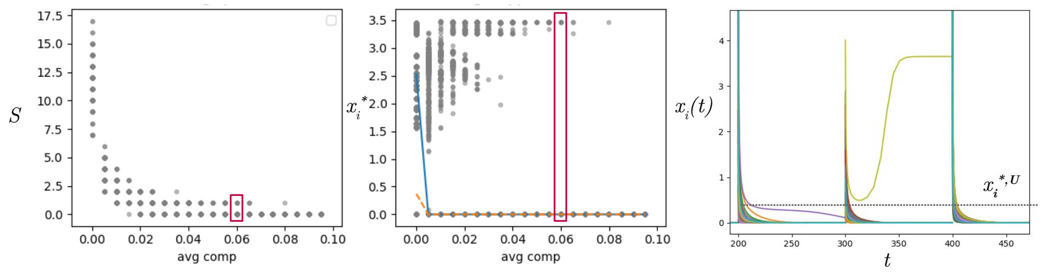


FIG. 8 **Mutual exclusion can coexist with community extinction states.** Strongly competitive systems with monostable species are characterized by mutual exclusion states where one competitor wins. With single-species bistabilities in place, strong competition can also drive all species (here $N = 20$) below their Allee Effect unstable threshold $x_i^{*,U}$, precluding survival of any competitor. The $N$ stable states of mutual exclusion coexist with the global extinction state (red box). Three community trajectories in the right panel show a region where both scenarios are possible, with the Allee Effect sometimes precluding the reinvasion of any of the previously dampened competitors ($\langle B_{ij}\rangle = 0.05$ and three different random initial conditions sorted from $U[0, 5]$).

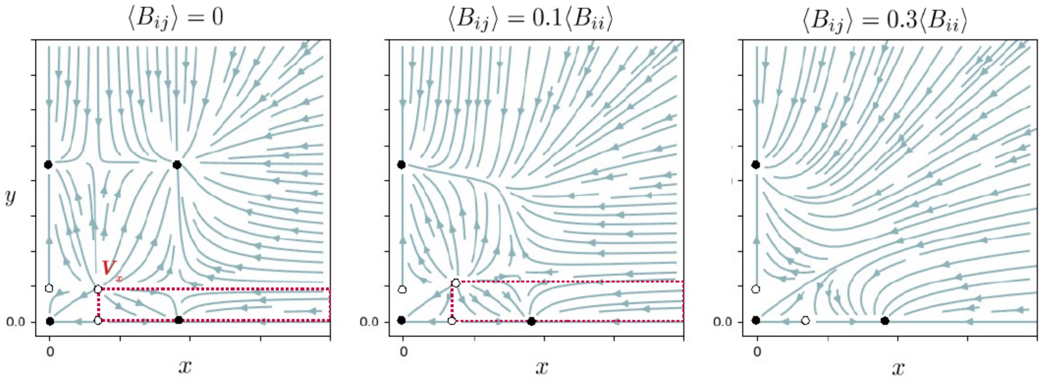


FIG. 9 **Global and exclusion states MSS in competitive 2D systems.** The red area corresponds to the volume of initial conditions $V_x$ where species $x$ would be present without species $y$, hereby dismantling the coexistence state expected from dimensionality reduction techniques. When competition increases, it is not these local states, but the global coexistence state that gets dismantled in favor of local exclusion: one species will be present unless the other overcomes its Allee Effect. When competition is large enough, the system moves towards absolute mutual exclusion: the final survivor is only dictated by the balance in initial conditions, consistent with observations of mutual exclusion in Figs. 3 and 4 of the Main Text.

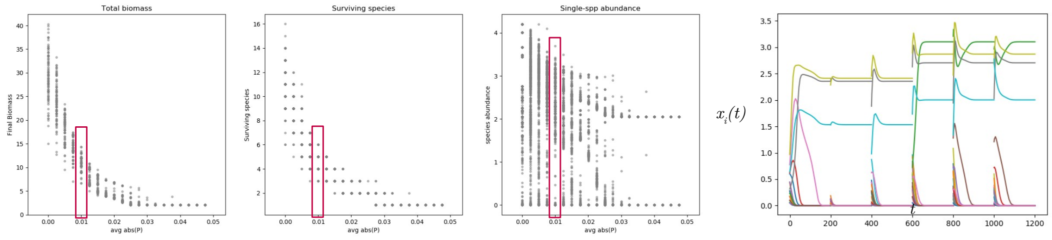


FIG. 10 **Predator-prey interactions in fully-connected networks.** Introducing predator-prey interactions does not necessarily lead to cyclic behavior, but can generate a multistability pattern very close to that of competitive dynamics, with mean absolute predation reducing the diversity and abundance of species in the community (See figure 2B in the main text). On the right, 6 community trajectories, each with a duration of $t = 200$ and with different initial conditions. Here predatory coupling is $\langle |P_{ij}|\rangle = 0.01$ (red box) and the system shows multiple community stable states resulting from single-species bistable dynamics under predation, equivalent to the local multistability phase with reduced diversity.

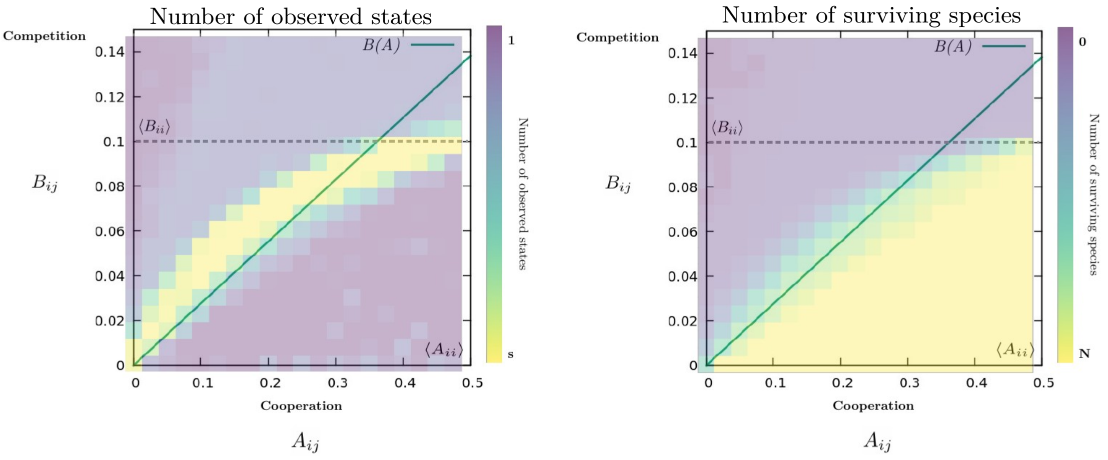


FIG. 11 **Effectively neutral interactions in the homogeneous system.** Analytical simplifications allow us to understand the existence of a domain where positive and negative multilayer interactions cancel out. Species become independently bistable and generate a multiplicity of community states. The underlying map is generated with overall parameter heterogeneity set at $\sigma = 0.01$, and the green line is extracted from the analytical derivation in the text. The background heatmap appears displaced to maintain the origin of the plotted line at $(0, 0)$ while keeping with the original heatmap format.

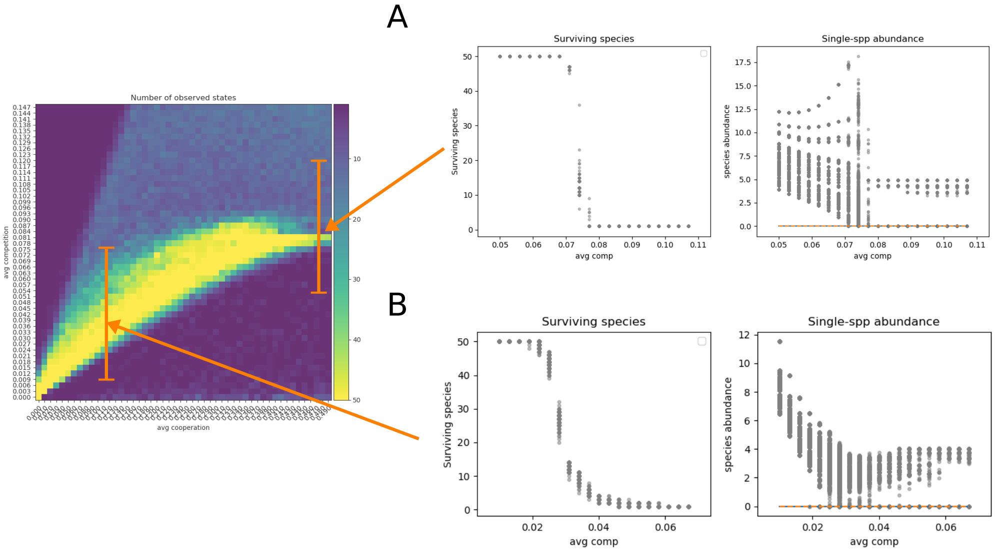


FIG. 12 **Transition from globally bistable to mutual exclusion domains.** Two instances of the number of surviving species (left) and single species abundances (right) for increasing competition. As competition increases, a community with local bistabilities undergoes a transition from a globally bistable phase towards a mutual exclusion phase, where high competition ensures only one species can survive. This transition, however, can take two different flavors. For high mutualism, ensuring that species don't perceive an Allee Effect, the transition is the classical abrupt shift at $B_{ij} = B_{ii}$ with a discontinuity in single species abundances (A), that has been interestingly discussed in (Kessler and Shnerb, 2015) in connection with Hubbell's neutral theory (Hubbell, 2011). Below this competition value, a generalized Lotka-Volterra system would allow for a single state with less than $N$ survivors. But in communities where local bistabilities are in place, this intermediate state becomes pervaded by many possible present-absent combinations and the transition is continuous (B).

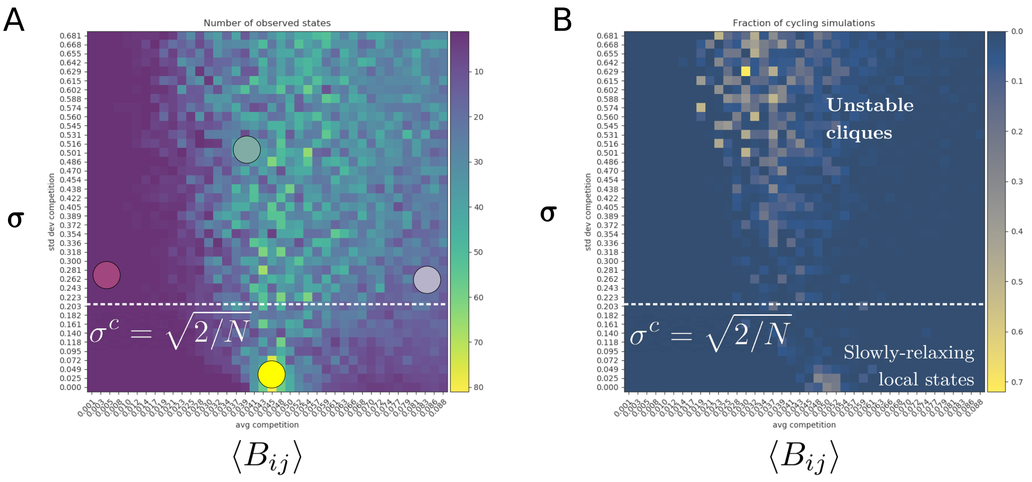


FIG. 13 **Competition-Heterogeneity multistability map for $N = 50$.** Number of stable states (A) and fraction of non-stabilized simulations (B) for the Allee Effect model of the Main Text, when plotting average competition strength against heterogeneity with $\langle A \rangle = 0.2$. Work in generalized Lotka-Volterra systems predicts a single critical transition into the clique domain at a standard deviation of $\sigma^c = \sqrt{2/N}$ for asymmetrical interaction matrices and very large $N \to \infty$. This critical value $\sigma^c = \sqrt{2/N}$ is a good estimate for when cliques and unstable behavior start to emerge for intermediate cooperation-competition ratios. However, when the system is predominantly cooperative or competitive, cliques do not emerge until much higher heterogeneity values. Moreover, the boundaries between MSS phases are not sharp for moderate community sizes of $N = 50$, meaning that different types of multistability can coexist even when the system is found inside the clique domain.

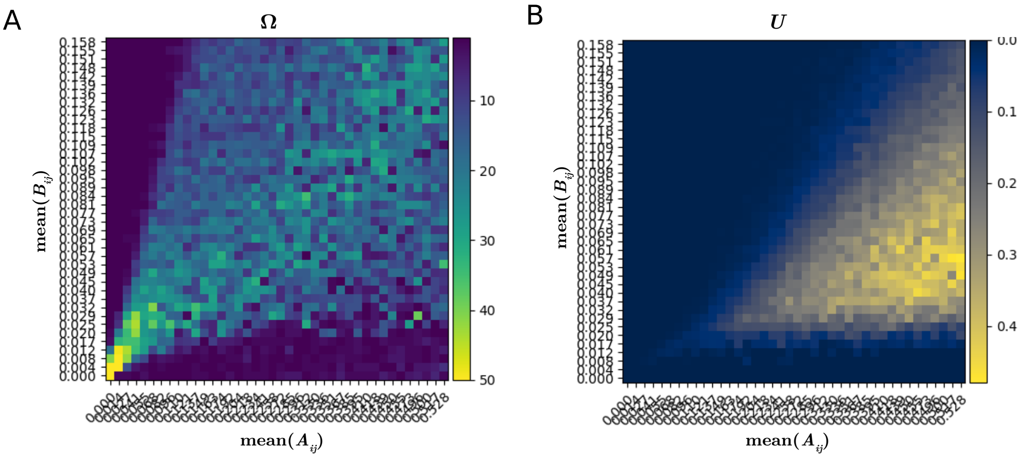


FIG. 14 **Number of stable states and fraction of non-stable simulations under heterogeneous competition.** (Linked to figure 3B in the main text) As opposed to analytical models in the many-species limit, not all cliques are unstable in medium-sized communities with $N = 50$ species. The cliques phase (A, replicated from Figure 3 in the Main Text) harbors a fraction of unstable simulations smaller than 1 (B). Moreover, this fraction appears to increase with interaction strength. As opposed to the rest of the heatmaps in the article, here each square does not represent the results of $s$ simulations for a given system, but rather the average of this over 50 different systems. Although the computational cost of this approach is significant, it enables us to gain a clearer understanding of how unstable scenarios arise in the cliques phase.

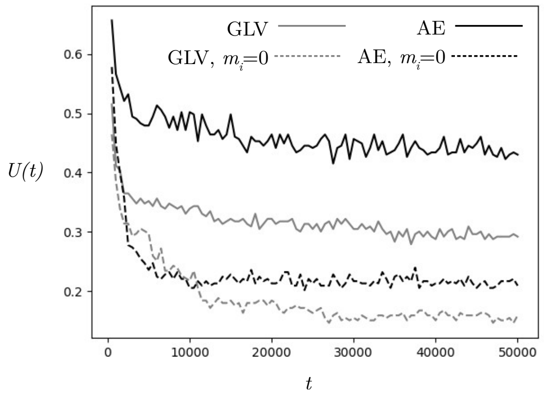


FIG. 15 **Long-term extinction of chaotic regimes under null migration.** As shown in metacommunity models (Roy *et al.*, 2020), chaotic regimes can exist in the form of pinball-like dynamics between unstable cliques if species cannot go extinct for good. Here, we analyze the fraction of unstable simulations along time $U(t)$ for parameters regions within the clique domain for the main Allee Effect model ($\langle A \rangle \in [0.2, 0.5]$, $\langle B \rangle \in [0.03, 0.1]$, $\sigma = 1.5$) and the generalized Lotka-Volterra model (see section III, $\langle A \rangle \in [-1.0, 0.0]$, $\sigma \in [0.05, 0.15]$). Every $t = 200$ timesteps we analyze if a state is persistent or varies (see section I). As expected, in systems without species remigration ($m_i = 0$), chaotic or cyclic regimes go extinct in time, as the system gets stuck in stable configurations.

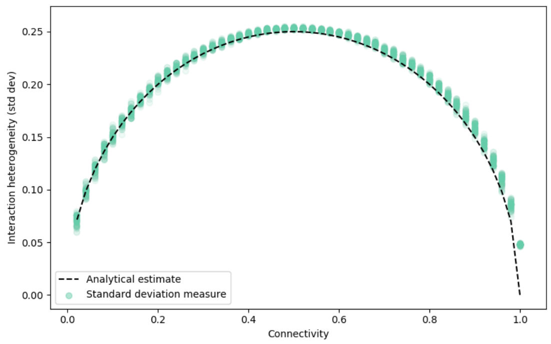


FIG. 16 **Relation between connectivity and matrix standard deviation under homogeneous interaction strengths** Green dots indicate numerically generated matrices following the procedure described in section I.C.4. The dark dashed line is the analytical estimate from section II.F.2. Standard deviation of a non-fully connected matrix with homogeneous values $a$ or 0 can only reach a maximum value of $\sigma = a/2$. Here $a = \langle A_{ii} \rangle = 0.5$.

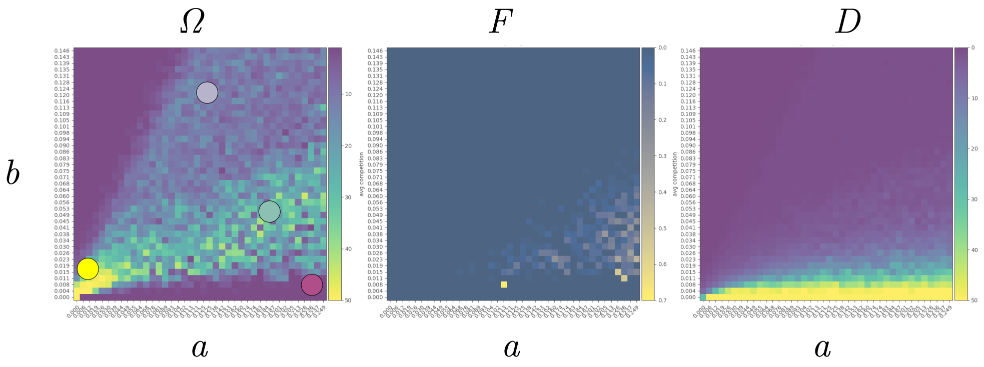


FIG. 17 **Multistability types under homogeneous interaction strengths but $p = 0.5$.** Number of observed states, fluctuations and diversity for an Erdős-Rényi graph with probability of edge existence $p = 0.5$. Reducing network connectivity while maintaining interaction strengths homogeneous gives rise to a qualitatively equivalent phase space to that of Fig. 14: clique-like states emerge together with unstable fluctuations, due to heterogeneity in interactions leading to intransitive competition.

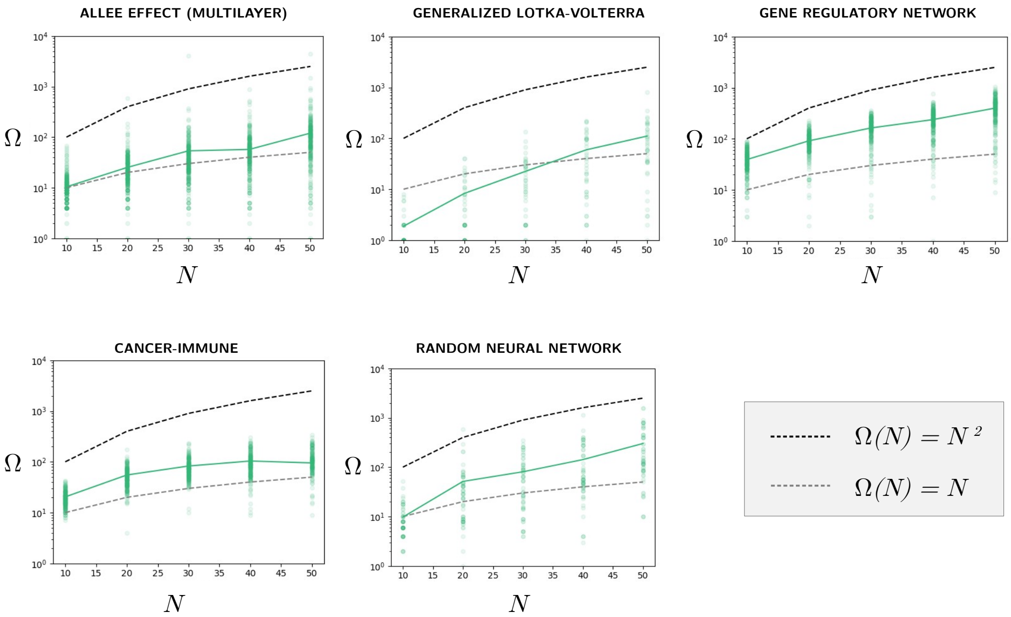


FIG. 18 **Number of observed states $\Omega$ with system size $N$ for the cliques regime.** Scaling of the number of observed states for the model of the main text and those studied in section III, for random $A, B, \sigma$ parameters chosen to fall inside the cliques regime for each model (see section III and figure 22 for location). Despite the complexity of the calculation (see text in section II.G.1), it appears as if the number of cliques scales as $\Omega(N) \sim N^{\theta}$, with $1 < \theta < 2$. However, we can observe how the GLV model (no local multistability regime) supports fewer cliques at low $N$ but has a steeper scaling. This might indicate that local multistabilities somehow contaminate the cliques regime, making it appear as holding a higher number of stable states. Variations in the number of dots for each regime indicate the computational cost of each model, related to the necessary time to stabilization for the chosen parameters. Decays at high $N$, such as that of the Cancer-Immune model, might be related to either true loss of stability, or very long time to stabilization.

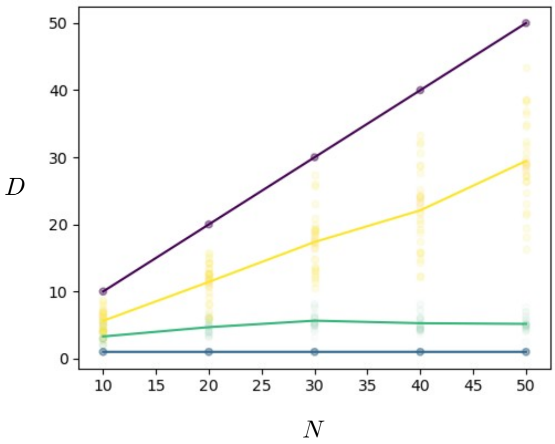


FIG. 19 **Number of surviving species $D$ as system size $N$ increases** Here we plot the diversity of MSS in each category, as the size of the starting species pool increases. As for the scaling figure in the main text (F4B), here we generate random systems, but each with parameters well within each of the regimes of figure 3 in the main text. This allows us to color each system according to what family of MSS it should belong. We see that global states contain all possible species in the community, mutual exclusion states contain one species, whereas local multistability and cliques contain an intermediate value of states. Interestingly, the number of survivors $D$ in local states results directly from how initial conditions are distributed (whether each species falls inside or outside its Allee Effect threshold). On the other hand, the diversity of cliques is much more constrained and does not appear to increase linearly with system size.

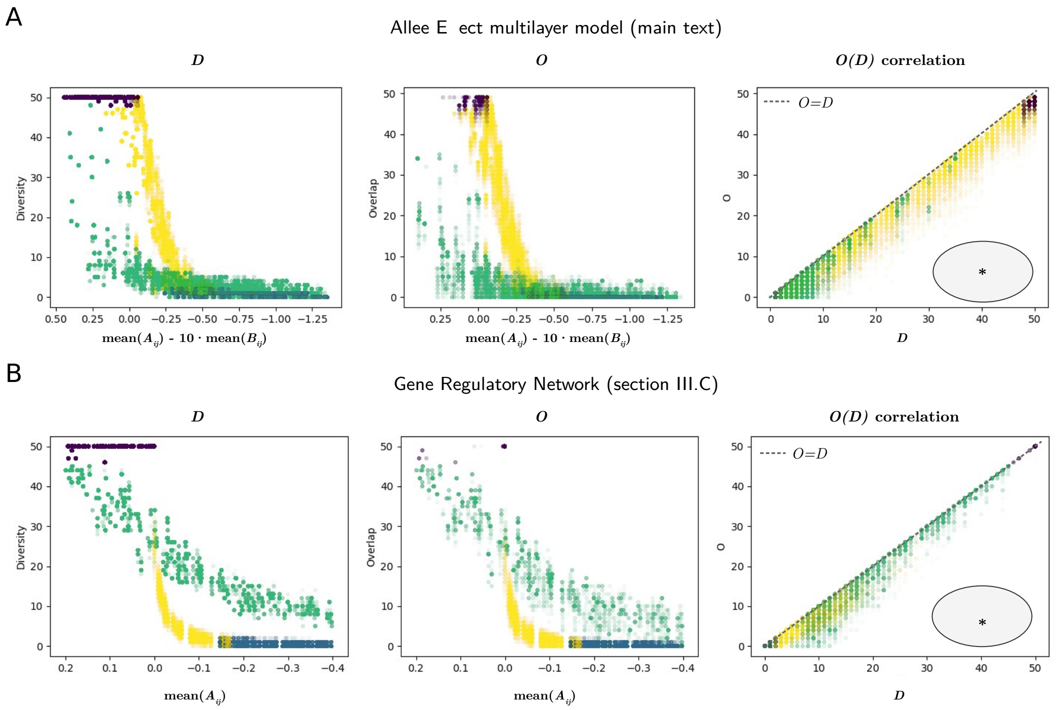


FIG. 20 **Each MSS category has a specific constraint in diversity and species overlap** Here we plot diversity *vs* interaction strength, overlap *vs* interaction strength and overlap *vs* diversity for the Allee Effect model of the Main Text (A) and the Gene Regulatory Network model of section III.C (B). We find that the diversity that MSS can take is rather constrained, meaning that states with large differences in diversity are rare except for the unlikely global-to-extinction shift. This is visualized by the $O(D)$ correlation: the amount of overlap between two states after a transition is very similar to their diversity: most species remain in place. No states deviate strongly from this correlation (gray $*$ area). This area would hold states of high diversity that transition into other states that share very few species (only global bistability could occupy it provided particularly strong perturbations). As global bistability is a rarity of strongly mutualistic systems of bistable species, this further reinforces the notion that catastrophic shifts do not naturally emerge in random high-dimensional models.

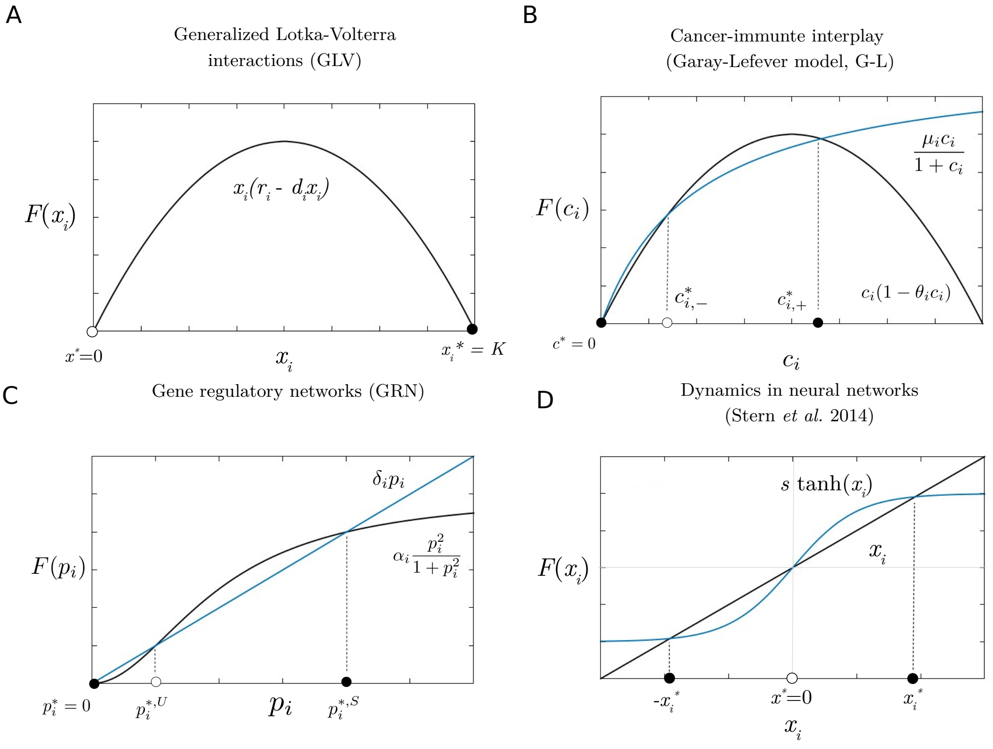


FIG. 21 **One-dimensional dynamics of the additional complex systems models under study** Unit activity (species abundance, cellular abundance, protein concentration and neural cluster activity) for the four studied models, namely (A) Generalized Lotka-Volterra, (B) Garay-Lefever model of the cancer-immune interaction, (C) a gene regulatory network model of activation and inhibition and (D) a firing-rate model of bistable neural clusters. We highlight here the particularity of (A) not showing bistability and hence no global nor local states will emerge in the high-dimensional model, and (D) not showing species extinction and hence no mutual exclusion states will emerge under strong competition.

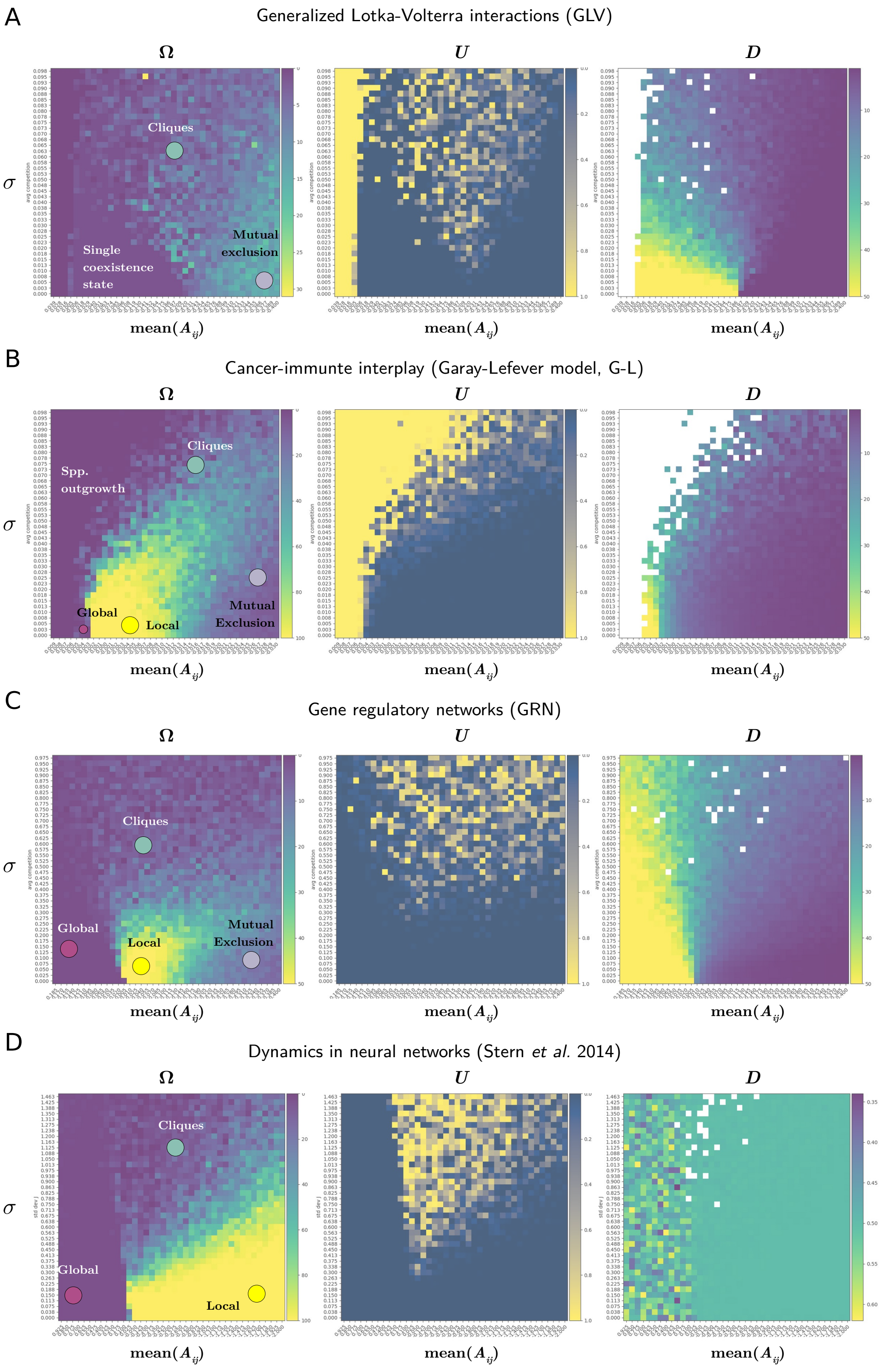


FIG. 22 **Stable states Ω, Fraction of unstable simulations $U = 1 - S$ and diversity $D$ across complex systems models**. Numerical estimate of the multistability phases in (A) GLV interactions, (B) Cancer-immune interplay, (C) GRN and (D) Neural firing-rate model. Even if there are differences in the phase diagram of each high-dimensional dynamical system arising from their different functional responses, the four studied models show multistability domains that hold qualitatively equivalent MSS to those of the Main Text model. Moreover, each dynamical system shows a particularity that is consistent with the proposed MSS taxonomy: no local nor global in the absence of species bistability (A) or no mutual exclusion in the absence of extinction (B).
